# IRF4 haploinsufficiency in a family with Whipple’s disease

**DOI:** 10.1101/197145

**Authors:** Antoine Guérin, Gaspard Kerner, Nico Marr, Janet G. Markle, Florence Fenollar, Natalie Wong, Sabri Boughorbel, Danielle T. Avery, Cindy S. Ma, Salim Bougarn, Matthieu Bouaziz, Vivien Beziat, Erika Della Mina, Tomi Lazarovt, Lisa Worley, Tina Nguyen, Etienne Patin, Caroline Deswarte, Rubén Martinez-Barricarte, Soraya Boucherit, Xavier Ayral, Sophie Edouard, Stéphanie Boisson-Dupuis, Vimel Rattina, Benedetta Bigio, Guillaume Vogt, Frédéric Geissmann, Lluis Quintana-Murci, Damien Chaussabel, Stuart G. Tangye, Didier Raoult, Laurent Abel, Jacinta Bustamante, Jean-Laurent Casanova

**Author notes:** Equal contributions (Gaspard Kerner, Nico Marr). Equal contributions (Janet G. Markle, Florence Fenollar, Natalie Wong, Sabri Boughorbel, Danielle T. Avery, Cindy S. Ma, Salim Bougarn). Equal contributions (Frédéric Geissmann, Lluis Quintana-Murci, Damien Chaussabel, Stuart G. Tangye). Equal contributions (Didier Raoult, Laurent Abel, Jacinta Bustamante, Jean-Laurent Casanova).

## Abstract

The pathogenesis of Whipple’s disease (WD) remains largely unknown, as WD strikes only a very small minority of the individuals infected with *Tropheryma whipplei* (Tw). Asymptomatic carriage of Tw is less rare. We studied a large multiplex French kindred, containing four otherwise healthy WD patients (mean age: 76.7 years) and five healthy carriers of Tw (mean age: 55 years). We used a strategy combining genome-wide linkage analysis and whole-exome sequencing to test the hypothesis that WD is inherited in an autosomal dominant (AD) manner, with age-dependent incomplete penetrance. WD was linked to 12 genomic regions covering 27 megabases in the four patients. These regions contained only one very rare non-synonymous variation: the R98W variant of *IRF4*. The five Tw carriers were heterozygous for R98W. Interferon regulatory factor 4 (IRF4) is a transcription factor with pleiotropic roles in immunity. We showed that R98W was a loss-of-function allele, like only five other exceedingly rare *IRF4* alleles of a total of 39 rare and common non-synonymous alleles tested. Furthermore, heterozygosity for R98W led to a distinctive pattern of transcription in leukocytes following stimulation with BCG or Tw. Finally, we found that *IRF4* had evolved under purifying selection and that R98W was not dominant-negative, suggesting that the IRF4 deficiency in this kindred was due to haploinsufficiency. Overall, haploinsufficiency at the *IRF4* locus selectively underlies WD in this multiplex kindred. This deficiency displays AD inheritance with incomplete penetrance, and chronic carriage probably precedes WD by several decades in Tw-infected heterozygotes.

## Introduction

Whipple’s disease (WD) was first described as an intestinal inflammatory disease by George H. Whipple in 1907 (1). Its infectious origin was suspected in 1961 (2), and the causal microbe, *Tropheryma whipplei* (Tw), a Gram-positive actinomycete, was detected by PCR in 1992 (3), and cultured in 2000 (4). Tw is probably transmitted between humans via the oro-oral or feco-oral routes. WD is a chronic condition with a late onset (mean age at onset: 55 years) (5) affecting multiple organs. The clinical manifestations of classical WD are arthralgia, diarrhea, abdominal pain, and weight loss (6–10). However, about 25% of WD patients display no gastrointestinal or osteoarticular symptoms, instead presenting with cardiac and/or neurological manifestations (8, 11–14). WD is fatal if left untreated, and relapses occur in 2 to 33% of treated cases, even after prolonged appropriate antibiotic treatment (15), (16). WD is rare and has been estimated to affect about one in a million individuals (6, 17, 18). However, about two thousand cases have been reported in at least nine countries worldwide, mostly in North America and Western Europe (11, 19–23). Chronic asymptomatic carriage of Tw is common in the general population, and this bacterium has been detected in feces, saliva, and intestinal mucosae. The prevalence of Tw carriage in the feces has been estimated at 2 to 11% for the general population, but can reach 26% in sewer workers and 37% in relatives of patients and carriers (11, 14, 17, 24–28).

Seroprevalence for specific antibodies against Tw in the general population varies from 50% in France to 70% in Senegal (4, 11, 14, 29). At least 75% of infected individuals clear Tw primary infections, but a minority (<25%) become asymptomatic carriers, a small proportion of whom develop WD (∼0.01%) (30). Tw infection is therefore necessary, but not sufficient, for WD development and it is unclear whether prolonged asymptomatic carriage necessarily precedes WD. The hypothesis that WD results from the emergence of a more pathogenic clonal strain of Tw was not supported by bacterial genotyping (31). WD mostly affects individuals of European origin, but does not seem to be favored by specific environments. WD is typically sporadic, but six multiplex kindreds have been reported, with cases often diagnosed years apart, suggesting a possible genetic component (8, 17, 32). WD patients are not prone to other severe infections (33). Moreover, WD has never been reported in patients with conventional primary immunodeficiencies (PIDs) (34). This situation is reminiscent of other sporadic severe infections, such as herpes simplex virus-1 encephalitis, severe influenza, recurrent rhinovirus infection, and severe varicella zoster disease, which are caused by single-gene inborn errors of immunity in some patients (35–38). We therefore hypothesized that WD might be due to monogenic inborn errors of immunity to Tw, with age-dependent incomplete penetrance.

## Results

### A multiplex kindred with WD

We investigated four related patients diagnosed with WD (P1, P2, P3, and P4) with a mean age at diagnosis of 58 years. They belong to a large non-consanguineous French kindred (Figure 1, A). The proband (P1), a 69-year-old woman, presented with right knee arthritis in 2011, after recurrent episodes of arthritis of the right knee since 1980. Tw was detected in the synovial fluid by PCR and culture, but not in saliva, feces, or small intestine tissue by PCR. Treatment with doxycycline and hydroxychloroquine was effective. At last follow-up, in 2016, P1 was well and Tw PCR on saliva and feces was negative. P2, a second cousin of P1, is a 76-year-old woman with classical WD diagnosed at 37 years of age in 1978 by periodic acid– Schiff (PAS) staining of a small intestine biopsy specimen. She was treated with sulfamethoxazole/trimethoprim. At last follow-up, in 2016, Tw PCR on saliva and feces was positive. P3, the father of P1, is a 92-year-old man with classical WD diagnosed at 62 years of age, in 1987, based on positive PAS staining of a small intestine biopsy specimen. Long-term sulfamethoxazole/trimethoprim treatment led to complete clinical and bacteriological remission. P4, the brother of P2, is a 70-year-old man who consulted in 2015 for arthralgia of the knees and right ulna-carpal joints. PCR and culture did not detect Tw in saliva and feces, but serological tests for Tw were positive. Treatment with methotrexate and steroids was initiated before antibiotics, the effect of which is currently being evaluated. All four patients are otherwise healthy. Saliva and/or feces samples from 18 other members of the family were tested for Tw (Figure 1, A; table S1). Five individuals are chronic carriers (mean age: 55 years) and 13 tested negative (mean age: 38 years). Nine additional relatives could not be tested. The distribution of WD in this kindred was suggestive of an AD trait with incomplete penetrance.

**Figure 1.**
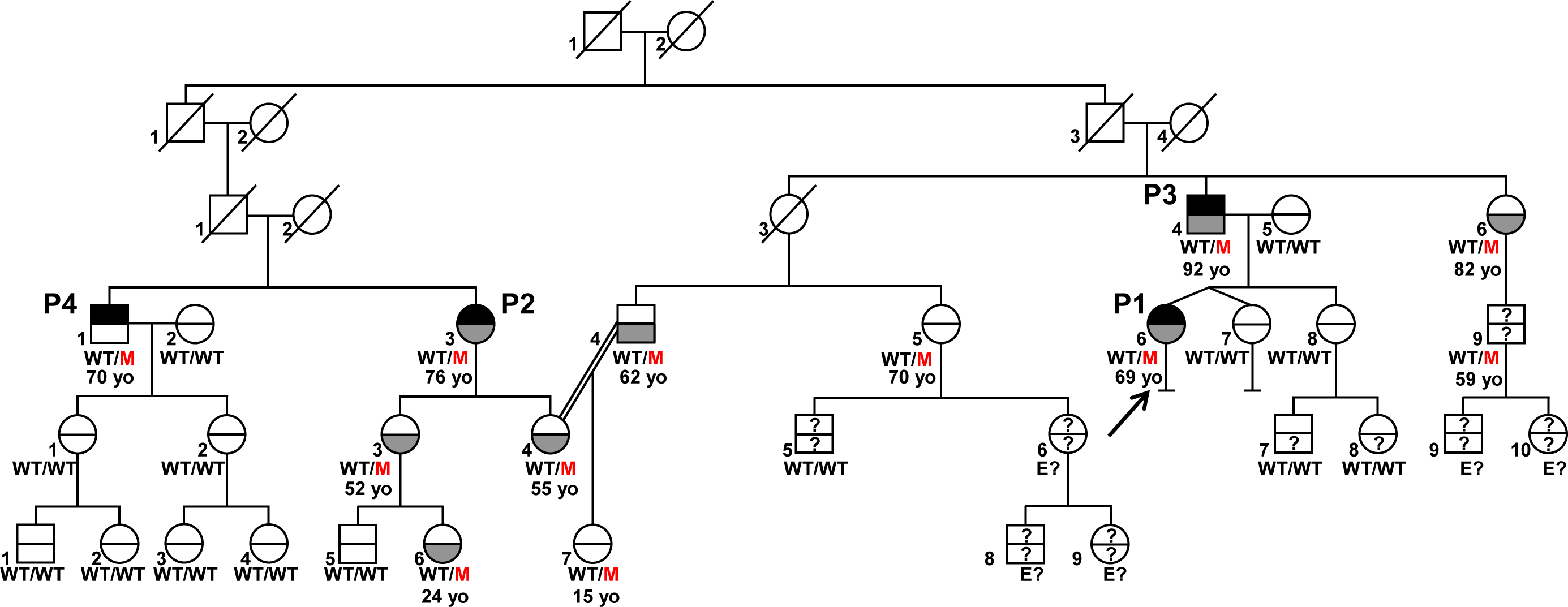
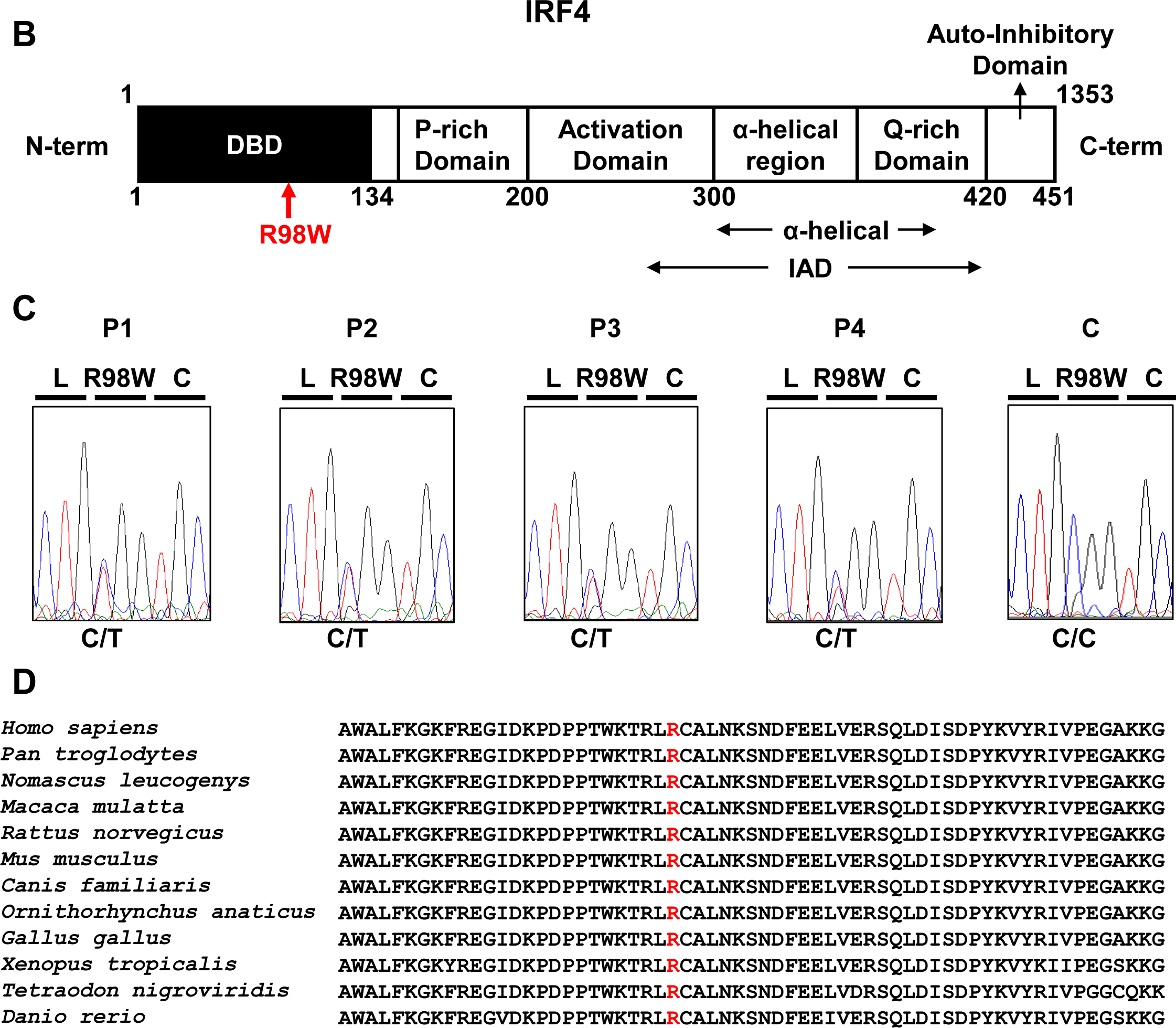
Autosomal dominant IRF4 deficiency. **A.** Pedigree of the kindred with allele segregation. Generations are designated by a Roman numeral (I, II, III, IV, V and VI), and each individual by an Arabic numeral (from left to right). Each symbol is divided into two parts: the upper part indicates the clinical status for WD (black: affected, white: healthy), the lower part indicates whether Tw was identified by PCR (in saliva, blood, feces or joint fluid) or by PAS staining on bowel biopsy specimens (gray: Tw-positive, white: Tw-negative, “?”: not tested). Whipple’s disease patients are indicated as P1, P2, P3, and P4; the proband is indicated with an arrow. Genotype status and age (for *IRF4*-heterozygous individuals) are reported below symbols. Individuals whose genetic status could not be evaluated are indicated by the symbol “E?”. **B.** Schematic representation of the IRF4 protein showing the DNA-binding domain (DBD), P-rich domain, activation domain, α-helical domain, Q-rich domain, IFR association domain (IAD) and auto-inhibitory domain. The R98W substitution is indicated in red. **C.** Electropherogram of *IRF4* genomic DNA sequences from a healthy control (C) and the patients (P1, P2, P3, P4). The R98W IRF4 mutation is caused by the replacement of an arginine with a tryptophan residue in position 98 (exon 3, c.292 C>T). Corresponding amino acids are represented above each electropherogram. **D.** Alignment of the R98W amino acid in the DBD domain of IRF4 in humans and 11 other animal species. R98 is indicated in red.

### A private heterozygous missense *IRF4* variant segregates with WD

We analyzed the familial segregation of WD by genome-wide linkage (GWL), using information from both genome-wide single-nucleotide polymorphism (SNP) microarrays and whole-exome sequencing (WES) (39). Multipoint linkage analysis was performed under an AD model with a very rare disease-causing allele (<10^−5^) and age-dependent incomplete penetrance (see Supplementary Results). Twelve chromosomal regions linked to WD were identified on chromosomes 1 (x3), 2, 3, 6, 7, 8, 10, 11, 12 and 17, with a LOD score close (>1.90) to the maximum expected value (1.95) (Figure S1, A). These regions covered 27.18 Mb and included 263 protein-coding genes. WES data analysis for these 263 genes identified 54 heterozygous non-synonymous coding variants common to all four WD patients (Table S2). Only one, a variant of the *IRF4* gene encoding a transcription factor, member of the *IRF* family (40), and located in a 200 kb linked region on chromosome 6 (Figure S1 A, B), was very rare, and was even found to be private [not found in the GnomAD database, http://gnomad.broadinstitute.org, or in our own WES database (HGID)], whereas all other variants had a frequency >0.001, which is inconsistent with the frequency of WD and our hypothesis of a very rare (<10^−5^) deleterious heterozygous allele. The variant is a c.292 C>T substitution in exon 3 of *IRF4,* replacing the arginine residue in position 98 with a tryptophan residue (R98W) (Figure 1, A, B, C). IRF4 is a transcription factor with an important pleiotropic role in innate and adaptive immunity, at least in mice (41). Mice heterozygous for a null *Irf4* mutation have not been studied, but homozygous null mice have various T- and B-cell abnormalities and are susceptible to both *Leishmania* and lymphocytic choriomeningitis virus (see Supplementary Results) (42–47). We confirmed the *IRF4* R98W mutation by Sanger sequencing genomic DNA from the blood of the four WD patients (Figure 1, C). Thirteen relatives of the WD patients were WT/WT at the *IRF4* locus, and 10 of these relatives (77%) tested negative for Tw carriage. Eight other relatives were heterozygous for the *IRF4* R98W mutation, five of whom (62.5%) were Tw carriers (mean age: 55 years) (Figure 1, A; table S1). Overall, 12 individuals from the kindred, including the four patients, the five chronic carriers of Tw, two non-carriers of Tw and one relative not tested for Tw, were heterozygous for *IRF4* R98W (Figure 1, A; table S1). The familial segregation of the *IRF4* R98W allele was therefore consistent with an AD pattern of WD inheritance with incomplete penetrance. Chronic Tw carriage also followed an AD mode of inheritance.

### R98W is predicted to be loss-of-function, unlike most other IRF4 variants

The R98 residue in the DNA-binding domain (DBD) of IRF4 is highly conserved in the 12 species for which *IRF4* was sequenced (Figure 1, B, D). It has been suggested that this residue is essential for IRF4 DNA-binding activity, because the R98A-C99A double mutant is loss-of-function (48, 49). The R98W mutation is predicted to be damaging by multiple programs (50); it has a CADD score of R98W (26.5), well above the mutation significance cutoff (MSC) of IRF4 (11.125) (Figure S2) (50, 51). The R98W variant was not present in the GnomAD database or our in-house HGID database of more than 4,000 exomes from patients with various infectious diseases. The mutant allele was not found in the sequences for the CEPH-HGDP panel of 1,052 controls from 52 ethnic groups, or in 100 French controls, confirming that this variant was very rare, probably private to this kindred. Therefore, the minor allele frequency (MAF) of this private allele is <4x10^−6^. Moreover, the *IRF4* gene has a gene damage index (GDI) of 2.85, a neutrality index score of 0.15 (52), and a purifying selection *f* parameter of 0.32 (among the <10% of genes in the genome subject to the greatest constraint; Figure S3A), strongly suggesting that *IRF4* has evolved under purifying selection (i.e., strong evolutionary constraints) (53). Biologically disruptive heterozygous mutations of *IRF4* are therefore likely to have clinical effects. We identified 156 other high-confidence heterozygous non-synonymous coding or splice variants of *IRF4* (Table S3) in public (GnomAD: 153 variants, all with MAF<0.009) and HGID (3 variants) databases: 147 were missense (two were also found in the homozygous state), four were frameshift indels, three were in-frame indels, one was a nonsense variant, and one was an essential splice variant. Up to 150 of the 156 variants are predicted to be benign, whereas only six were predicted to be potentially loss-of-function (LOF) according to the GnomAD classification. Comparison of the CADD score and MAF of these *IRF4* variants, showed R98W to have the second highest CADD score of the four variants with a MAF < 4 x10^−6^ (Figure S2). These findings suggest that the private heterozygous *IRF4* variant of this kindred is biochemically deleterious, unlike most other rare (MAF<0.009) non-synonymous variants in the general population, 150 of 156 of which were predicted to be benign (54).

### R98W is loss-of-function, unlike most other IRF4 variants

We first characterized IRF4 R98W production and function *in vitro,* in an overexpression system. We assessed the effect of the *IRF4* R98W mutation on IRF4 levels by transiently expressing WT or mutant R98W in HEK293-T cells. IRF4 R98A-C99A, which is LOF for DNA-binding (49), was included as a negative control. In total cell extracts, mutant IRF4 proteins were more abundant than the WT protein, and had the expected molecular weight (MW) of 51 kDa, as shown by western blotting (Figure 2A). The R98 residue has been shown to be located in a nuclear localization signal, the complete disruption of which results in a loss of IRF4 retention in the nucleus (55). We therefore analyzed the subcellular distribution of IRF4 WT and R98W proteins, in total, cytoplasmic and nuclear extracts from transiently transfected HEK293-T cells. The R98W mutant was more abundant than the WT protein in total cell and cytoplasmic extracts, but these proteins were similarly abundant in nuclear extracts (Figure 2B). We performed luciferase reporter assays to assess the ability of the mutant IRF4 protein to induce transcription from interferon-stimulated response element (*ISRE*) motif-containing promoters. Unlike the WT protein, both R98W and R98A-C99A failed to activate the (*ISRE*)_3_ promoter (Figure 2C). We observed no dominant-negative effect of the IRF4 R98W protein (Figure S4). We assessed the ability of R98W to bind DNA, in an electrophoretic mobility shift assay (EMSA) (Figure 2, D, E). Signal specificity was assessed by analyzing both supershift with an IRF4-specific antibody and by competition with an unlabeled competitor probe. The R98W mutation abolished IRF4 binding to the *ISRE cis* element (Figure 2D), and binding of the IRF4-PU.1 complex to interferon composite elements (EICEs) containing both IRF4 and PU.1 recognition motifs (Figure 2E). The R98W allele of *IRF4* is therefore LOF for both DNA binding and the induction of transcription. Next, we tested 39 of the 156 other *IRF4* variants, including all variants found in GnomAD with a MAF above 4.5x10^−5^ (sixteen variants), or with a CADD score above 30 (ten variants), variants predicted to be LOF (five variants including two with a CADD score above 30; the splice variant could not be tested) and variants in the HGID database (three private variants and seven also present in GnomAD) (Figure S3, B, C). Thirty one variants were normally expressed (two of them with higher molecular weight), the five predicted LOF tested were not detectable (as expected since the antibody epitope is in the IRF4 C terminus), as well as three missense (Figure S5, A, B). When tested for (*ISRE*)_3_ promoter activation, only four variants from public databases (P208Q, R259W, R376C and R376H, MAF between 4.1 x10^−6^ and 7.2x10^−5^) and one from the HGID database (G279-H280 del, private to one family) were LOF (Figure S5, C, D). Our data show that the R98W *IRF4* allele is LOF, like only five very rare other non-synonymous *IRF4* variants out of the 39 variants tested.

**Figure 2.**
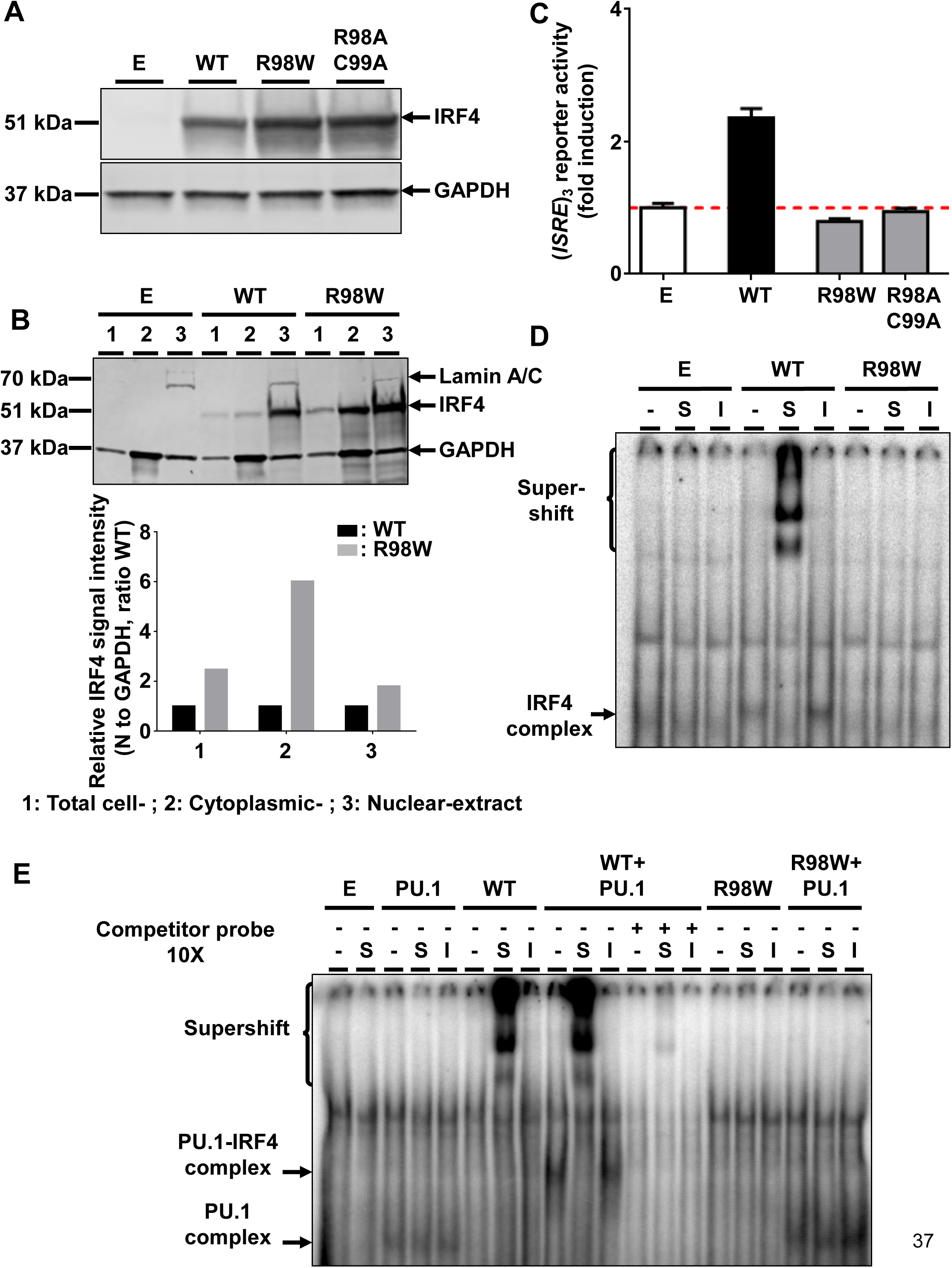
Molecular characterization of the R98W *IRF4* mutation (loss of DNA binding). **A.** HEK293-T cells were transfected with an empty pcDNA3.1 plasmid (E) or with pcDNA3.1 plasmids carrying wild-type (WT) *IRF4*, R98W or R98A-C99A *IRF4* mutant alleles. Total cell extracts were subjected to western blotting; the upper panel shows IRF4 levels and the lower panel shows GAPDH levels, used as a loading control. The results shown are representative of three independent experiments. **B.** (upper panel) HEK293-T cells were transfected with an empty pcDNA3.1 plasmid (E) or with pcDNA3.1 plasmids carrying the wild-type *IRF4* (WT) or R98W *IRF4* mutant alleles. Total cell (1), cytoplasmic (2) and nuclear (3) extracts were subjected to western blotting. Lamin A/C and GAPDH were used as loading controls. (lower panel) IRF4 signal intensity for R98W-transfected cells and WT-transfected cells, in various cell compartments (total, cytoplasmic and nuclear), normalized against the GAPDH signal, as shown by western blotting. The results shown are representative of three independent experiments. **C**. Luciferase activity of HEK293-T cells cotransfected with an (*ISRE*)_3_ reporter plasmid plus the pcDNA3.1 empty vector (E) and a plasmid encoding WT or R98W or R98A/C99A mutants. Results are show, as fold induction of activity relative to E-transfected cells. The red dotted line indicates mean activity for E-transfected cells. The mean and standard error of three experiments are shown. **D**. Electrophoretic mobility shift assay (EMSA) of nuclear extracts of HEK293-T cells transfected with E, WT or R98W plasmids. Extracts were incubated with a ^32^P-labeled *ISRE* probe. Extracts were incubated with specific anti-IRF4 antibody (S) to detect DNA-protein complex supershift, with isotype antibody (I) to demonstrate the specificity of the complex, and with no antibody (-), as a control. The results shown are representative of three independent experiments. **E.** EMSA of nuclear extracts of HEK293-T cells transfected with E, PU.1, WT, R98W, or cotransfected with PU.1 and WT or PU.1 and R98W plasmids. Extracts were incubated with a ^32^P-labeled λB probe (EICE). Extracts were incubated with specific anti-IRF4 antibody (S) to detect DNA-protein complex supershift, with isotype antibody (I) to demonstrate the IRF4 specificity of the complex and with no antibody (-), as a control. Experiments in the presence of excess of non-radioactive probe (cold probe) demonstrated the probe specificity of the complexes. The results shown are representative of three independent experiments.

### AD IRF4 deficiency phenotypes in heterozygous EBV-B cells

We investigated the cellular phenotype of heterozygosity for the R98W allele in EBV-transformed B-cell lines (EBV-B cells) from patients. We performed reverse transcription-quantitative polymerase chain reaction (RT-qPCR) on EBV-B cells from P1, P3, two healthy heterozygous relatives (*IRF4* WT/R98W), four healthy *IRF4-*WT homozygous relatives, 25 patients from our WD cohort (with unknown genetic etiologies) and seven healthy unrelated individuals (*IRF4* WT/WT). Cells from individuals heterozygous for the R98W mutation (patients and healthy carriers) had higher *IRF4* mRNA levels than those from WT homozygous relatives, unrelated WD cohort patients and EBV-B cells from unrelated healthy controls (Figure S6, A). We compared the relative abundances of WT and R98W *IRF4* mRNA in EBV-B cells from heterozygous carriers of the mutation, by performing TA-cloning experiments on P1, P3, one healthy heterozygous relative, one relative homozygous for WT *IRF4*, and two previously tested unrelated controls. In heterozygous carriers of the mutation (patients and healthy relatives) 48.1%-60% of the total *IRF4* mRNA carried the R98W mutation, whereas the rest was WT (Figure S6, B). We evaluated the levels and distribution of IRF4 protein by western blot in EBV-B cells from P1, P2, P3, one healthy heterozygous relative, three healthy homozygous WT relatives and five unrelated healthy individuals. As in transfected HEK293-T cells, IRF4 protein levels were high in both total cell and cytoplasmic extracts of EBV-B cells from heterozygous carriers (Figure S7, A, B). By contrast, IRF4 protein levels in EBV-B cell nuclei were similar in heterozygous mutation carriers and controls (Figure S7, C). As IRF4 is a transcription factor, we then analyzed the steady-state transcriptome of EBV-B cells from three healthy homozygous WT relatives and three WT/R98W heterozygotes (P1, P3, VI.6). We identified 37 protein-coding genes as differentially expressed between subjects heterozygous for *IRF4* and those homozygous WT for *IRF4* (18 upregulated and 19 downregulated; data not shown). We identified no marked pathway enrichment based on these genes. EBV-B cells from *IRF4*-heterozygous individuals had a detectable phenotype, in terms of IRF4 production and function, consistent with AD IRF4 deficiency underlying WD.

### AD IRF4 deficiency phenotypes in heterozygous leukocytes

We assessed IRF4 levels in myeloid and lymphoid cells from healthy controls (see Supplementary Results). IRF4 levels were highest in CD4^+^ T cells, particularly after stimulation with CD2/CD3/CD28 beads (see Supplementary Results). We therefore assessed the IRF4 protein expression profile in CD4^+^ T cells from four controls, P1 and P3, with and without (non-stimulated, NS) stimulation with CD2/CD3/CD28-coated beads. The results were consistent with those for transfected HEK293-T and EBV-B cells, as IRF4 levels in total and cytoplasmic extracts were higher in CD4^+^ T cells from P1 and P3, whereas IRF4 levels in the nucleus were similar in heterozygous mutation carriers and controls (Figure 3, A, B, C). We checked for transcriptomic differences associated with genotype and/or infection, by investigating the transcriptomes of peripheral mononuclear blood cells (PBMCs) from six *IRF4-*heterozygous individuals (three patients, P1-P3; and three healthy relatives, HET1-HET3) and six *IRF4* WT-homozygous individuals (four healthy relatives, WT1-WT4; and two unrelated controls, C1-C2) with and without *in vitro* infection with Tw, or *Mycobacterium bovis*-Bacillus Calmette-Guerin (BCG) for 24 hours. We performed unsupervised hierarchical clustering of the differentially expressed (DE) transcripts (infected versus uninfected) to analyze the overall responsiveness of PBMCs from individual subjects to BCG and Tw infections *in vitro.* Heterozygous individuals clearly clustered separately from homozygous WT individuals (Figure 4, A), revealing a correlation between genotype and response to infection. In homozygous WT subjects, 402 transcripts from 193 unique genes were responsive to BCG infection, and 119 transcripts from 29 unique genes were responsive to Tw infection (Table S4 A, B). Due to the small number of Tw-responsive transcripts linked to unique genes, we were unable to detect any pathway enrichment in this specific condition. However, we identified 24 canonical pathways as enriched after the exposure of PBMCs to BCG. We ranked these pathways according to the difference in mean *z*-score between homozygous WT and heterozygous subjects (Figure 4, B). The top 10 pathways included the interferon signaling network, the Th1 pathway network, the HMGB1 signaling network, the p38 MAPK signaling network, the NF-κB signaling network, the dendritic cell maturation network and the network responsible for producing nitric oxide and reactive species. These pathways were highly ranked mostly due to *IFNG* and *STAT1,* which were strongly downregulated in *IRF4* heterozygotes, particularly in P1, P2 and P3, relative to WT homozygotes. IRF4 is predicted to bind the promoter regions of 47% of the genes identified in the BCG study (91 of 193 genes), including those of *IFNG* and *STAT1*. Subjects heterozygous for *IRF4* also had lower levels of *LTA* expression and lower levels of *IL2RA* expression were observed specifically in patients (Table S4). These data suggest a general impairment of the T-cell response in subjects heterozygous for *IRF4* upon BCG infection *in vitro*. Moreover, the lower levels of *CD80* expression suggest a possible impairment of myeloid and/or antigen-presenting cell function upon BCG infection in patients but not in healthy heterozygous or homozygous WT subjects (Table S4). Peripheral leukocytes from *IRF4*-heterozygous individuals therefore had a phenotype in terms of IRF4 production and function.

**Figure 3.**
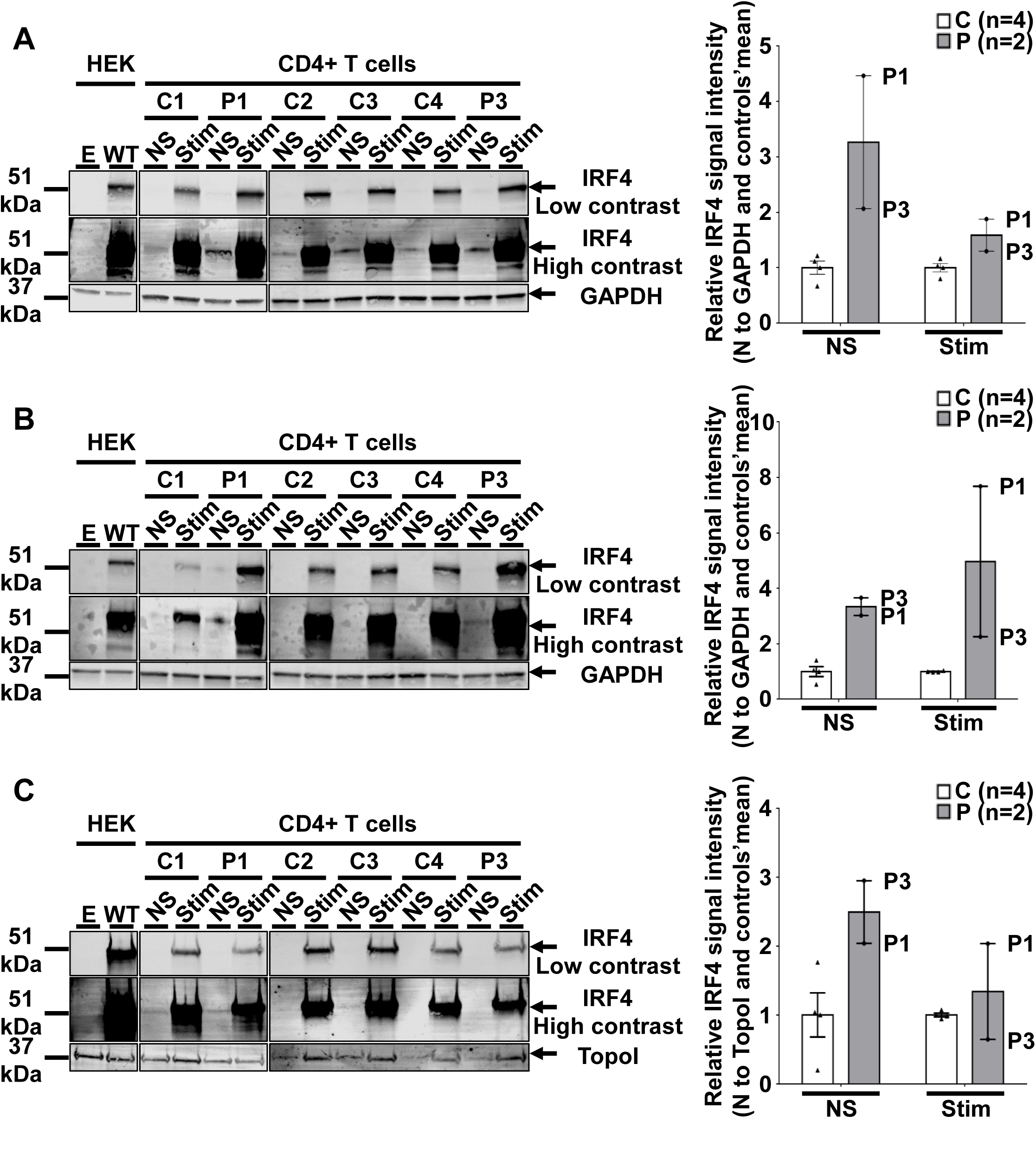
IRF4 protein levels in CD4_+_ T cells. **A.-C.** (Left) Total-cell (A), cytoplasmic (B) and nuclear (C) extracts from CD4^+^ T cells from four unrelated controls (C1 to C4) and two patients (P1 and P3) stimulated with CD2/CD3/CD28-coated beads (Stim) or left unstimulated (NS). Protein extracts from HEK293-T cells transfected with E or WT plasmids were used as controls for the specific band corresponding to IRF4. (Right) Representation of IRF4 signal intensity for each individual relative to the mean signal for unrelated controls (*n*=4) obtained by western blotting (Supp. Figure 8 A-C left) normalized against the GAPDH signal (total, cytoplasmic extracts) or the laminin A/C signal (nuclear extracts).

**Figure 4.**
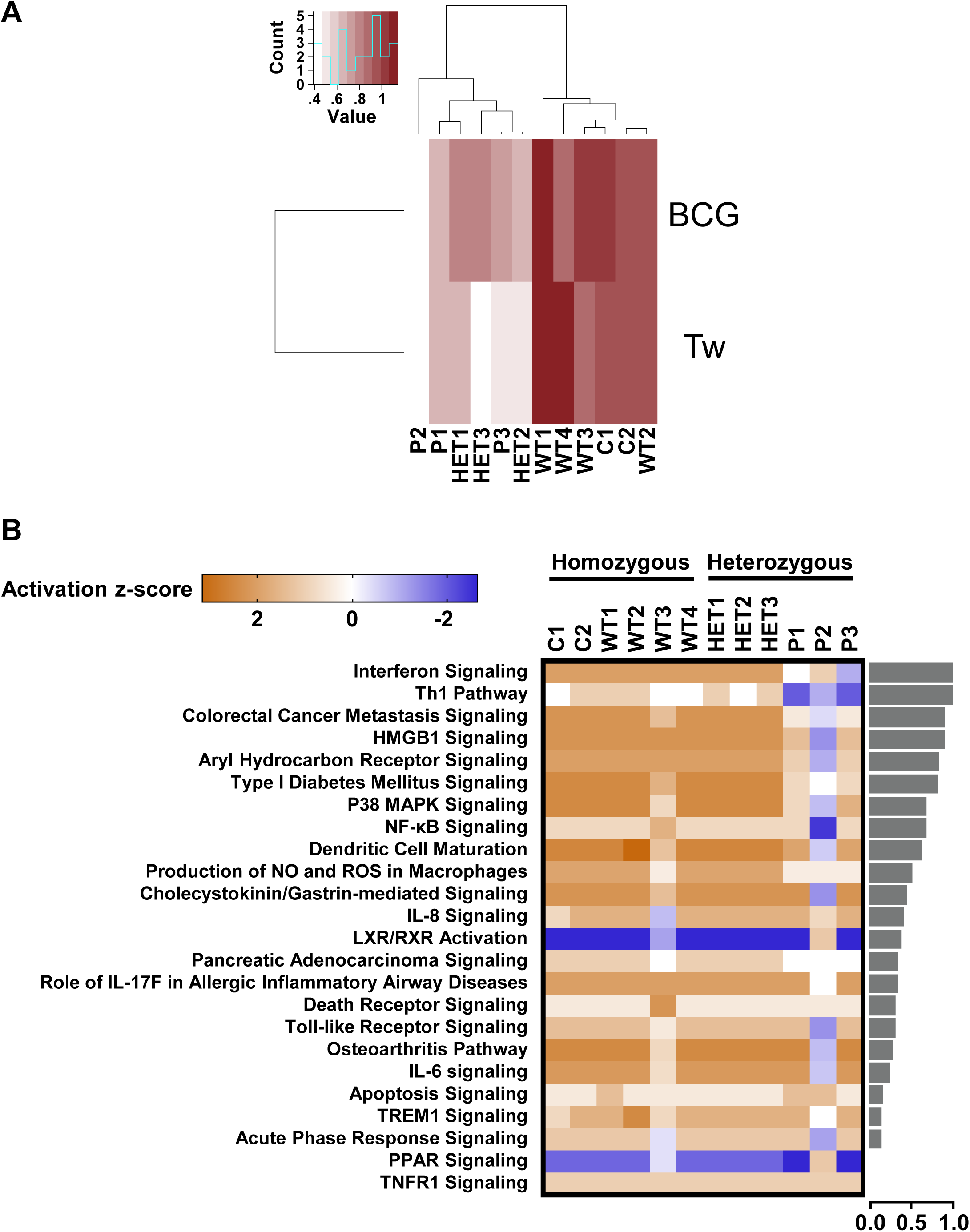
Overall transcriptional responsiveness of PBMCs following *in vitro* exposure to Tw and BCG and pathway activity analysis for genes responsive to BCG exposure. **A.** The overall responsiveness of individual subjects following stimulation with BCG and Tw, relative to non-stimulated conditions (along the horizontal axis) is shown as a heatmap. Subjects were grouped by unsupervised hierarchical clustering. **B.** Enriched canonical pathways were ranked according to differences in mean activation *z*-score between genotypes (homozygous vs. heterozygous). The activation *z*-scores for each individual and pathway are shown as heat maps. Pathways predicted to be activated are depicted in orange, pathways predicted to be inhibited are depicted in blue. A lack of prediction concerning activation is depicted in white. Individuals are presented in columns, pathways in rows. The pathways are ranked from most different between genotypes (at the top of the list) to least different (at the bottom). The differences in mean activation *z*-scores between homozygous and heterozygous individuals for each pathway are depicted as bars to the right of the heat maps (the direction of difference is not shown). The Ingenuity Pathway Analysis (IPA) tool was used to generate a list of the most significant canonical pathways and their respective activation *z*-scores.

## Discussion

WD was initially described as an inflammatory disease (1), but subsequently shown to be infectious (2–4). We provide evidence that WD is also a genetic disorder. We show here that, in a large multiplex kindred, heterozygosity for the private, loss-of-function R98W mutation of *IRF4* underlies an AD form of WD with incomplete penetrance. The causal relationship between *IRF4* genotype and WD was demonstrated as follows. First, the *IRF4* R98W mutation is the only non-synonymous rare variant segregating with WD in this kindred. Second, the mutation was demonstrated experimentally to be loss-of-function, unlike 34 of 39 other non-synonymous *IRF4* variants in the general population, including all eight variants with a MAF greater than 0.0001. Only six of the 156 identified *IRF4* variants were predicted to be LOF, five of which were experimentally tested and shown not to be LOF. Moreover, *IRF4* has evolved under purifying selection, suggesting that deleterious heterozygous variants of this gene entail fitness costs (56–58). Third, EBV-B cell lines heterozygous for *IRF4* R98W have a distinctive phenotype, particularly for IRF4 expression. This mutation also has a strong functional impact on the gene expression of *IRF4* R98W-heterozygous PBMCs stimulated with BCG or Tw. These findings unequivocally show that heterozygosity for the R98W allele of *IRF4* is the genetic etiology of WD in this kindred. Other patients may also develop WD due to inborn errors of immunity. WD is a late-onset infectious disease. This observation therefore extends our model, in which life-threatening infectious diseases striking otherwise healthy individuals during primary infection can result from single-gene inborn errors of immunity (35, 36).

In this kindred with AD IRF4 deficiency, haploinsufficiency was identified as the key mechanism, although IRF4 protein levels in the cytoplasmic compartment were higher in patients with the mutation than in wild-type homozygotes. The protein was not more abundant in the nucleus, where IRF4 exerts its effects on transcription. Moreover, not only is *IRF4* subject to purifying selection, but the R98W mutation is itself LOF, with no detectable dominant-negative effect at cell level. Haploinsufficiency is an increasingly recognized mechanism underlying AD inborn errors of immunity (58, 59). It is commonly due to loss-of-expression alleles, contrasting with the negative dominance typically exerted by expressed proteins, but many mutations are known to cause haploinsufficiency without actually preventing protein production (58–60). Incomplete penetrance is common in conditions resulting from haploinsufficiency. In this kindred, incomplete penetrance may result from a lack of Tw infection, or a lack of WD development in infected individuals. All five chronic carriers of Tw were heterozygous, suggesting that AD IRF4 deficiency also favors the development of chronic Tw carriage, also with incomplete penetrance. The five asymptomatic carriers were 24 to 82 years old, whereas the four patients were 69 to 92 years old. All were heterozygous. The impact of IRF4 R98W may therefore increase with age, initially facilitating chronic carriage in Tw-infected individuals, and subsequently predisposing chronic carriers to WD. Future studies will attempt to define the cellular basis of WD in individuals with *IRF4* mutations. The apparently normal development and function of all myeloid and lymphoid blood subsets studied in patients (see Supplementary Results), and the selective predisposition of these individuals to WD suggest that the disease mechanism is subtle and specifically affects protective immunity to Tw, and that it may act in the gastrointestinal tract.

## Materials and Methods

Informed consent was obtained from all family members, and the study was approved by the national ethics committee.

### Genome-wide analysis

Genome-wide linkage analysis was performed by combining genome-wide array and whole-exome sequencing (WES) data (39). In total, nine family members were genotyped with the Genome-Wide Human SNP Array 6.0. Genotype calling was achieved with the Affymetrix Power Tools Software Package (http://www.affymetrix.com/estore/partners_programs/programs/developer/tools/powertools.affx). SNPs were selected with population-based filters (61), resulting in the use of 905,420 SNPs for linkage analysis. WES was performed as described in the corresponding section, in four family members, P1, P2, P3 and P4. In total, 64,348 WES variants were retained after application of the following filtering criteria: genotype quality (GQ) > 40, minor read ratio (MRR) > 0.3, individual depth (DP) > 20x, retaining only diallelic variants with an existing RS number and a call rate of 100%. Parametric multipoint linkage analysis was performed with the Merlin program (62), using the combined set of 960,267 variants. We assumed an AD mode of inheritance with a frequency of the deleterious allele of 10^−5^ and a penetrance varying with age (0.8 above the age of 65 years, and 0.02 below this threshold). Data for the family and for Europeans from the 1000G project were used to estimate allele frequencies and to define linkage clusters, with an r^2^ threshold of 0.4.

The method used for WES have been described elsewhere (63, 64). Briefly, genomic DNA extracted from the patients’ blood cells was sheared with a Covaris S2 Ultrasonicator (Covaris). An adapter-ligated library was prepared with the Paired-End Sample Prep kit V1(Illumina). Exome capture was performed with the SureSelect Human All Exon kit (71 Mb version - Agilent Technologies). Paired-end sequencing was performed on an Illumina Genome Analyzer IIx (Illumina), generating 72- or 100-base reads. We used a BWA-MEM aligner (65) to align the sequences with the human genome reference sequence (hg19 build). Downstream processing was carried out with the Genome analysis toolkit (GATK) (66) SAMtools (67), and Picard Tools (http://picard.sourceforge.net). Substitution calls were made with a GATK UnifiedGenotyper, whereas indel calls were made with a SomaticIndelDetectorV2. All calls with a read coverage <2x and a Phredscaled SNP quality <20 were filtered out. Single-nucleotide variants (SNV) were filtered on the basis of dbSNP135 (http://www.ncbi.nlm.nih.gov/SNP/) and 1000 Genomes (http:browser.1000genomes.org/index.html) data. All variants were annotated with ANNOVAR (68). All *IRF4* mutations identified by WES were confirmed by Sanger sequencing.

### Tw detection

PCR and serological tests for Tw were performed as previously described (29).

### Cell culture and subpopulation separation

PBMCs were isolated by Ficoll-Hypaque density centrifugation (GE Healthcare) from cytopheresis or whole-blood samples obtained from healthy volunteers and patients, respectively. PBMCs and EBV-B cells were cultured in RPMI medium supplemented with 10% FBS, whereas HEK293-T cells were cultured in DMEM medium supplemented with 10% FBS. Subsets were separated by MACS, using magnetic beads conjugated with the appropriate antibody (Miltenyi Biotec) according to the manufacturer’s protocol.

### Site-directed mutagenesis and transient transfection

The full-length cDNA of *IRF4* and *PU.1* was inserted into the pcDNA_™_3.1D/V5-His-TOPO^®^ vector with the directional TOPO expression kit (Thermo Fisher Scientific). Constructs carrying mutant alleles were generated from this plasmid by mutagenesis with a site-directed mutagenesis kit (QuikChangeII XL; Agilent Technologies), according to the manufacturer’s instructions. HEK293 T cells were transiently transfected with the various constructs, using the Lipofectamine LTX kit (Thermo Fisher Scientific) in accordance with the manufacturer’s instructions.

### Cell lysis and western blotting

Total protein extracts were prepared by mixing cells with lysis buffer (50 mM Tris-HCl pH 7.4, 150 mM NaCl, 0.5% Triton X-100, and 2 mM EDTA supplemented with protease inhibitors (Complete, Roche) and phosphatase inhibitor cocktail (PhoStop, Roche), 0.1 mM dithiothreitol DTT (Life Technologies, California, USA), 10 μg/ml pepstatin A (Sigma, #P4265), 10 μg/ml leupeptin (Sigma, #L2884), 10 μg/ml antipain dihydrochloride (Sigma, #A6191) and incubating for 40 minutes on ice. A two-step extraction was performed to separate the cytoplasmic and nuclear content of the cells; cells were first mixed with a membrane lysis buffer (10 mM Hepes pH 7.9, 10 mM KCl, 0.1 mM EDTA, 0.1 mM EGTA, 0.05 % NP40, 25 mM NaF supplemented with 1 mM PMSF, 1 mM DTT, 10 μg/ml leupeptin, 10 μg/ml aprotinin) and incubated for 30 minutes on ice. The lysate was centrifuged at 10,000 x *g.* The supernatant, corresponding to the cytoplasm-enriched fraction, was collected and the nuclear pellet was mixed with nuclear lysis buffer (20 mM Hepes pH 7.9, 0.4 M NaCl, 1,mM EDTA, 1, mM EGTA, 25% glycerol supplemented with 1 mM PMSF, 1 mM DTT, 10 μg/ml leupeptin, 10 μg/ml aprotinin). Equal amounts of protein, according to a Bradford protein assay (BioRad, Hercules, California, USA), were resolved by SDS-PAGE in a Criterion_™_TGX_™_10% precast gel (Biorad) and transferred to a low-fluorescence PVDF membrane. Membranes were probed with unconjugated antibody: anti-IRF4 (Santa Cruz, M-17) antibody was used at a dilution of 1:1000 and antibodies against GAPDH (Santa Cruz, FL-335), topoisomerase I (Santa Cruz, C-21), and lamin A/C (Santa Cruz, H-110) were used as loading controls. The appropriate HRP-conjugated or infrared dye (IRDye)-conjugated secondary antibodies were incubated with the membrane for the detection of antibody binding by the ChemiDoc MP (Biorad) or Licor Odyssey CLx system (Li-Cor, Lincoln, Nebraska, USA) respectively.

### EMSA

Double-stranded unlabeled oligonucleotides (cold probes) were generated by annealing in TE buffer (pH 7.9) supplemented with 33.3 mM NaCl and 0.67 mM MgCl_2_. The annealing conditions were 100°C for 5 minutes, followed by cooling overnight at room temperature. After centrifugation at 3,000 x *g* centrifugation at 4°C for 30 minutes, the pellet was suspended in water. We labeled 0.1 μg of cold probe in Klenow buffer supplemented with 9.99 mM dNTP without ATP, 10 U Klenow fragment (NEB) and 50 μCi d-ATP-^32^P, at 37°C for 60 minutes. Labeled probes were purified on Illustra MicroSpin G-25 Columns (GE Healthcare Life Sciences) according to the manufacturer’s protocol. We incubated10 μg of nuclear protein lysate for 30 minutes on ice with ^32^P-labeled (a-dATP) *ISRE* probe (5’ – gat cGG GAA AGG GAA ACC GAA ACT GAA-3’) designed on the basis of the *ISG15* promoter or λB probe (5’-gat cGC TCT TTA TTT TCC TTC ACT TTG GTT AC-3’) described by Brass <u>et al</u>. in 1999 (49). For supershift assays, nuclear protein lysates were incubated for 30 minutes on ice with 2 μg of anti-IRF4 (Santa Cruz, M-17) antibody or anti-goat Ig (Santa Cruz) antibody. Protein/oligonucleotide mixtures were then subjected to electrophoresis in 12.5% acrylamide/bis-acrylamide 37.5:1 gels in 0.5% TBE migration buffer for 80 minutes at 200 mA. Gels were dried on Whatman paper at 80°C for 30 minutes and placed in a phosphor-screen cassette for five days. Radioactivity levels were analyzed with the Fluorescent Image Analyzer FLA-3000 system (Fujifilm). For oligonucleotide probes tagged at the 5’ end with a fluorescent IRD700 tag (Metabion), fluorescence was measured with the Licor Odyssey CLx system (Li-Cor, Lincoln, Nebraska, USA) immediately after electrophoresis of the protein/oligonucleotide mixture as described above.

### Luciferase reporter assays

The (*ISRE*)__3__ reporter plasmid, which contains three repeats of the *ISRE* sequence separated by spacers was kindly provided by Prof. Aviva Azriel (Department of Biotechnology and Food Engineering, Technion-Israel Institute of Technology). HEK293 T cells were transiently transfected with the (*ISRE*)_3_ reporter plasmid (100 ng/well for a 96-well plate), the pRL-SV40 vector (40 ng/well) and a *IRF4* WT or mutant pcDNA_™_3.1D/V5-His-TOPO^®^ plasmid (25 ng/well, made up to 50 ng with empty plasmid), with the Lipofectamine LTX kit (Thermo Fisher Scientific), according to the manufacturer’s instructions. Cells were used for luciferase assays 24 h after transfection, with the Dual-Luciferase^®^ 1000 assay system kit (Promega), according to the manufacturer’s protocol. Signal intensity was determined with a Victor^™^ X4 plate reader (Perkin Elmer). Experiments were performed in triplicate and (*ISRE*)_3_ reporter activity is expressed as fold induction relative to cells transfected with the empty vector. Negative dominance was assessed by performing the same protocol with the following modifications: (*ISRE*)_3_ reporter plasmid (100 ng/well for a 96-well plate), pRL-SV40 vector (40 ng/well) WT and mutant plasmids were used to cotransfect cells, with a constant amount of WT plasmid (25 ng/well) but various amounts of mutant plasmid (25 ng/well alone made up to 50 ng with empty plasmid or 25 ng/well, 12.5 ng/well, or 6.25 ng/well made up to 25 ng with empty plasmid) for a total amount of 190 ng/well.

### Microarrays

For the microarray analysis of PBMCs, cells from six *IRF4-*heterozygous individuals (three patients, P1-P3; and three healthy relatives, HET1-HET3) and six *IRF4* WT-homozygous individuals (four healthy relatives, WT1-WT4; and two unrelated controls, C1-C2) were dispensed into a 96-well plate at a density of 200,000 cells/well and were infected *in vitro* with live Tw at a multiplicity of infection (MOI) of 1, or with live BCG (*M. bovis*-BCG, Pasteur substrain) at a MOI of 20, or were left uninfected (mock). Two wells per condition were combined 24 h post-infection for total RNA isolation with the ZR RNA Microprep_™_kit (Zymo Research). For the microarray on EBV-B cells, we used 400,000 cells from three *IRF4-* heterozygous mutation carriers and three WT individuals from the kindred for total RNA isolation with the ZR RNA Microprep_™_kit (Zymo Research). Microarray experiments on both PBMCs and EBV-B cells were performed with the Affymetrix GeneChip Human Transcriptome Array 2.0. Raw expression data were normalized by the robust multi-array average expression (RMA) method implemented in the affy R package (69, 70). Normalized expression data were processed as follows, to select transcripts substantially affected by *in vitro* infection of PBMC samples obtained from the subjects described above with BCG or with Tw. First, fold-changes in expression between mock-infected and BCG-infected or Tw-infected conditions were calculated for each individual separately. For each set of conditions, transcripts were further filtered based on a minimal 1.5-fold change (FC) in expression (up- or downregulation), with a minimum absolute difference in expression of more than 150 relative to unstimulated samples. In a final filtering stage, transcripts satisfying the previous filters in four of the six homozygous individuals with a normal genotype (WT and HET) for each *in vitro* infection condition were retained for downstream analysis and production of the corresponding figures. We analyzed the relative response, by counting the number of probes for which differences were observed between heterozygous individuals and subjects with a WT homozygous genotype. We calculated the number of probes affected by stimulation in samples from control subjects, for the same stimulus. The overall transcriptional responsiveness of individual subjects to both Tw and BCG is depicted as a heatmap, and individual subjects were grouped by unsupervised hierarchical clustering. Responsive transcripts were further analyzed with Ingenuity Pathway Analysis (IPA) Software, Version 28820210 (QIAGEN) (71) for functional interpretation. In brief, the FC values for each individual and treatment were used as input data for the identification of canonical pathway enrichment (*z*-score cut-off was set at 0.1). The activation *z*-score values calculated for the identified pathways were exported from IPA and used to calculate mean values and differences between WT homozygote and heterozygotes and for graphical representation, with Microsoft Excel and GraphPad Prism Version 7.0, respectively. The direction of the difference was not considered further. Negative mean difference values were converted into positive values before the ranking of the canonical pathways according to the difference between the genotypes. The microarray data used in this study have been deposited in the NCBI Gene Expression Omnibus (GEO), under accession number GSE102862.

### *IRF4* qPCR

Total RNA was prepared from the EBV-B cells of patients heterozygous for *IRF4* mutations and WT family members or unrelated individuals (healthy control and a patient with Tw carriage). RNA was prepared from 500,000 cells with the RNeasy Micro kit, according to the manufacturer’s instructions (Qiagen). A mixture of random octamers and oligo dT-16 was used with the MuLV reverse transcriptase (High-Capacity RNA-to-cDNA_™_kit, Thermo Fisher Scientific), to generate cDNA. Quantitative real-time PCR was performed with the TaqMan^®^ Universal PCR Master Mix (Roche), the *IRF4-*specific primer (Hs01056533_m1, Thermo Fisher Scientific) and the endogenous human β-glucuronidase (*GUSB*) as a control (4326320E, Thermo Fisher Scientific). Data were analyzed by the ΔΔCt method, with normalization against *GUSB*.

### *IRF4* TA-cloning

The full-length cDNA generated from the EBV-B cells of heterozygous and WT-homozygous individuals was used for the PCR amplification of exon 3 of *IRF4*. The products obtained were cloned with the TOPO TA cloning kit (pCR2.1®-TOPO^®^ TA vector, Thermo Fisher Scientific), according to the manufacturer’s instructions. They were then used to transform chemically competent bacteria, and 100 clones per individual were Sanger-sequenced with M13 primers (forward and reverse).

### *Ex vivo* naïve and effector/memory CD4_+_ T-cell stimulation

CD4^+^ T cells were isolated as previously described (72). Briefly, cells were labeled with anti-CD4, anti-CD45RA, and anti-CCR7 antibodies, and naïve (defined as CD45RA^+^ CCR7^+^ CD4^+^) T cells or effector/memory T cells (defined as CD45RA^−^CCR7^±^ CD4^+^) were isolated (> 98% purity) with a FACS Aria cell sorter (BD Biosciences). Purified naïve or effector/memory CD4^+^ T cells were cultured with T-cell activation and expansion beads (anti-CD2/CD3/CD28; Miltenyi Biotec) for 5 days; culture supernatants were then used to assess the secretion of IL-2, IL-4, IL-5, IL-6, IL-9, IL-10, IL-13, IL-17A, IL-17F, IFN*γ* and TNF*α* with a cytometric bead array (BD) and the secretion of IL-22, by ELISA.

### *In vitro* differentiation of naïve CD4_+_ T cells

Naïve CD4^+^ T cells (CD45RA^+^CCR7^+^) were isolated (> 98% purity) from healthy controls or patients, with a FACS Aria sorter (BD Biosciences). They were cultured under polarizing conditions, as previously described (72). Briefly, cells were cultured with T-cell activation and expansion beads (anti-CD2/CD3/CD28; Miltenyi Biotec) alone or under Th1 (IL-12 [20 ng/ml; R&D Systems]) or Th17 (TGFβ, IL-1β [20 ng/ml; Peprotech], IL-6 [50 ng/ml; PeproTech], IL-21 [50 ng/ml; PeproTech], IL-23 [20 ng/ml; eBioscience], anti-IL-4 [5 μg/ml], and anti-IFN-γ [5 μg/ml; eBioscience]) polarizing conditions. After five days, culture supernatants were used to assess the secretion of the cytokines indicated, by ELISA (IL-22), or with a cytometric bead array (all other cytokines).

## Acknowledgments

We thank the patients and their families for participating in the study. We thank Yelena Nemirovskaya, Tatiana Kochetkov, Lahouari Amar, Cécile Patissier, Céline Desvallées, Dominick Papandrea, Mark Woollett, and Amy Gall for technical and secretarial assistance and all members of the Laboratory of Human Genetics of Infectious Diseases for helpful discussions. We acknowledge the use of the biological resources of the Imagine Institute DNA biobank (BB-33-00065). A.G. was supported by ANR-IFNPHOX (ANR-13-ISV3-0001-01) and Imagine Institute. The Laboratory of Human Genetics of Infectious Diseases is supported in part by grants from the St. Giles Foundation, The Rockefeller University, Institut National de la Santé et de la Recherche Médicale (INSERM), Paris Descartes University, and the European Research Council (ERC), the Integrative Biology of Emerging Infectious Diseases Laboratory of Excellence (ANR-10-LABX-62-IBEID) and the French National Research Agency (ANR) under the “Investments for the future” program (grant number ANR-10-IAHU-01), ANR-IFNPHOX (ANR-13-ISV3-0001-01, to J.B.), ANR-GENMSMD (ANR-16-CE17-0005-01, to J.B.). Research in the Quintana-Murci laboratory was supported by the Pasteur Institute, the Centre National de la Recherche Scientifique (CNRS), the French Government’s Investissement d’Avenir program, (ANR-10-LABX-62-IBEID), IEIHSEER (ANR-14-CE14-0008-02) and TBPATHGEN (ANR-14-CE14-0007-02), and the European Union’s Seventh Framework Program (FP/2007–2013)/ERC Grant Agreement No. 281297. S.G.T. and C.S.M. are supported by grants and fellowship awarded by the National Health and Medical Research Council of Australia (1113904, 1042925) and the Office of Health and Medical Research of the New South Wales State Government. T.N. and L.W. are supported by Australian Postgraduate Research Awards from the University of NSW.

## Supplementary Results

### Case report

All members of the multiplex kindred studied, the pedigree of which is shown in Figure 1, live in France and are of French descent.

Patient 1 (P1, proband) was born in 1948 and presented arthritis of the right knee in 2011, after recurrent episodes of arthritis of this joint associated with effusion since 1980. *Tropheryma whipplei* (Tw) was detected in synovial fluid by PCR and culture in 2011, but was not detected by PCR in saliva, feces, and small-bowel biopsy specimens. Physical examination revealed a large effusion of the right knee, limiting mobility. The fluid aspirated from this joint contained 4,000 erythrocytes/mm^3^ and 8,800 leukocytes/mm^3^, but no crystals or evidence of microbes. Synovial hypertrophy of the right knee and a narrowing of the right internal femoro-tibial joint were detected on MRI. X ray showed an extension of the right femoro-tibial joint and erosion of the posterior part of the femoro-tibial joint. However, erythrocyte sedimentation rate (ESR) (3 mm/h) and C-reactive protein (CRP) (1.8 mg/l) determinations gave negative results. P1 received methotrexate (15 mg/week) for four months, without remission. Antibiotic treatment with doxycycline (200 mg/day) was then initiated immediately. The arthralgia resolved, but right knee effusion persisted. Hydroxychloroquine was therefore added to the treatment regimen. At last follow-up, in 2016, the patient was well and PCR for Tw was negative for saliva and feces samples.

P2, a second cousin of P1, was born in 1941 and was diagnosed with classical WD and digestive problems in 1978, based on positive periodic acid–Schiff (PAS) staining of a small intestine biopsy specimen. She was treated with sulfamethoxazole/trimethoprim. At last follow-up, in 2016, Tw PCR was positive for the saliva and feces.

P3, the father of P1, was born in 1925 and was diagnosed with classical WD in 1987 on the basis of positive PAS staining of a small intestine biopsy specimen. Clinical manifestations included diarrhea, abdominal pain and weight loss. P3 displayed no extraintestinal manifestations. He was successfully treated with sulfamethoxazole/trimethoprim, with complete clinical and bacteriological remission.

P4, the brother of P2, was born in 1947 and sought medical advice in 2015 for arthralgia affecting the knees and right ulna-carpal joints. The other joints were unaffected. A culture of the joint fluid was negative for bacteria, but Tw was not sought. Tw was not detected in the saliva and feces by PCR or culture, but serological tests for Tw were positive. P4 had no rheumatoid factor, anti-cyclic citrullinated peptide antibodies (anti-CCP), or anti-nuclear antibodies. The fluid aspirated from the right knee contained 4,800 erythrocytes/mm^3^ and 10,900 leukocytes/mm^3^ (91% neutrophils and 9% lymphocytes) without crystals. Synovial fluid culture was negative for bacteria, but Tw was not sought. Blood tests revealed an ESR of 30 mm/h and a CRP concentration of 50 mg/l, with no rheumatoid factor, anti-cyclic citrullinated peptide antibodies (anti-CCP) or anti-nuclear antibodies. An X ray revealed a narrowing of the joint space in the knees and vertebral hyperostosis were visible. The joints of the hands were unaffected. The patient was treated with anti-inflammatory drugs, without success. Treatment with methotrexate and steroids was introduced, followed by antibiotics, the effect of which is currently being evaluated.

Saliva and/or feces samples from 18 other members of the family were checked for the presence of Tw, by a PCR specifically targeting *T. whipplei,* as previously described (Figure 1A) (73). Five individuals were found to be chronic carriers (mean age: 55 years) and 13 were not (mean age: 38 years). Testing was not possible for nine other relatives. The overall distribution of WD in this kindred was suggestive of an AD trait with incomplete penetrance.

### Production and function of IRF4 in the patients’ peripheral leukocytes

We investigated IRF4 levels by western blotting on total cell lysates from peripheral blood mononuclear cells (PBMCs) and their subpopulations isolated from six healthy controls. We showed that IRF4 was produced in large amounts in total B lymphocytes (CD19^+^), but also naïve and memory B lymphocytes (CD19^+^CD27^−^ and CD19^+^CD27^+^, respectively; Figure S8). IRF4 was less strongly expressed in CD3^+^ T lymphocytes and was not detectable in CD14^+^ monocytes or CD56^+^ natural killer (NK) cells. IRF4 levels were low in naïve CD3^+^ T cells and CD4^+^ T cells, but not detectable in naïve CD8^+^ T cells in the basal state, relative to controls cells (western blot, data not shown). We showed, in CD3^+^ T cells and CD4^+^ T cells, that phorbol myristate acetate (PMA)-ionomycin stimulation induced the production of even larger amounts of IRF4 expression following activation with CD2/CD3/CD28-coated beads (data not shown). We therefore assessed the pattern of IRF4 protein production in CD4^+^ T cells from four controls, P1 and P3, either non-stimulated (NS) or stimulated with CD2/CD3/CD28-coated beads. Our results are consistent with our findings for HEK293 T cells, as IRF4 production by P1 and P3 CD4^+^ T cells was stronger than that by CD4^+^ T cells from unrelated controls, in total cell lysates (4.5 times higher for P1 and 2.1 times higher for P3 in NS conditions; 1.9 times higher for P1 and 1.3 times higher for P3 under stimulation), in the cytoplasmic compartment (3 times higher for P1 and 3.7 times higher for P3 in NS conditions; 7.7 times higher for P1 and 2.3 times higher for P3 under stimulation) and in the nuclear compartment in NS conditions (2 times higher for P1 and 3 times higher for P3) but not under stimulation (2 times higher for P1 and 0.6 times higher for P3) (Figure 3, A, B, C). Analyses of the overall pattern of IRF4 expression in the cells of controls and patients showed that the patients’ CD4^+^ T lymphocytes had higher total IRF4 levels and that a higher proportion of IRF4 was located in the cytoplasm than in controls, as observed for EBV-B cells and with the overexpression system.

We characterized the transcriptomic differences linked to genotype and/or to infection, by performing microarray studies on PBMCs (at early time points or in the absence of stimulation for 24 h (NS)) from six *IRF4-*heterozygous individuals (three patients, P1-P3; and three healthy relatives, HET1-HET3) and six individuals homozygous for the WT *IRF4* allele (four healthy relatives, WT1-WT4; and two unrelated controls, C1-C2). A comparison of NS samples from the *IRF4-*heterozygous and *IRF4* WT-homozygous groups showed that differentially regulated transcripts could be detected only at an early time point (49 genes upregulated and 16 downregulated in the heterozygous group relative to the WT homozygotes), but not after 24 h. It was not, therefore, possible to distinguish genes differentially regulated due to the *IRF4* mutation from those differentially regulated as a result of a systemic immune response in the patients.

### Expression and function of IRF4 in the patients’ myeloid cells

An analysis of the three main subsets of DCs present in human blood —CD303^+^ pDCs (CD123^+^ CD303^+^ HLA-DR^+^ Lin^−^ CD14^−^ CD16^−^), CD1c^+^ cDCs (CD1c^+^ CD11c^+^ HLA-DR^+^ Lin^−^ CD14^−^ CD16^−^) and CD141^+^ cDCs (CD141^+^ CD11c^+^ HLA-DR^+^ Lin^−^ CD14^−^ CD16^−^) — showed that the frequencies of these subsets in P1 were similar to those in controls (Figure S9). We assessed IRF4 expression levels and the effect of IRF4 deficiency in myeloid cells, by first studying monocyte-derived dendritic cells (MDDCs) from controls, in which IRF4 was produced after stimulation with LPS but not after stimulation with IFN-*γ.* IRF4 was not produced by immature MDDC (data not shown). We also generated monocyte-derived macrophages (MDMs) for controls and patients, using different conditions of differentiation and activation to obtain M1-like and M2-like MDMs. We found that IRF4 was present in similar amounts in MDMs from patients and controls, regardless of the differentiation or activation conditions used (Figure S10, A and B). FACS analysis of MDMs with common and specific markers (for M1-like and M2-like MDMs) showed that CD11b, CD86, CD206, CD209 and HLA-DR expression was similar for MDMs from patients and MDMs from controls (Figure S10, C and D). Thus, all the myeloid subsets studied developed and functioned normally in the patient, suggesting a subtle disease mechanism.

### Lymphoid immunological phenotype

We then assessed the potential impact of heterozygosity for the R98W mutation of *IRF4* on the development and function of lymphoid subsets. In mice, Irf4 plays a major role in the generation, differentiation and functions of various immune cells, including T and B lymphocytes (74, 75). We analyzed peripheral leukocytes from the three patients (P1, P2, P3) and from the P1’s healthy dizygotic twin sister (WT). All subjects had normal numbers and percentages of T, B, and NK cells for age (table S5). We also performed deeper B-cell immunophenotyping by flow cytometry. The frequency of memory B cells within the total B-cell population was normal, as was the frequency of class-switched B cells in the memory compartment in cells from the patients (P1, P2 and P3), as shown by comparisons with healthy controls (Figure S11, A). We assessed *ex vivo* cytokine production by differentiated memory CD4^+^ Th cells. Cells from the patients (P2 and P3) produced more IL-9 and less IL-17A and IL-17F than cells from controls (Figure. S11, B). We found no significant difference in the production of IL-2, IL-4, IL-5, IL-10, IL-13, IL-22, IFN-γ and tumor necrosis factor-alpha (TNF-α) between patients and controls (Figure S11, B). Analyses of CD4^+^ T-cell differentiation *in vitro* showed that, contrary to the *ex vivo* data obtained for memory CD4^+^ T cells, naïve CD4^+^ T cells from the patients were able to produce relatively normal amounts of IL-17-A/F (Figure S11, C). The *in vitro* production of IFN-γ, IL-10 and IL-21 by naïve CD4^+^ T cells from patients was also unimpaired (Figure S11, C). Furthermore, studies of the *in vitro* function of naïve and total memory B cells upon costimulation with CD40 ligand (CD40L) and IL-21 showed that cells from patients (P2 and P3) produced similar amounts of IgA, IgG and IgM to cells from healthy controls (data not shown). Thus, overall, in the experimental conditions tested, the *IRF4* R98W mutation has no global effect on the development or function of adaptive immune system lymphocytes, suggesting that haploinsufficiency for IRF4 does not underlie WD through adaptive immune responses.

### Additional information concerning the mouse model

Irf4-deficient homozygous mice have lymphoid defects relating to differentiation (Th1, B cells), proliferation (T and B cells), Ig levels (IgG and IgM) and activity (abolition of IL-4 and IFN-*γ* production by CD4^+^ T cells after infection with *Leishmania major*). They also display splenomegaly, adenomegaly, lower counts of plasmacytoid dendritic cells (pDCs) in the spleen and thymus, of the abolition of plasma cells in the spleen and abnormal class-switch recombination. Moreover, homozygous mice have no germinal center in the spleen and lymph nodes and are more susceptible to *Leishmania* and lymphocytic choriomeningitis virus (42–47). Mice heterozygous for an *Irf4*-null mutation have not been studied.

## Supplementary Figures legends

**Supplementary Figure 1.**
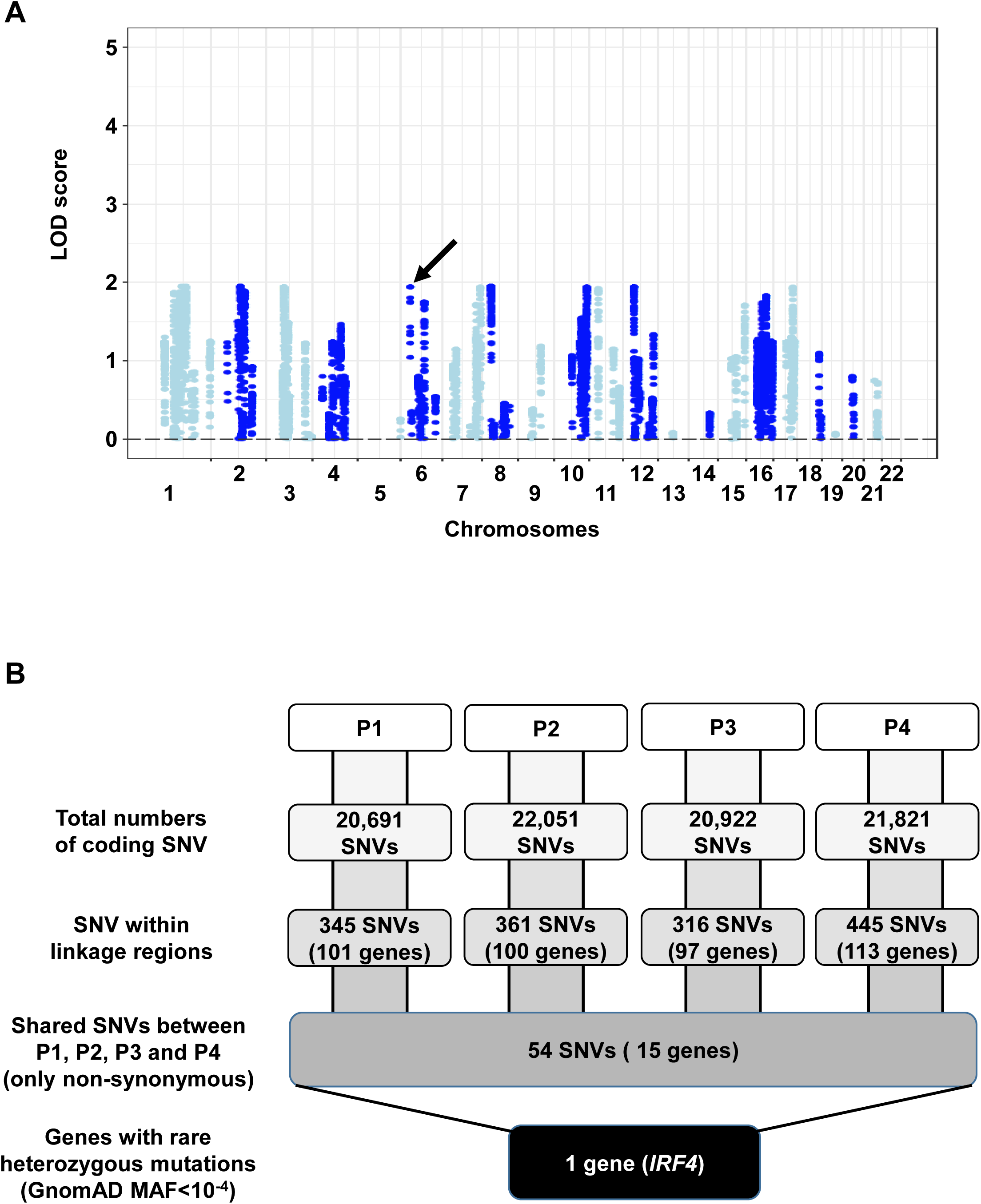
Genome-wide linkage and whole-exome sequencing analyses. **A**. Genome wide linkage analysis was performed by combining Genome-wide array and whole-exome sequencing (WES) data assuming an autosomal dominant (AD) mode of inheritance. LOD (logarithm of odds) scores are shown for the four patients considered together. The maximum expected LOD score is 1.95, based on an AD model with incomplete penetrance. *IRF4* is located within a linkage region (LOD=1.94) on chromosome 6 (indicated by a black arrow). **B.** A refined analysis of WES data identified *IRF4* as the only protein-coding gene carrying a rare heterozygous mutation common to P1, P2, P3 and P4 within the linkage regions.

**Supplementary Figure 2.**
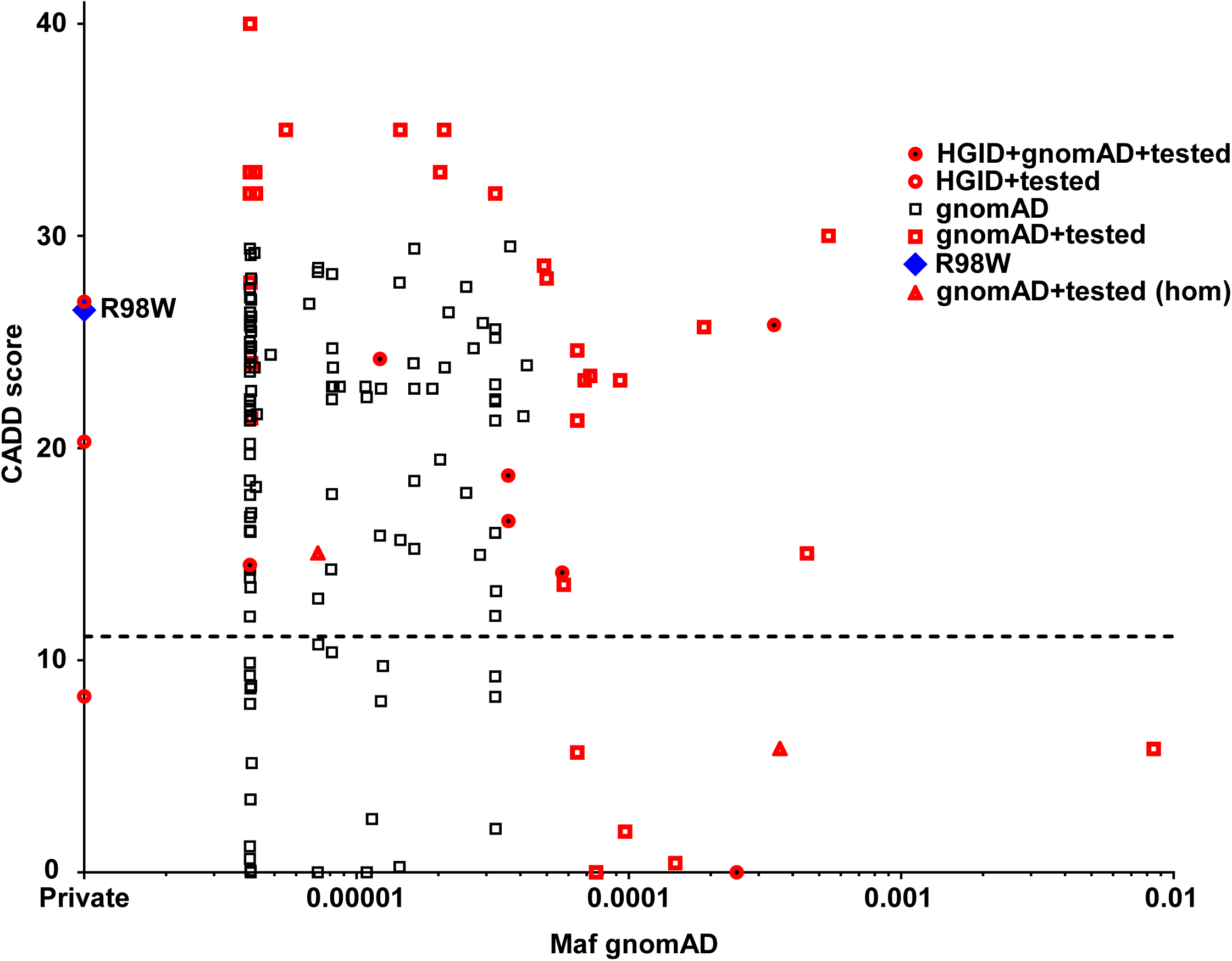
Analysis *in silico* of *IRF4* variants. **A.** Minor allele frequency and combined annotation–dependent depletion score (CADD) of all coding variants reported in public database (GnomAD) (http://gnomad.broadinstitute.org) and in-house (HGID) databases. The dotted line corresponds to the mutation significance cutoff (MSC) with 95% confidence interval. The variants studied *in vitro* here are shown in red, in bold typeface. The R98W variant is represented by a blue square.

**Supplementary Figure 3.**
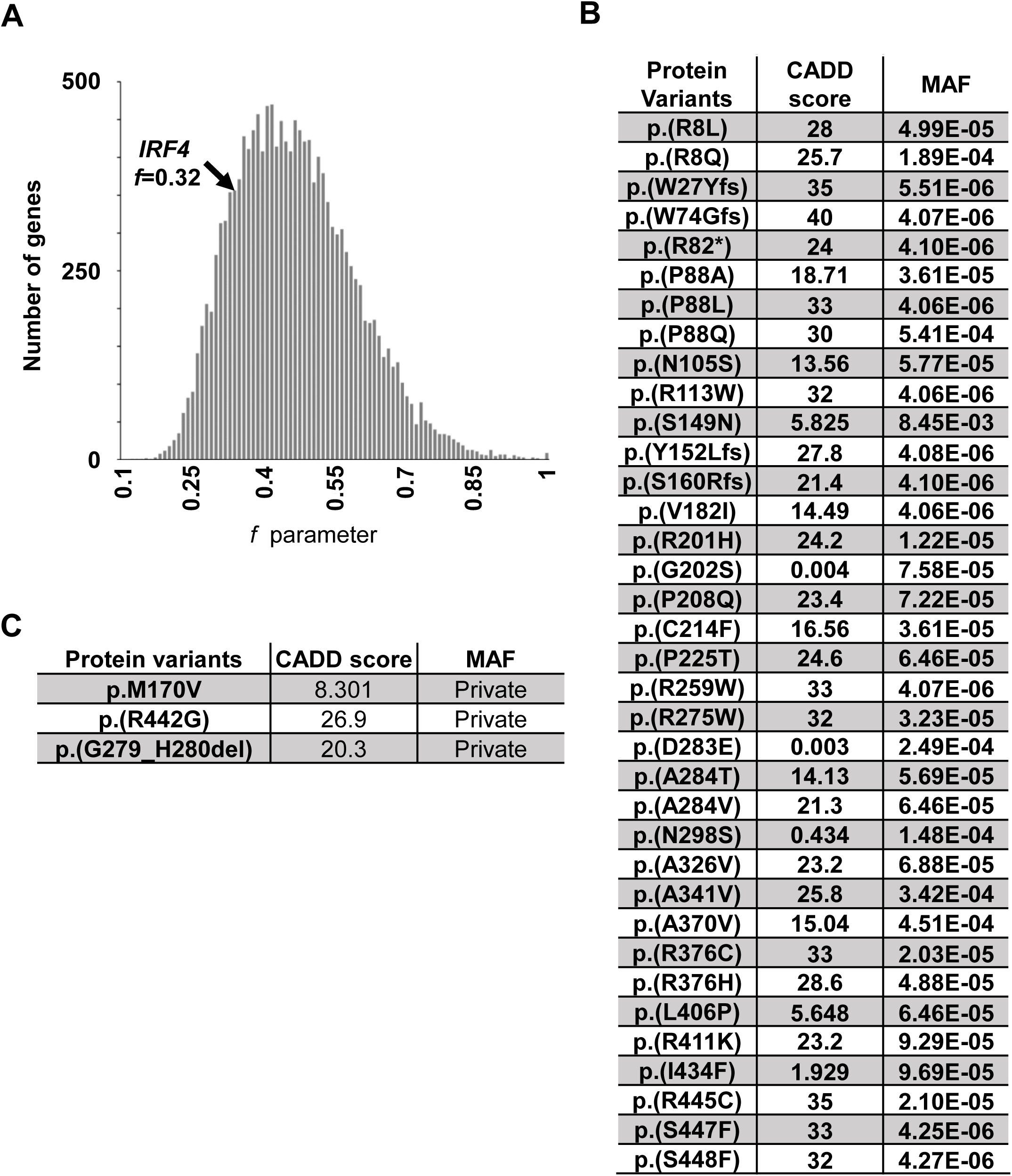
List of variants and strength of purifying selection on *IRF4*. **A.** Genome-wide distribution of the strength of purifying selection, estimated by the *f* parameter (53), acting on 14,993 human genes. *IRF4* is at the 9.4^th^ percentile of the distribution, indicating that it is more constrained than most human genes. **B**. List of *IRF4* missense and in-frame deletion variants reported in a public database (GnomAD) and studied *in vitro* here. **C.** List of *IRF4* missense and in-frame deletion variants found in the HGID database and studied *in vitro* here.

**Supplementary Figure 4.**
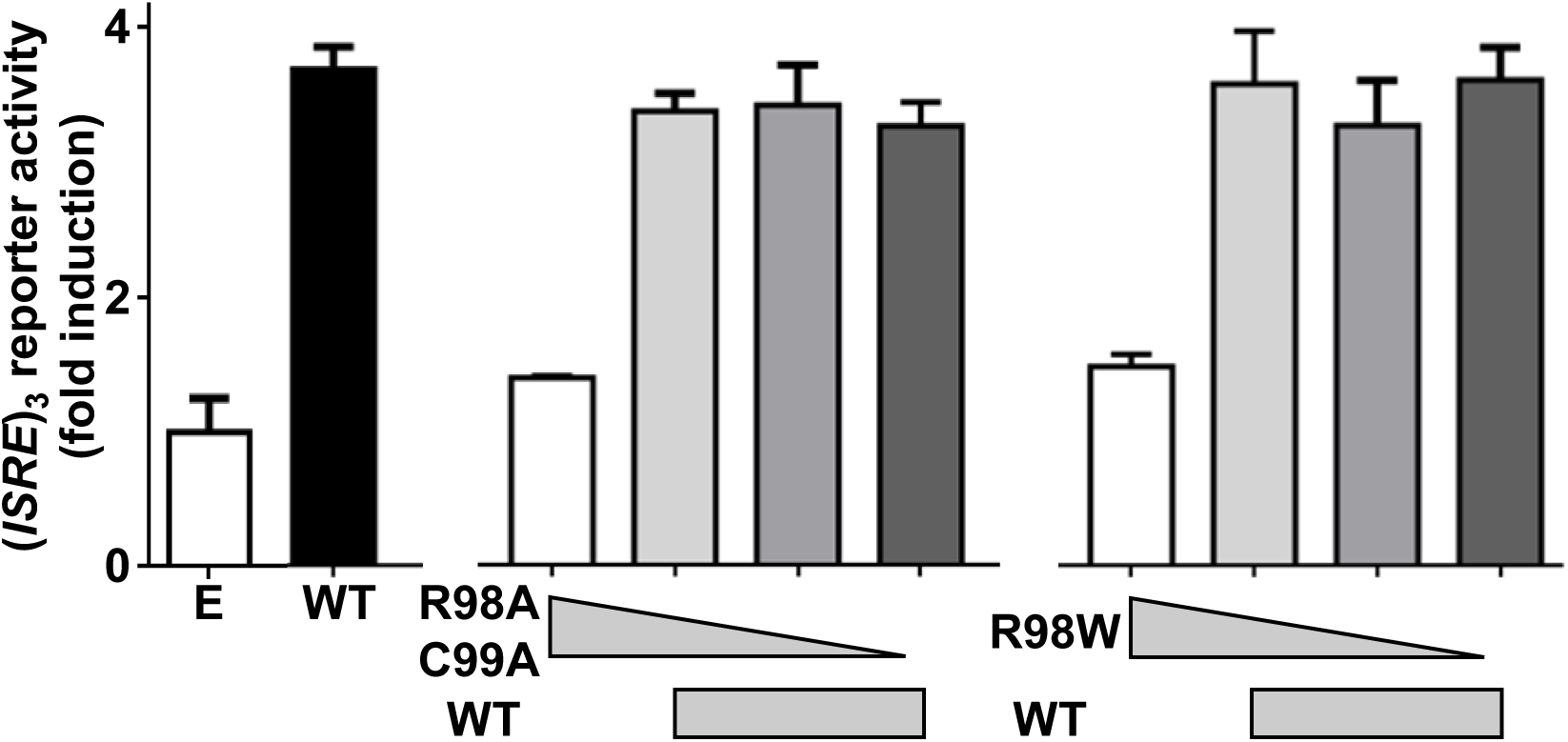
Functional activity of IRF4. Luciferase activity of HEK293-T cells cotransfected with an (*ISRE*)_3_ reporter plasmid plus E and WT or various amounts of R98W or R98A/C99A plasmids. Results are showed as fold induction of activity relative to E-transfected cells. The results shown are the mean ± S.D. of three independent experiments.

**Supplementary Figure 5.**
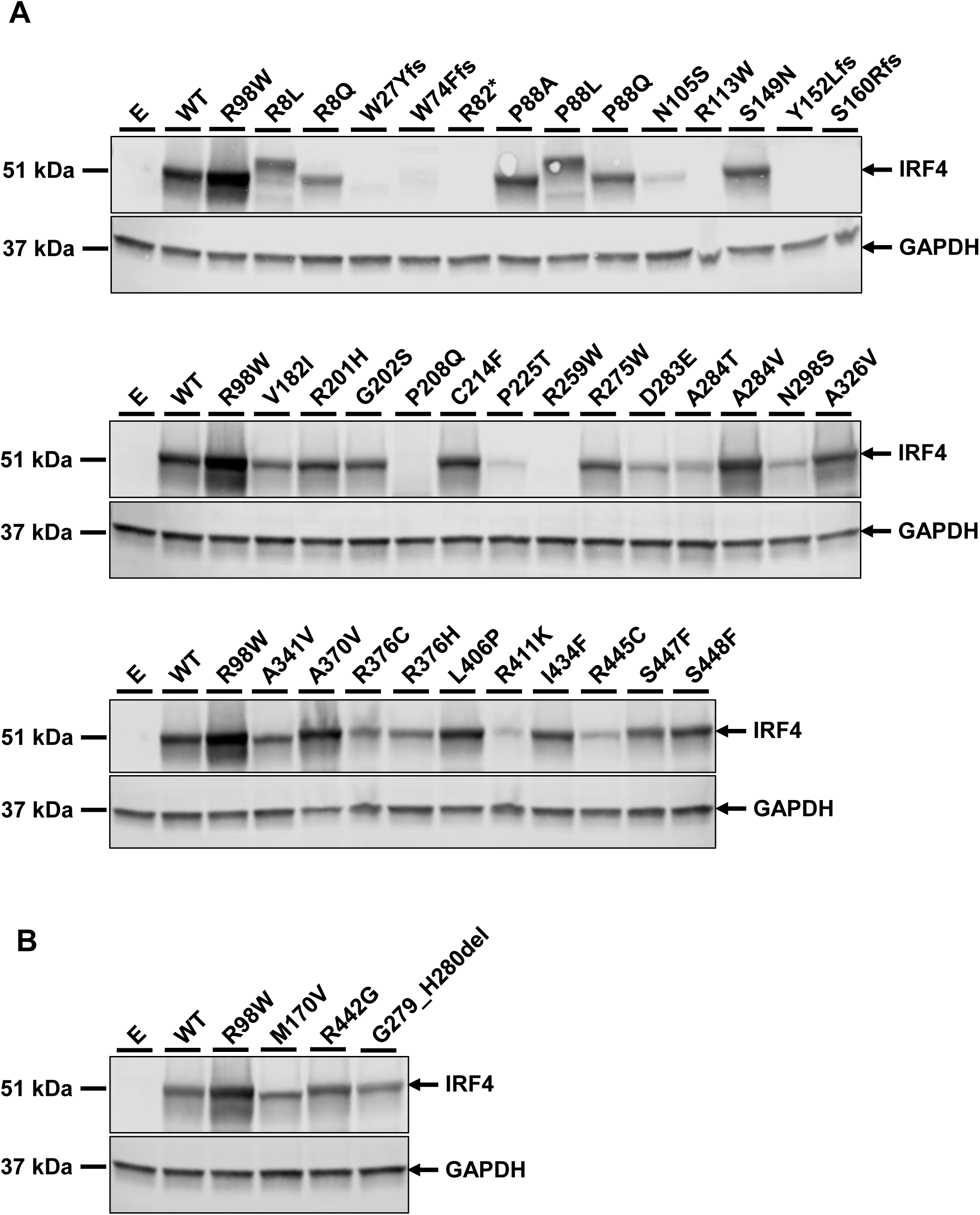
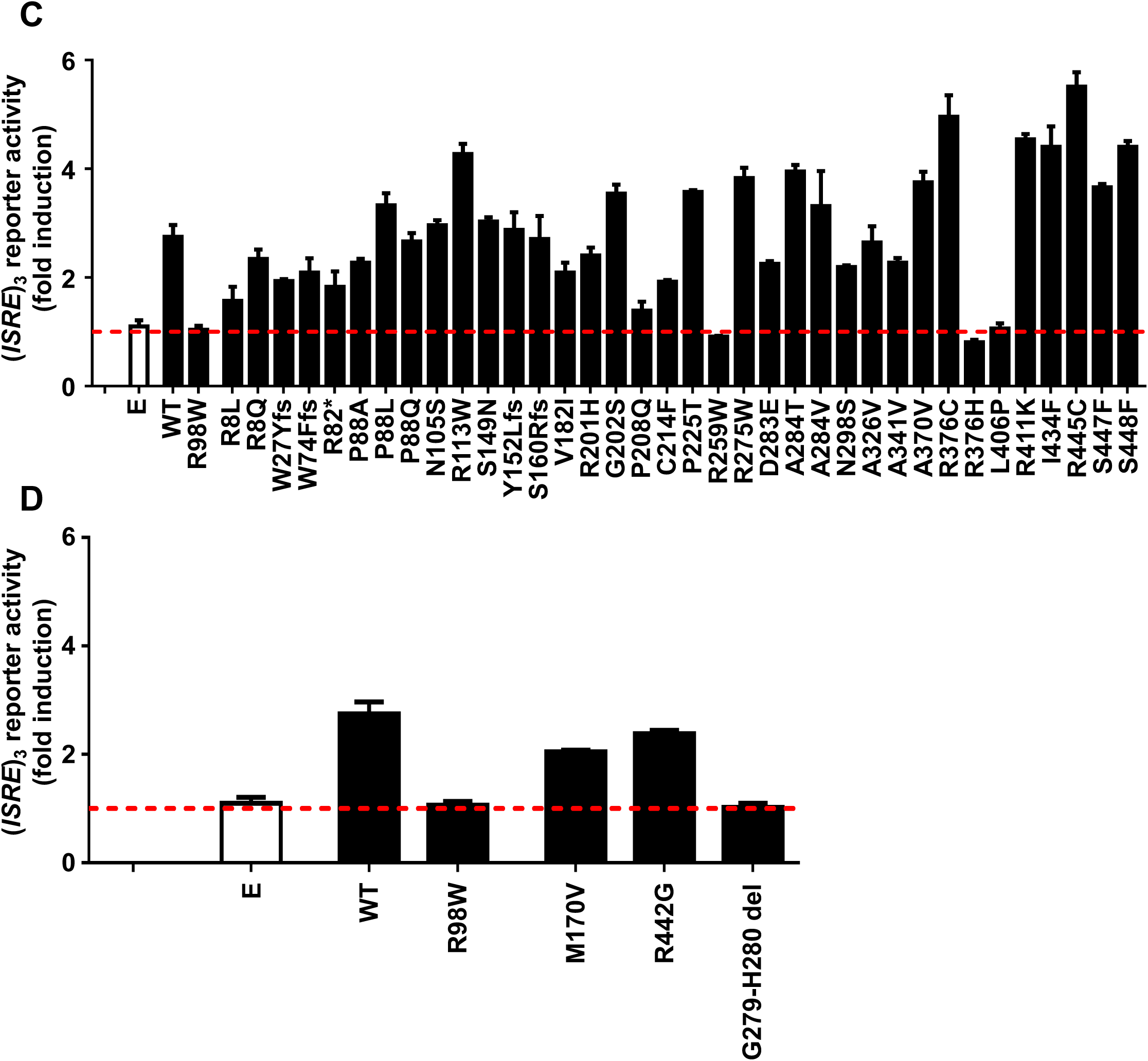
Protein levels and functional impact of *IRF4* variants. **A.-B.** HEK293-T cells were transfected with E or WT, R98W or several *IRF4* variants reported in a public database (**A**, see Figure S3B) or in the HGID database (**B**, see Figure S3C). Total cell extracts were subjected to western blotting; the upper panel shows IRF4 levels and the lower panel shows GAPDH levels, used as a loading control. The results shown are representative of three independent experiments. **C.-D.** Luciferase activity of HEK293-T cells cotransfected with an (*ISRE*)_3_ reporter plasmid plus E and plasmids containing WT, R98W or several IRF4 variants reported in public databases (**C**, see Figure S3B) or in the HGID database (**D**, see Figure S3C). Results are shown as fold induction of activity relative to E-transfected cells. The red dotted line indicates the mean fold induction in E-transfected cells. The results shown are the mean ± SD of three independent experiments.

**Supplementary Figure 6.**
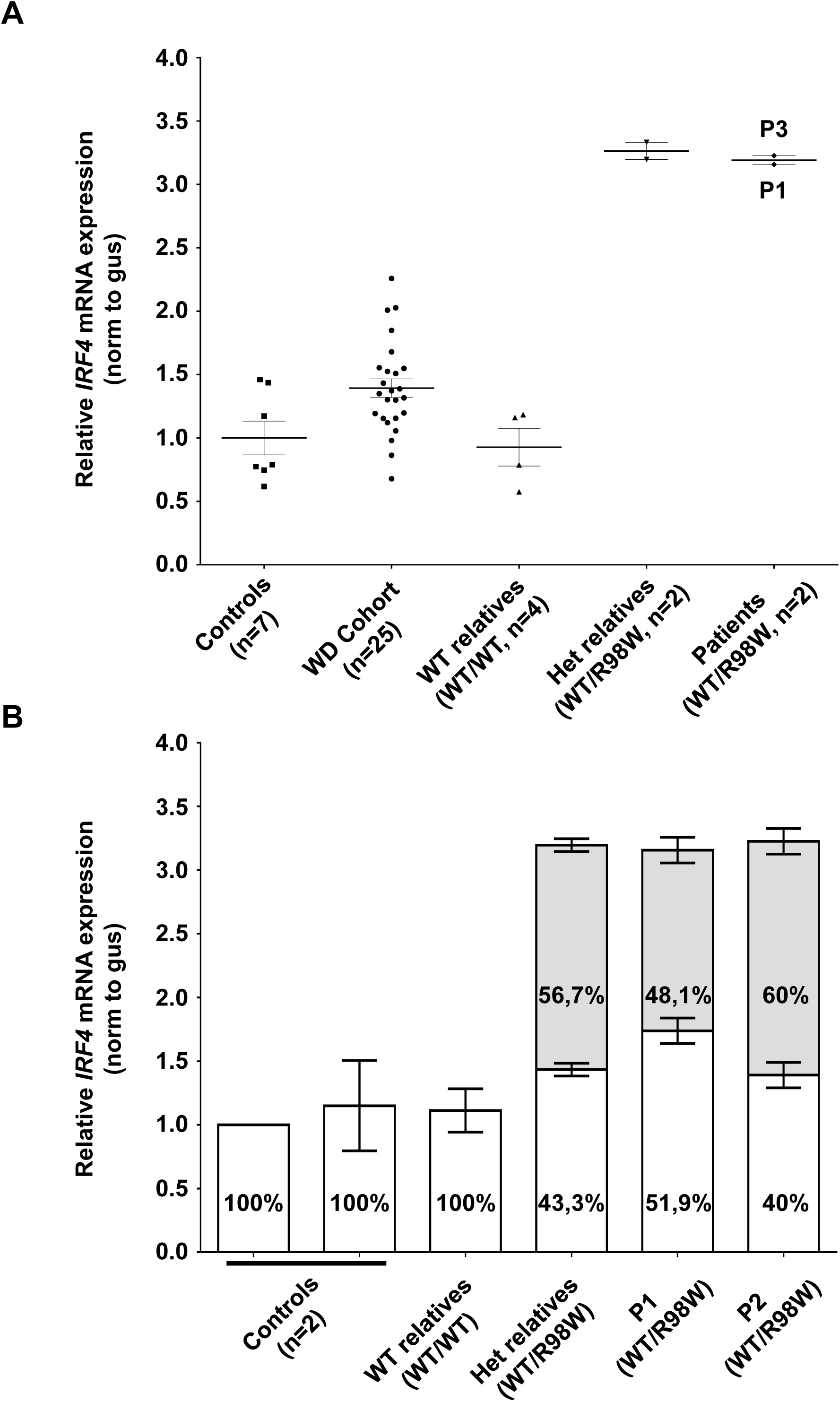
*IRF4* mRNA levels in EBV-B cells. **A.** Total RNA extracted from controls (n=7), patients diagnosed with Whipple’s disease (n=25) not related to this kindred, healthy homozygous WT relatives (*n*=4), patients with monoallelic *IRF4* mutations (*n*=2) and asymptomatic heterozygous relatives with monoallelic *IRF4* mutations (*n*=2) was subjected to RT-qPCR for total *IRF4*. Data are displayed as 2-ΔΔCt after normalization according to endogenous GUS control gene expression (ΔCt) and the mean of controls (ΔΔCt). The results shown are the mean ± SD of three independent experiments. **B.** The *IRF4* mRNA levels reported in Figure S7A are represented on histograms; the percentage within each bar is the deduced frequency of each mRNA obtained by the TA-cloning of cDNA generated from EBV-B cells from controls (*n*=2), healthy homozygous WT relatives (*n*=1), patients with monoallelic *IRF4* mutations (*n*=2) and asymptomatic heterozygous relatives with monoallelic *IRF4* mutations (*n*=1).

**Supplementary Figure 7.**
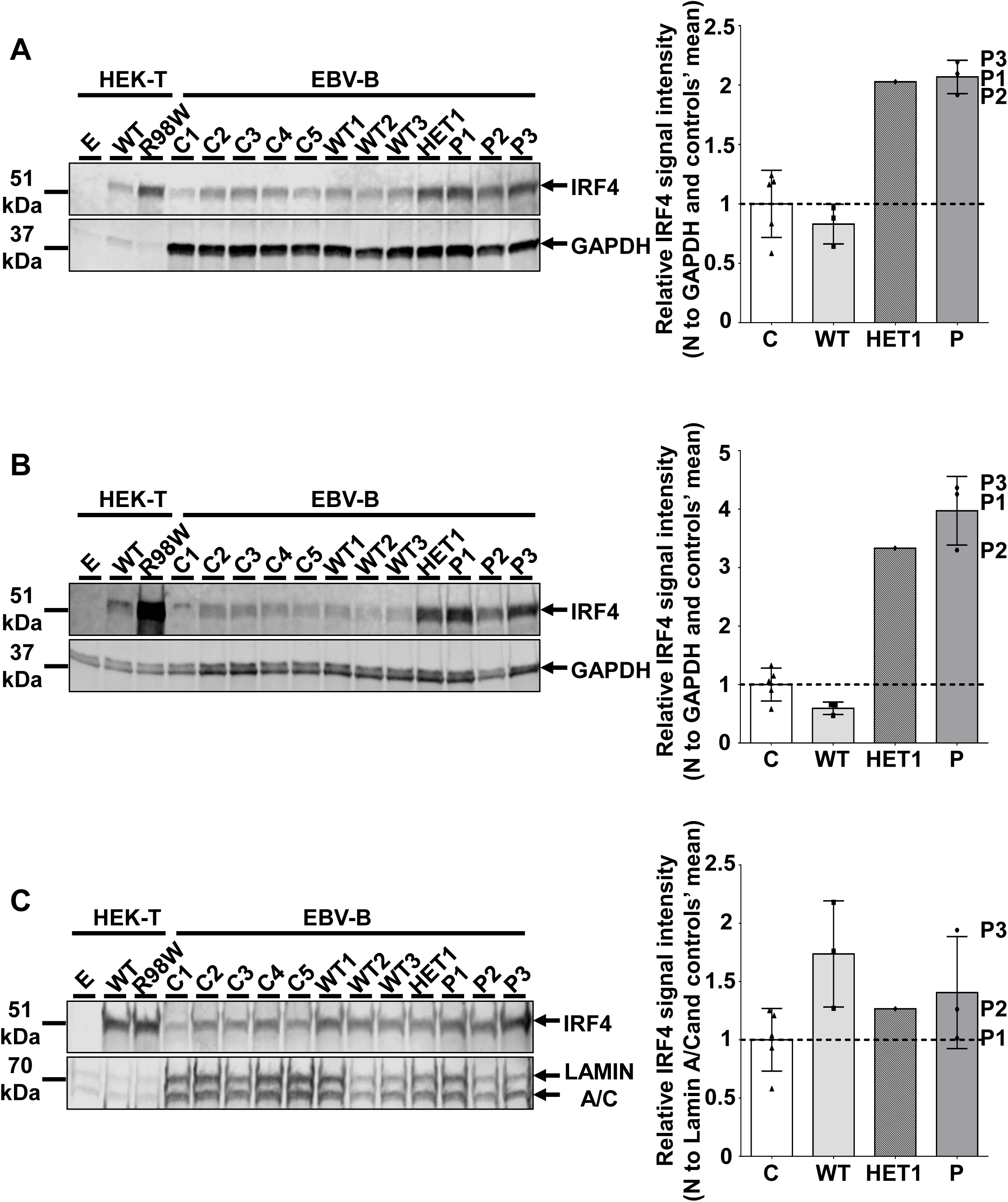
IRF4 protein levels in EBV-B cells. **A.-C.** (Left) Total cell (A), cytoplasmic (B) and nuclear (C) extracts (b) from five healthy controls (C1 to C5), three homozygous WT relatives (WT1, WT2, WT3), three patients (P1 to P3) and one asymptomatic heterozygous relative from the kindred (HET1). Protein extracts from HEK293-T cells transfected with E, WT or R98W plasmids were used as controls for the specific band corresponding to IRF4. (Right) Representation of IRF4 signal intensity for each individual relative to the mean signal for unrelated controls (*n*=5) obtained by WB and represented by black dotted lines (Supp. Figure A left) normalized against the GAPDH signal (total, cytoplasmic extracts) or the lamin A/C signal (nuclear extracts). The results shown are representative of two independent experiments.

**Supplementary Figure 8.**
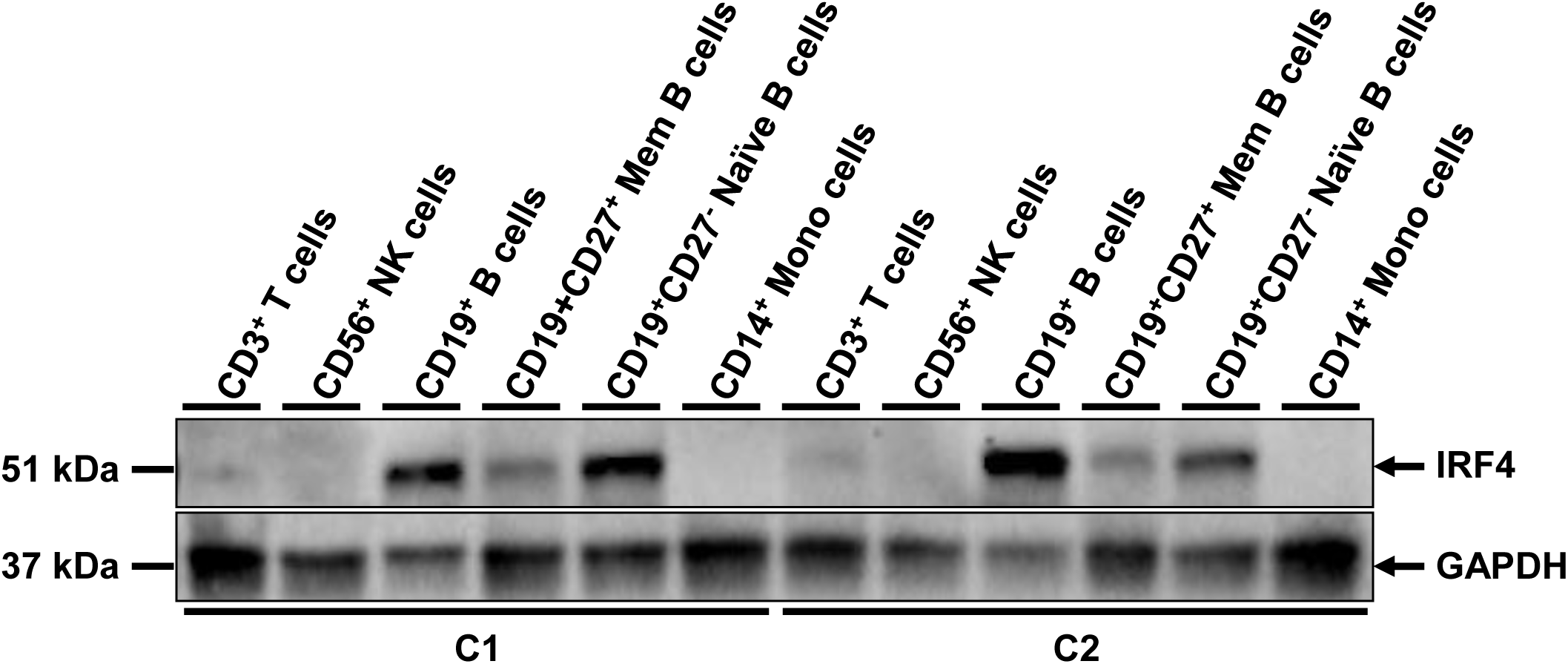
IRF4 protein levels in PBMC subpopulations. Total cell extracts from PBMC subpopulations (CD3^+^ T cells, CD56^+^ NK cells, CD19^+^ B cells, CD19^+^CD27^+^ memory B cells, CD19^+^CD27^−^ naive B cells, CD14^+^ monocytes) from two unrelated healthy controls were subjected to western blotting. The upper panel shows IRF4 levels and the lower panel shows GAPDH levels, used as a loading control. The results shown are representative of two independent experiments.

**Supplementary Figure 9.**
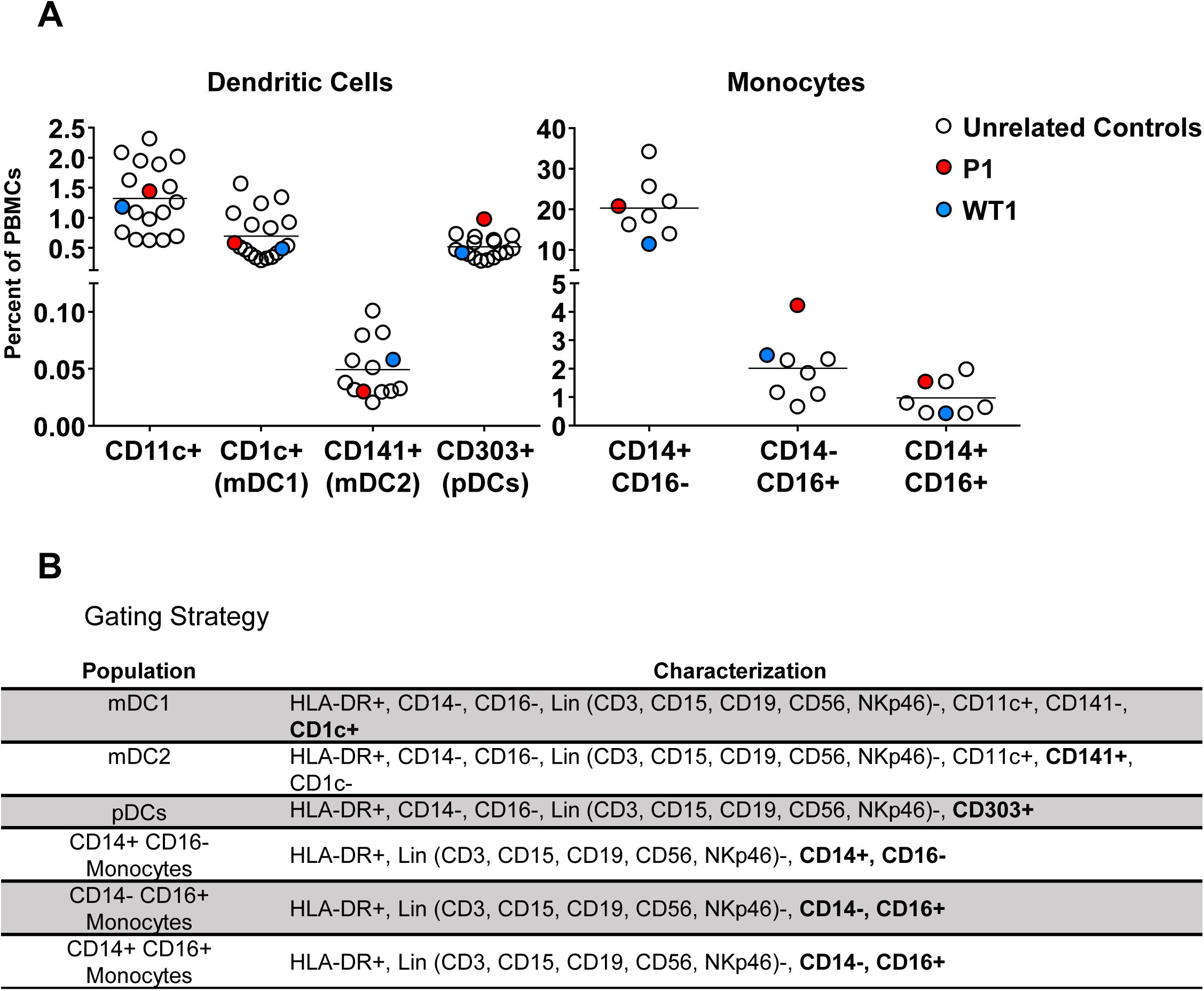
Percentage of dendritic cells and monocyte subtypes within total PBMCs. **A.** Percentage of CD11c^+^, myeloid dendritic cells (mDC1 and mDC2) and plasmacytoid dendritic cells (pDCs) (left), and monocyte subtypes (right) among total PBMCs from unrelated controls, a patient (P1) and a homozygous WT relative (WT1). **B.** Gating strategy to define the dendritic cell and monocyte subtypes.

**Supplementary Figure 10.**
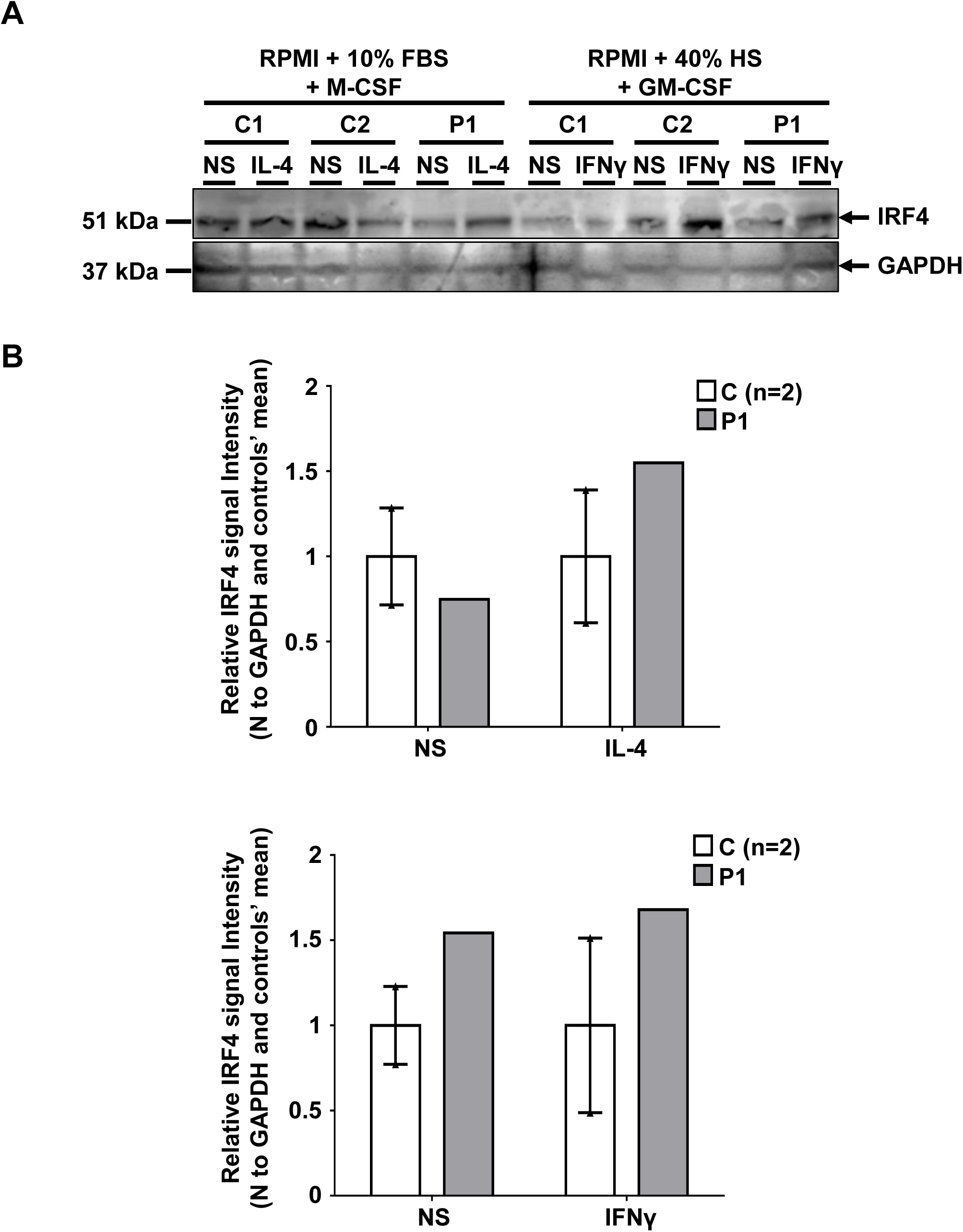
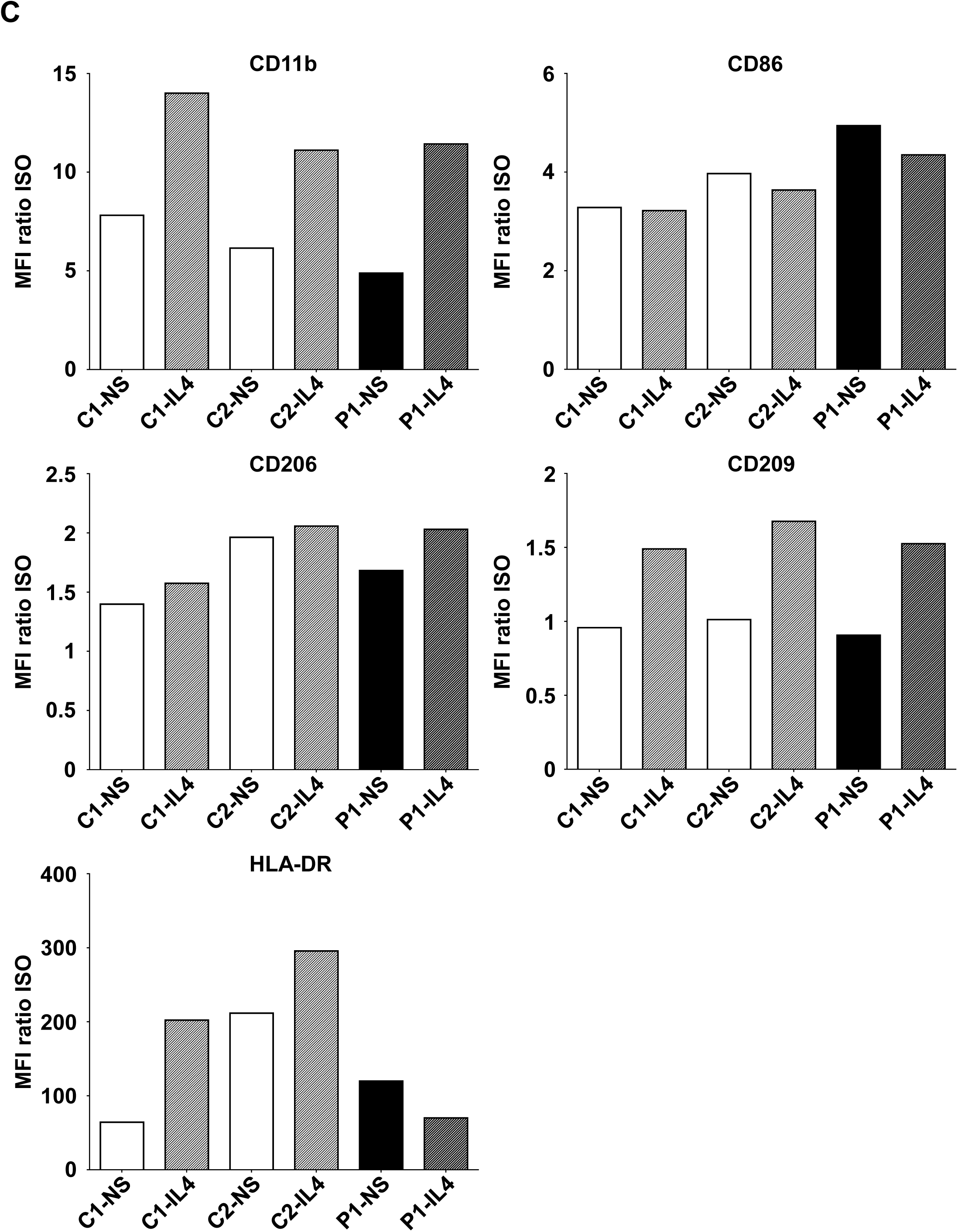
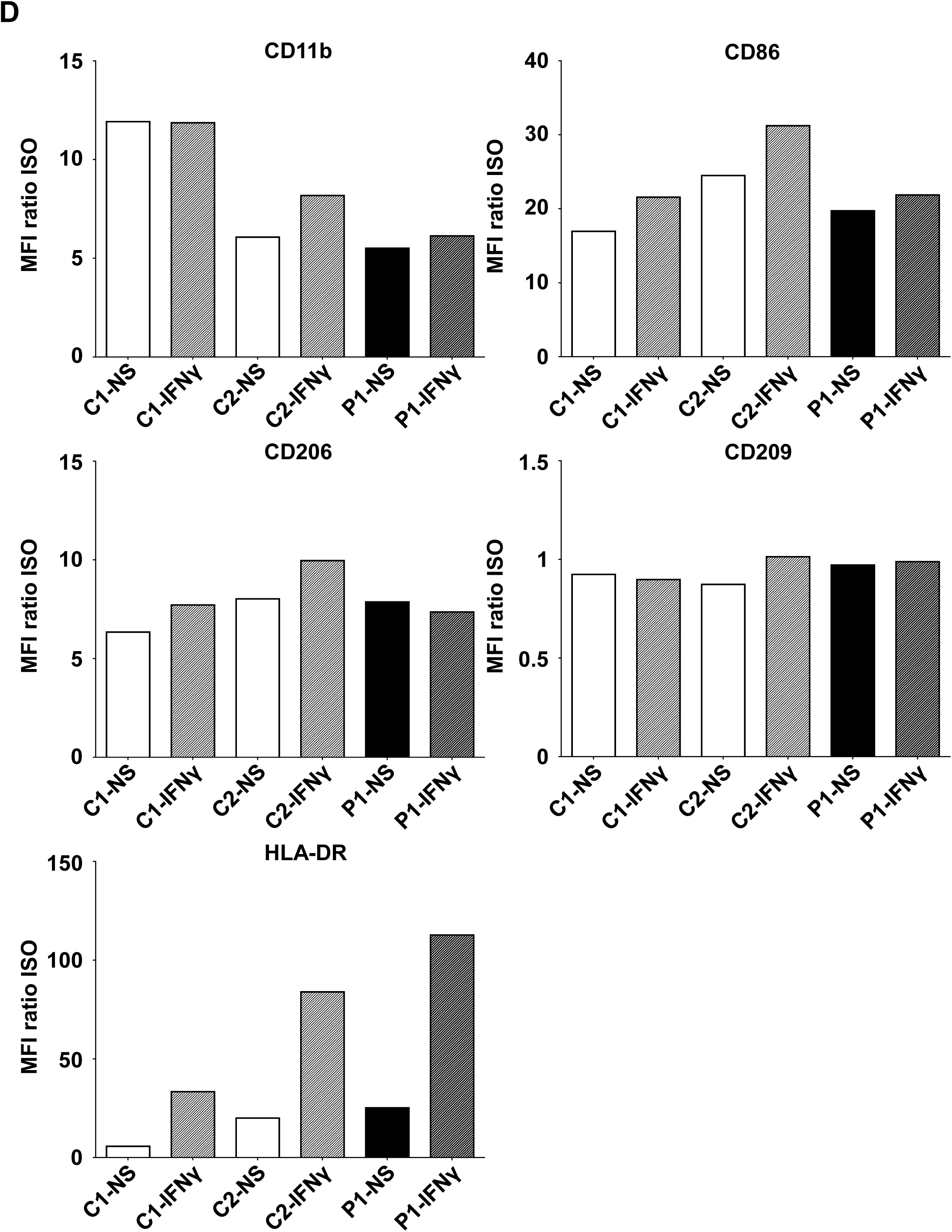
IRF4 levels in the patients’ monocyte-derived macrophages. **A.** IRF4 protein levels, as determined by western blotting on total cell extracts from M2-like (left panel) or M1-like (right panel) monocyte-derived macrophages (MDMs) from two unrelated healthy controls (C1 and C2) and P1, either left non-stimulated (NS) or stimulated with IL-4 (for M2-like MDMs) or IFN-γ (for M1-like MDMs). **B.** IRF4 signal intensity for each individual relative to the mean signal for controls on western blots. **C.-D.** CD11b, CD86, CD206, CD209 and HLA-DR mean fluorescence intensity (MFI) for M2-MDM (C) and M1-MDM (D) from P1 and two healthy unrelated controls (C1 and C2), either left non-stimulated (NS) or stimulated with IL-4 (for M2-like MDMs) or IFN-*γ* (for M1-like MDMs).

**Supplementary Figure 11.**
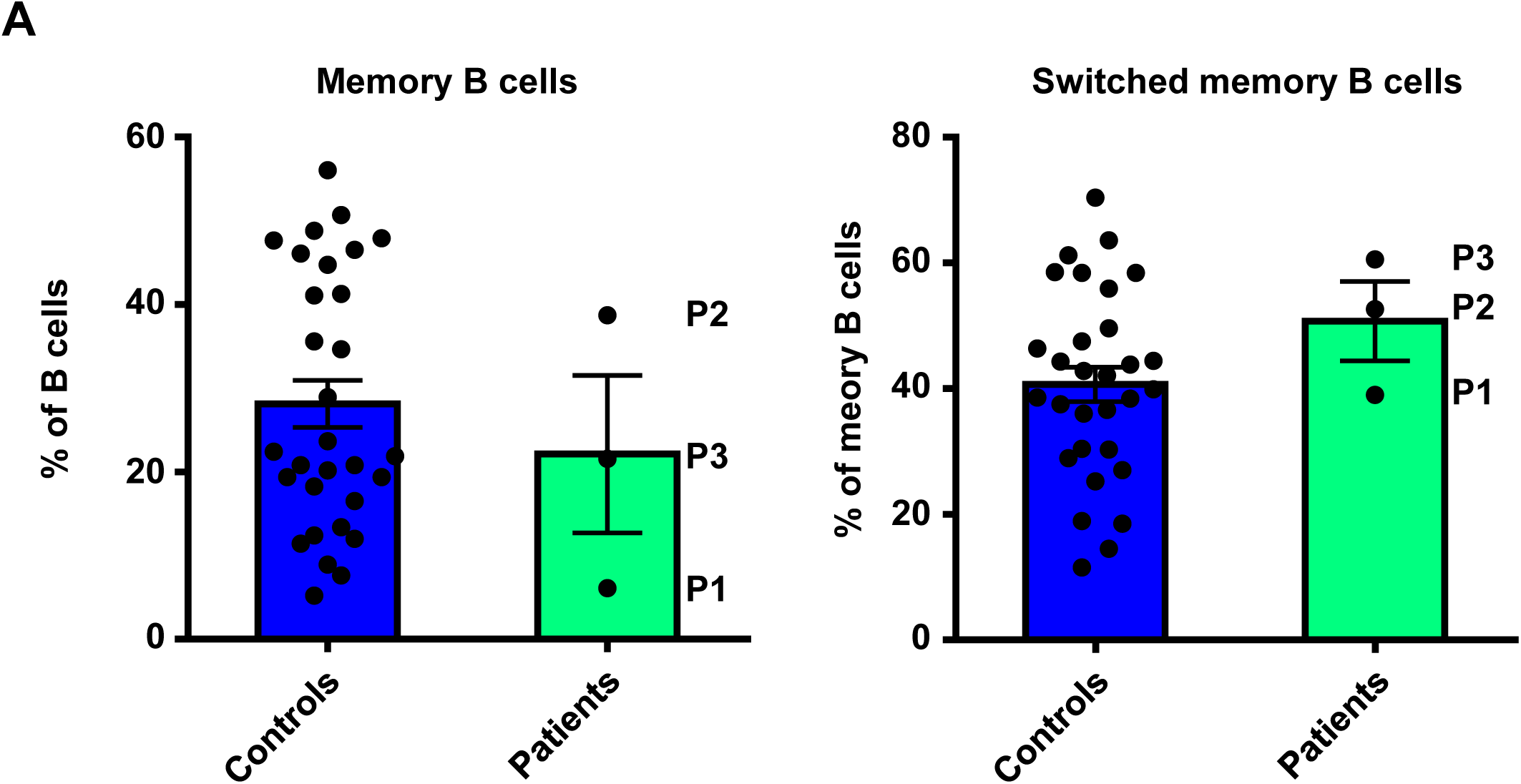
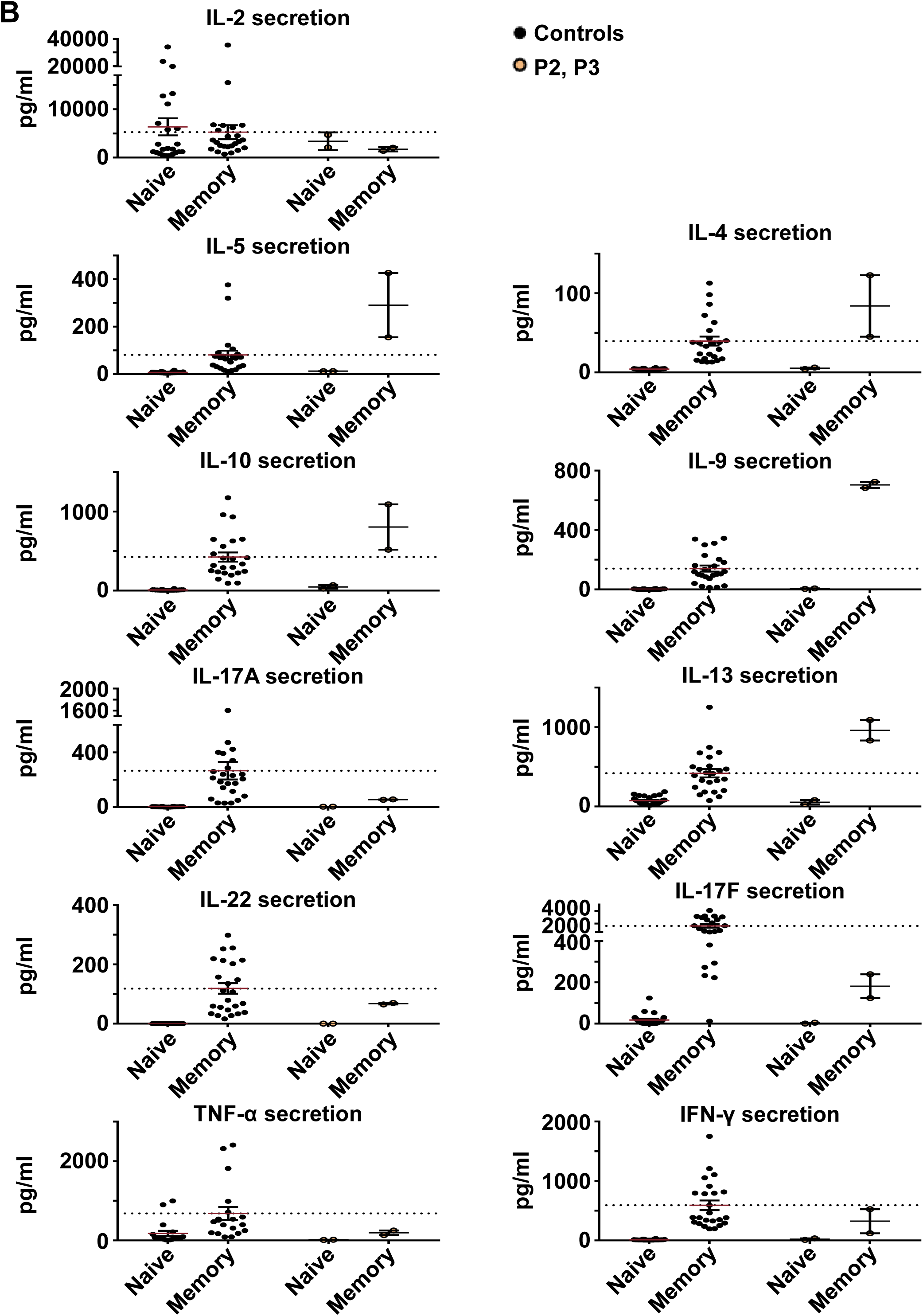
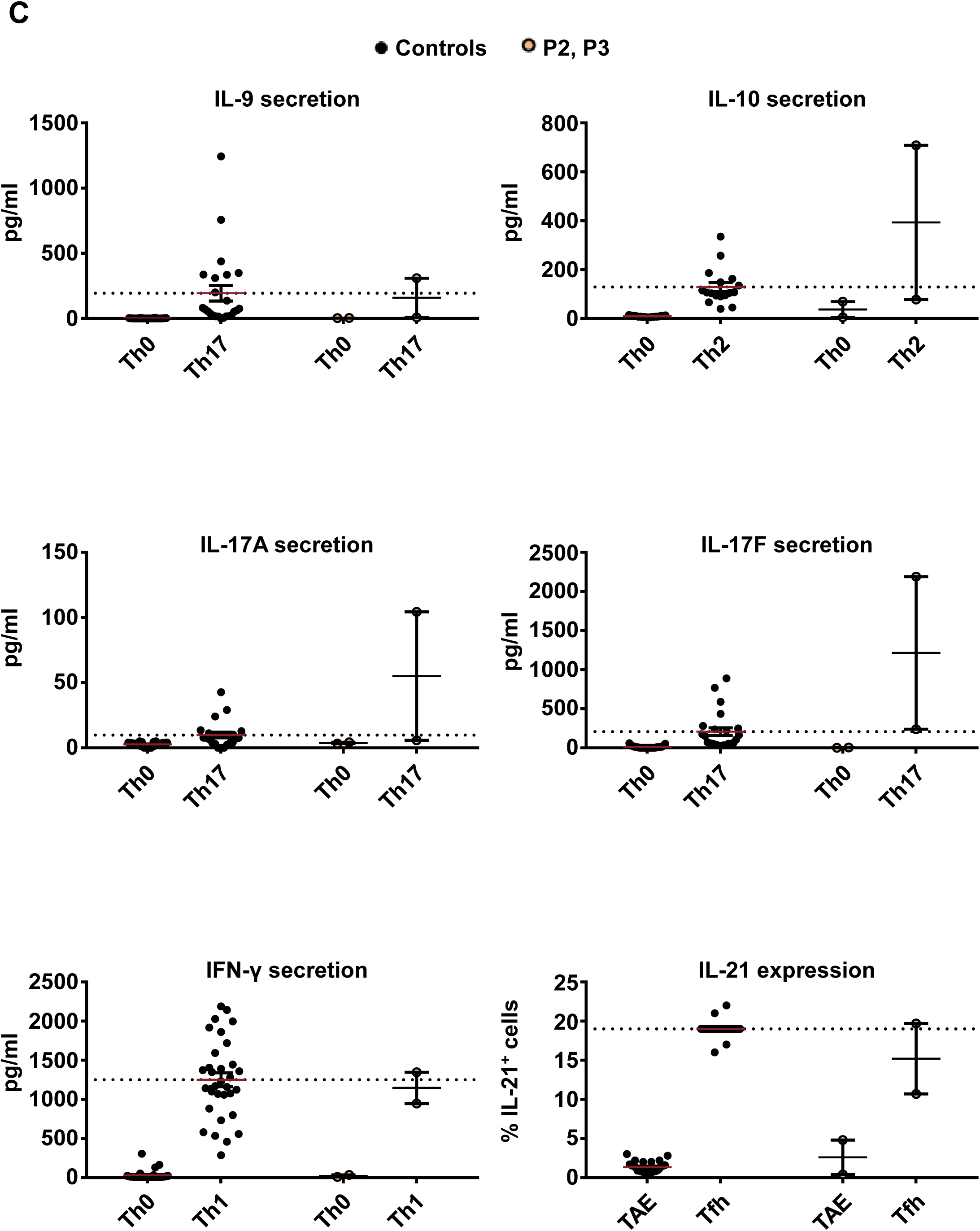
Percentage of memory B cells, *in vitro* differentiation of CD4_+_ T cells and *ex vivo* cytokine production by CD4_+_ memory T cells. **A.** PBMCs from unrelated controls and patients (P1, P2 and P3) were stained with antibodies against CD20, CD10 and CD27, IgM, IgG or IgA. Percentages of memory B cells (CD20^+^ CD10^−^ CD27^+^) were determined, and the proportion of memory B cells that had undergone class switching to express IgG or IgA was then calculated. No significant differences were observed between unrelated controls and patients. **B.** Naïve and memory CD4^+^ T cells from unrelated controls and patients (P2 and P3) were purified by sorting and cultured with TAE beads. The secretion of IL-2, IL-4, IL-5, IL-9, IL-10, IL-13, IL-17A, IL-17F, IL-22, IFN-γ and TNF-α was measured five days later. No significant differences were observed between unrelated controls and patients. **C.** Naïve CD4^+^ T cells from unrelated controls and patients (P2 and P3) were stimulated with TAE beads alone or under Th1, Th2, Th17 or Tfh polarizing conditions. The production of IL-10, IL-21, IL-17A, IL-17F and IFN-γ was measured five days later, in the corresponding polarizing conditions. No significant differences were observed (as in B) between unrelated controls and patients.

**Supplementary Table 1.** Kindred information summary. For each subject, Tw carriage status, *IRF4* genotype, clinical status and date of birth (DOB) are reported. NA: not available; Pos: positive; Neg: negative; Tw: *Tropheryma whipplei*; WD: Whipple’s disease; E?: genotype not assessed.

| ID | TW carriage (PCR) |  |  | IRF4 genotype | Clinical status | DOB |
| --- | --- | --- | --- | --- | --- | --- |
|  | Feces | Saliva | Joint fluid |  |  |  |
| III.4 (P3) | Pos. then Neg. | Neg. | NA | WT/R98W | Classical WD | 21/05/1925 |
| III.5 | Neg. | Neg. | NA | WT/WT | Healthy | 14/08/1922 |
| III.6 | NA | Pos. | NA | WT/R98W | Healthy | 01/04/1935 |
| IV.1 (P4) | Neg. | Neg. | NA | WT/R98W | Althralgia WD | 14/04/1947 |
| IV.2 | Neg. | Neg. | NA | WT/WT | Healthy | 20/07/1951 |
| IV.3 (P2) | Pos. | Pos. | NA | WT/R98W | Classical WD | 30/08/1941 |
| IV.4 | Pos. | Pos. | NA | WT/R98W | Healthy | 23/07/1955 |
| IV.5 | Neg. | Neg. | NA | WT/R98W | Healthy | 06/11/1951 |
| IV.6 (P1) | Neg. | Neg. | Pos. | WT/R98W | Althralgia WD | 17/02/1948 |
| IV.7 | Neg. | Neg. | NA | WT/WT | Healthy | 17/02/1948 |
| IV.8 | Neg. | Neg. | NA | WT/WT | Healthy | 09/04/1961 |
| IV.9 | NA | NA | NA | WT/R98W | Healthy | 26/04/1958 |
| V.1 | Neg. | Neg. | NA | WT/WT | Healthy | 13/05/1974 |
| V.2 | Neg. | Neg. | NA | WT/WT | Healthy | 07/05/1978 |
| V.3 | Pos. | Neg. | NA | WT/R98W | Healthy | 11/09/1965 |
| V.4 | Pos. | Pos. | NA | WT/R98W | Healthy | 14/07/1962 |
| V.5 | NA | NA | NA | WT/WT | Healthy | 09/08/1969 |
| V.6 | NA | NA | NA | E? | Healthy | NA |
| V.7 | NA | NA | NA | WT/WT | Healthy |  |
| V.8 | NA | NA | NA | WT/WT | Healthy | 31/08/1994 |
| V.9 | NA | NA | NA | E? | Healthy | NA |
| V.10 | NA | NA | NA | E? | Healthy | NA |
| VI.1 | Neg. | Neg. | NA | WT/WT | Healthy | 17/02/2004 |
| VI.2 | Neg. | Neg. | NA | WT/WT | Healthy | 21/09/2006 |
| VI.3 | NA | Neg. | NA | WT/WT | Healthy | 16/05/2006 |
| VI.4 | Neg. | Neg. | NA | WT/WT | Healthy | 29/03/2011 |
| VI.5 | Neg. | Neg. | NA | WT/WT | Healthy | 09/08/1996 |
| VI.6 | Pos. | Neg. | NA | WT/R98W | Healthy | 04/10/1993 |
| VI.7 | Neg. | Neg. | NA | WT/R98W | Healthy | 24/08/2002 |
| VI.8 | NA | NA | NA | E? | Healthy | NA |
| VI.9 | NA | NA | NA | E? | Healthy | NA |

**Supplementary Table S2.**
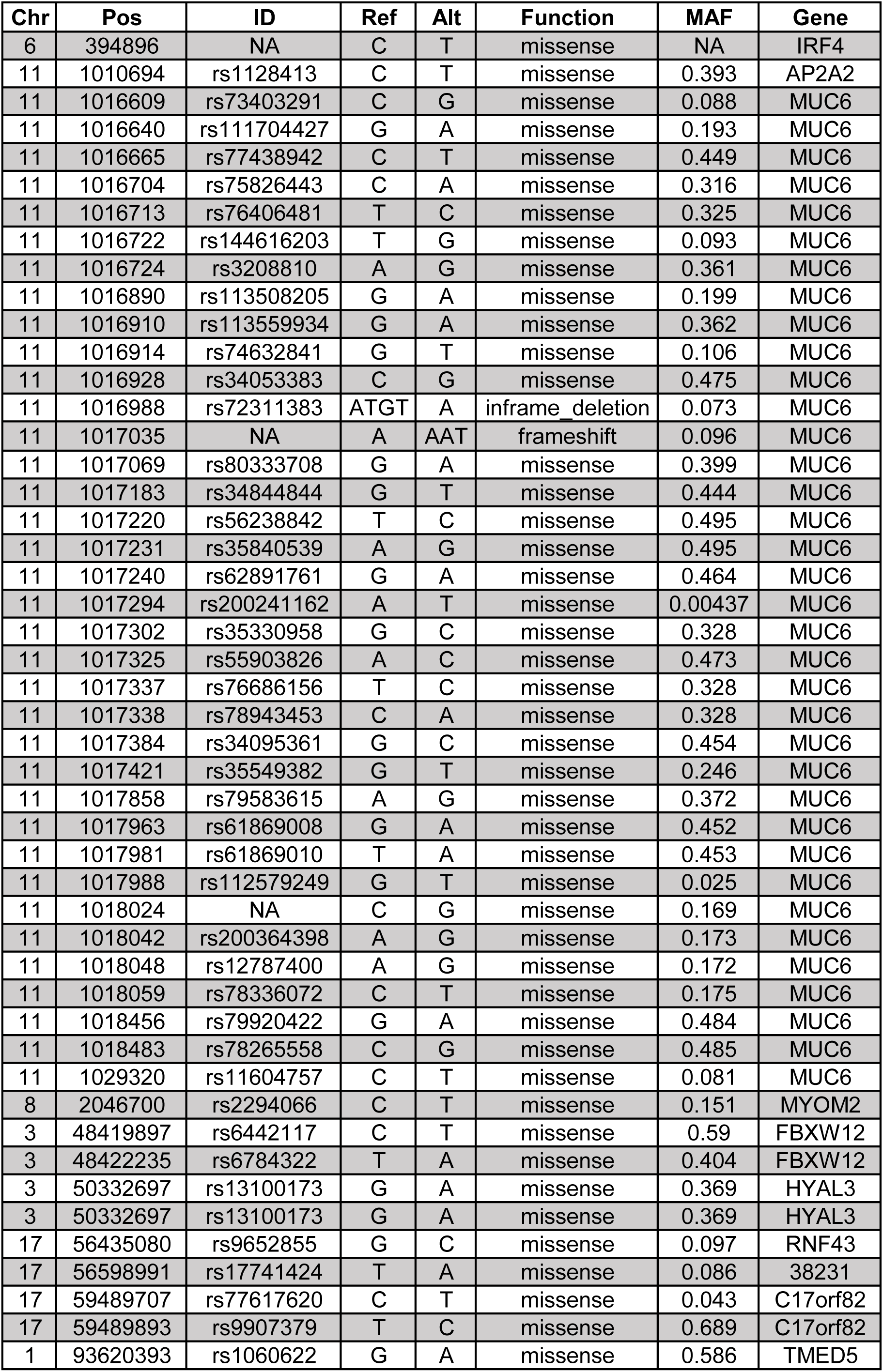
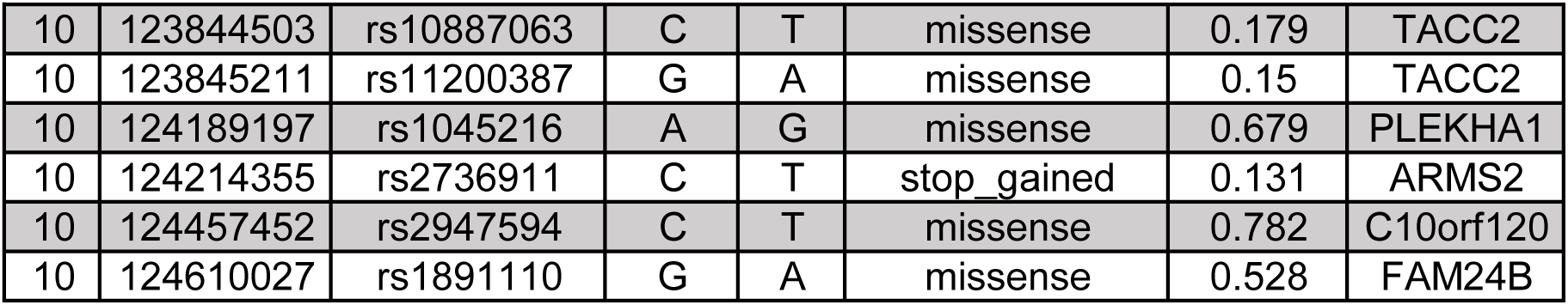
Non-synonymous variants within the linkage regions found in WES data from patients.

| Chr | Pos | ID | Ref | Alt | Function | MAF | Gene |
| --- | --- | --- | --- | --- | --- | --- | --- |
| 6 | 394896 | NA | C | T | missense | NA | IRF4 |
| 11 | 1010694 | rs1128413 | C | T | missense | 0.393 | AP2A2 |
| 11 | 1016609 | rs73403291 | C | G | missense | 0.088 | MUC6 |
| 11 | 1016640 | rs111704427 | G | A | missense | 0.193 | MUC6 |
| 11 | 1016665 | rs77438942 | C | T | missense | 0.449 | MUC6 |
| 11 | 1016704 | rs75826443 | C | A | missense | 0.316 | MUC6 |
| 11 | 1016713 | rs76406481 | T | C | missense | 0.325 | MUC6 |
| 11 | 1016722 | rs144616203 | T | G | missense | 0.093 | MUC6 |
| 11 | 1016724 | rs3208810 | A | G | missense | 0.361 | MUC6 |
| 11 | 1016890 | rs113508205 | G | A | missense | 0.199 | MUC6 |
| 11 | 1016910 | rs113559934 | G | A | missense | 0.362 | MUC6 |
| 11 | 1016914 | rs74632841 | G | T | missense | 0.106 | MUC6 |
| 11 | 1016928 | rs34053383 | C | G | missense | 0.475 | MUC6 |
| 11 | 1016988 | rs72311383 | ATGT | A | inframe_deletion | 0.073 | MUC6 |
| 11 | 1017035 | NA | A | AAT | frameshift | 0.096 | MUC6 |
| 11 | 1017069 | rs80333708 | G | A | missense | 0.399 | MUC6 |
| 11 | 1017183 | rs34844844 | G | T | missense | 0.444 | MUC6 |
| 11 | 1017220 | rs56238842 | T | C | missense | 0.495 | MUC6 |
| 11 | 1017231 | rs35840539 | A | G | missense | 0.495 | MUC6 |
| 11 | 1017240 | rs62891761 | G | A | missense | 0.464 | MUC6 |
| 11 | 1017294 | rs200241162 | A | T | missense | 0.00437 | MUC6 |
| 11 | 1017302 | rs35330958 | G | C | missense | 0.328 | MUC6 |
| 11 | 1017325 | rs55903826 | A | C | missense | 0.473 | MUC6 |
| 11 | 1017337 | rs76686156 | T | C | missense | 0.328 | MUC6 |
| 11 | 1017338 | rs78943453 | C | A | missense | 0.328 | MUC6 |
| 11 | 1017384 | rs34095361 | G | C | missense | 0.454 | MUC6 |
| 11 | 1017421 | rs35549382 | G | T | missense | 0.246 | MUC6 |
| 11 | 1017858 | rs79583615 | A | G | missense | 0.372 | MUC6 |
| 11 | 1017963 | rs61869008 | G | A | missense | 0.452 | MUC6 |
| 11 | 1017981 | rs61869010 | T | A | missense | 0.453 | MUC6 |
| 11 | 1017988 | rs112579249 | G | T | missense | 0.025 | MUC6 |
| 11 | 1018024 | NA | C | G | missense | 0.169 | MUC6 |
| 11 | 1018042 | rs200364398 | A | G | missense | 0.173 | MUC6 |
| 11 | 1018048 | rs12787400 | A | G | missense | 0.172 | MUC6 |
| 11 | 1018059 | rs78336072 | C | T | missense | 0.175 | MUC6 |
| 11 | 1018456 | rs79920422 | G | A | missense | 0.484 | MUC6 |
| 11 | 1018483 | rs78265558 | C | G | missense | 0.485 | MUC6 |
| 11 | 1029320 | rs11604757 | C | T | missense | 0.081 | MUC6 |
| 8 | 2046700 | rs2294066 | C | T | missense | 0.151 | MYOM2 |
| 3 | 48419897 | rs6442117 | C | T | missense | 0.59 | FBXW12 |
| 3 | 48422235 | rs6784322 | T | A | missense | 0.404 | FBXW12 |
| 3 | 50332697 | rs13100173 | G | A | missense | 0.369 | HYAL3 |
| 3 | 50332697 | rs13100173 | G | A | missense | 0.369 | HYAL3 |
| 17 | 56435080 | rs9652855 | G | C | missense | 0.097 | RNF43 |
| 17 | 56598991 | rs17741424 | T | A | missense | 0.086 | 38231 |
| 17 | 59489707 | rs77617620 | C | T | missense | 0.043 | C17orf82 |
| 17 | 59489893 | rs9907379 | T | C | missense | 0.689 | C17orf82 |
| 1 | 93620393 | rs1060622 | G | A | missense | 0.586 | TMED5 |
| 10 | 123844503 | rs10887063 | C | T | missense | 0.179 | TACC2 |
| 10 | 123845211 | rs11200387 | G | A | missense | 0.15 | TACC2 |
| 10 | 124189197 | rs1045216 | A | G | missense | 0.679 | PLEKHA1 |
| 10 | 124214355 | rs2736911 | C | T | stop_gained | 0.131 | ARMS2 |
| 10 | 124457452 | rs2947594 | C | T | missense | 0.782 | C10orf120 |
| 10 | 124610027 | rs1891110 | G | A | missense | 0.528 | FAM24B |

**Supplementary Table S3.** 156 non-synonymous heterozygous coding or splice variants reported in the GnomAD and/or HGID databases.

| Chr | Pos | ID | Ref | Alt | fuction | AACChange | CADD | MAF |
| --- | --- | --- | --- | --- | --- | --- | --- | --- |
| 6 | 393174 | . | C | G | missense | p.Arg8Gly | 26.8 | 6.71E-06 |
| 6 | 393175 | rs139884486 | G | A | missense | p.Arg8Gln | 25.7 | 0.000189 |
| 6 | 393175 | rs139884486 | G | T | missense | p.Arg8Leu | 28 | 4.99E-05 |
| 6 | 393231 | rs766107516 | TGGCTG | T | frameshift | p.Trp27TyrfsTer50 | 35 | 5.51E-06 |
| 6 | 393256 | rs538907751 | G | C | missense | p.Gly35Ala | 24.4 | 4.84E-06 |
| 6 | 393264 | rs369688140 | C | T | missense | p.Pro38Ser | 29.5 | 3.67E-05 |
| 6 | 393273 | rs751037944 | G | C | missense | p.Val41Leu | 24.7 | 2.69E-05 |
| 6 | 393284 | rs753781892 | CGAG | C | inframe del | p.Glu46del | 22.9 | 8.68E-06 |
| 6 | 393292 | . | A | G | missense | p.Lys47Arg | 22.8 | 1.90E-05 |
| 6 | 394822 | rs754867881 | CT | C | frameshift | p.Trp74GlyfsTer28 | 24 | 4.10E-06 |
| 6 | 394848 | . | C | T | stop gained | p.Arg82Ter | 40 | 4.07E-06 |
| 6 | 394849 | . | G | A | missense | p.Arg82Gln | 24.4 | 4.07E-06 |
| 6 | 394859 | . | C | G | missense | p.Ile85Met | 12.09 | 3.23E-05 |
| 6 | 394866 | rs144593192 | C | G | missense | p.Pro88Ala | 18.71 | 3.61E-05 |
| 6 | 394867 | rs202124383 | C | A | missense | p.Pro88Gln | 30 | 0.000541 |
| 6 | 394867 | rs202124383 | C | T | missense | p.Pro88Leu | 33 | 4.06E-06 |
| 6 | 394902 | rs767879003 | G | A | missense | p.Ala100Thr | 24.7 | 4.06E-06 |
| 6 | 394918 | rs200303421 | A | G | missense | p.Asn105Ser | 13.56 | 5.77E-05 |
| 6 | 394932 | rs374588785 | C | A | missense | p.Leu110Met | 24 | 1.62E-05 |
| 6 | 394941 | rs753842732 | C | T | missense | p.Arg113Trp | 32 | 4.06E-06 |
| 6 | 394981 | rs745876631 | G | A | missense | p.Arg126Lys | 22.9 | 8.24E-06 |
| 6 | 394986 | rs771936972 | G | A | missense | p.Val128Ile | 18.17 | 4.27E-06 |
| 6 | 395004 | rs748154669 | A | G | missense | p.Lys134Glu | 21.6 | 4.31E-06 |
| 6 | 395009 | rs200504236 | T | C | splice donor | c.403+2T>C | 23.9 | 5.22E-05 |
| 6 | 395864 | rs11557493 | C | G | missense | p.Leu141Val | 15.25 | 1.63E-05 |
| 6 | 395881 | rs115613112 | G | A | missense | p.Met146Ile | 15.67 | 1.45E-05 |
| 6 | 395885 | . | A | G | missense | p.Met148Val | 3.441 | 4.08E-06 |
| 6 | 395886 | . | T | C | missense | p.Met148Thr | 8.28 | 3.23E-05 |
| 6 | 395889 | rs73717071 | G | A | missense | p.Ser149Asn | 5.825 | 0.008454 |
| 6 | 395892 | . | A | AT | frameshift | p.Tyr152LeufsTer60 | 27.8 | 4.08E-06 |
| 6 | 395894 | . | C | T | missense | p.Pro151Ser | 13.44 | 4.08E-06 |
| 6 | 395898 | . | A | G | missense | p.Tyr152Cys | 10.74 | 7.24E-06 |
| 6 | 395903 | rs764577689 | A | G | missense | p.Met154Val | 0.009 | 1.09E-05 |
| 6 | 395903 | rs764577689 | A | T | missense | p.Met154Leu | 0.09 | 4.08E-06 |
| 6 | 395906 | rs776661032 | A | C | missense | p.Thr155Pro | 0.005 | 4.09E-06 |
| 6 | 395916 | rs750505973 | A | T | missense | p.Tyr158Phe | 8.071 | 1.23E-05 |
| 6 | 395919 | rs771434117 | CT | C | frameshift | p.Ser160ArgfsTer11 | 21.4 | 4.10E-06 |
| 6 | 395927 | rs766413794 | C | G | missense | p.Pro162Ala | 5.153 | 4.12E-06 |
| 6 | 397116 | rs767166788 | C | G | missense | p.His167Gln | 9.287 | 4.06E-06 |
| 6 | 397123 | . | A | G | missense | p.Met170Val | 8.301 | private |
| 6 | 397125 | . | G | A | missense | p.Met170Ile | 21.3 | 4.06E-06 |
| 6 | 397127 | rs755905242 | T | C | missense | p.Met171Thr | 14.97 | 2.84E-05 |
| 6 | 397128 | . | G | A | missense | p.Met171Ile | 18.47 | 4.06E-06 |
| 6 | 397129 | . | C | T | missense | p.Pro172Ser | 14.26 | 4.06E-06 |
| 6 | 397130 | rs777399667 | C | A | missense | p.Pro172Gln | 22.3 | 4.06E-06 |
| 6 | 397138 | . | G | A | missense | p.Asp175Asn | 23.6 | 4.06E-06 |
| 6 | 397154 | rs778403209 | A | G | missense | p.Asp180Gly | 19.46 | 2.03E-05 |
| 6 | 397155 | . | C | G | missense | p.Asp180Glu | 1.224 | 4.06E-06 |
| 6 | 397157 | rs747437180 | A | G | missense | p.Tyr181Cys | 23.6 | 4.06E-06 |
| 6 | 397159 | rs781586995 | G | A | missense | p.Val182Ile | 14.49 | 4.06E-06 |
| 6 | 397170 | rs770110760 | G | C | missense | p.Gln185His | 22.3 | 8.12E-06 |
| 6 | 397172 | . | C | T | missense | p.Pro186Leu | 23.6 | 4.06E-06 |
| 6 | 397175 | . | A | G | missense | p.His187Arg | 16.01 | 3.23E-05 |
| 6 | 397178 | . | C | T | missense | p.Pro188Leu | 22.9 | 1.08E-05 |
| 6 | 397183 | rs763272146 | A | G | missense | p.Ile190Val | 9.874 | 4.06E-06 |
| 6 | 397184 | rs771094772 | T | A | missense | p.Ile190Asn | 21.8 | 4.06E-06 |
| 6 | 397186 | rs774593906 | C | T | missense | p.Pro191Ser | 15.87 | 1.22E-05 |
| 6 | 397201 | . | A | G | missense | p.Met196Val | 0.002 | 7.21E-06 |
| 6 | 397205 | rs372869810 | C | T | missense | p.Thr197Met | 17.83 | 8.12E-06 |
| 6 | 397205 | rs372869810 | C | T | missense | p.Thr197Met | 17.83 | 8.12E-06 |
| 6 | 397208 | rs763891187 | T | C | missense | p.Phe198Ser | 10.38 | 8.12E-06 |
| 6 | 397213 | . | C | T | missense | p.Pro200Ser | 9.286 | 4.06E-06 |
| 6 | 397216 | . | C | T | missense | p.Arg201Cys | 25.8 | 4.06E-06 |
| 6 | 397217 | rs753427028 | G | A | missense | p.Arg201His | 24.2 | 1.22E-05 |
| 6 | 397219 | rs143144957 | G | A | missense | p.Gly202Ser | 0.004 | 7.58E-05 |
| 6 | 397234 | . | G | A | missense | p.Gly207Ser | 20.2 | 4.06E-06 |
| 6 | 397238 | rs757910134 | C | A | missense | p.Pro208Gln | 23.4 | 7.22E-05 |
| 6 | 398831 | rs538810711 | G | T | missense | p.Cys214Phe | 16.56 | 3.61E-05 |
| 6 | 398863 | . | C | A | missense | p.Pro225Thr | 24.6 | 6.46E-05 |
| 6 | 398878 | rs147454166 | G | A | missense | p.Ala230Thr | 18.45 | 1.63E-05 |
| 6 | 398884 | rs375133421 | G | A | missense | p.Gly232Arg | 24.4 | 4.06E-06 |
| 6 | 398887 | rs776593233 | G | A | missense | p.Val233Ile | 13.26 | 3.25E-05 |
| 6 | 398888 | rs761579830 | T | C | missense | p.Val233Ala | 16.11 | 4.06E-06 |
| 6 | 398894 | rs765027259 | C | A | missense | p.Thr235Lys | 16.75 | 4.07E-06 |
| 6 | 398914 | rs772851591 | G | A | missense | p.Ala242Thr | 24.7 | 8.15E-06 |
| 6 | 398915 | . | C | T | missense | p.Ala242Val | 21.5 | 4.08E-06 |
| 6 | 398925 | rs765864636 | G | C | missense | p.Leu245Phe | 22.7 | 4.10E-06 |
| 6 | 401425 | rs758839744 | C | G | missense | p.Asp249Glu | 26.2 | 4.11E-06 |
| 6 | 401430 | rs199834880 | G | A | missense | p.Arg251Gln | 23.8 | 8.20E-06 |
| 6 | 401438 | . | A | G | missense | p.Ile254Val | 8.678 | 4.08E-06 |
| 6 | 401444 | rs148760922 | C | A | missense | p.Leu256Met | 25.7 | 4.08E-06 |
| 6 | 401448 | . | A | T | missense | p.Tyr257Phe | 16.05 | 4.08E-06 |
| 6 | 401453 | . | C | T | missense | p.Arg259Trp | 35 | 1.45E-05 |
| 6 | 401454 | rs781276700 | G | A | missense | p.Arg259Gln | 27.1 | 4.07E-06 |
| 6 | 401492 | rs752393433 | G | A | missense | p.Glu272Lys | 33 | 4.07E-06 |
| 6 | 401501 | rs771669183 | C | T | missense | p.Arg275Trp | 32 | 3.23E-05 |
| 6 | 401502 | rs139858073 | G | A | missense | p.Arg275Gln | 27.6 | 2.53E-05 |
| 6 | 401509 | . | CCATGGA | C | indel-inframe | p.Gly279_His280del | 20.3 | private |
| 6 | 401510 | rs765634575 | C | T | missense | p.His278Tyr | 22 | 4.06E-06 |
| 6 | 401520 | rs751008894 | C | T | missense | p.Thr281Met | 22.8 | 1.63E-05 |
| 6 | 401527 | rs114916515 | C | G | missense | p.Asp283Glu | 0.003 | 0.000249 |
| 6 | 401528 | rs767774669 | G | A | missense | p.Ala284Thr | 14.13 | 5.69E-05 |
| 6 | 401528 | rs767774669 | G | T | missense | p.Ala284Ser | 0.642 | 4.06E-06 |
| 6 | 401529 | . | C | T | missense | p.Ala284Val | 21.3 | 6.46E-05 |
| 6 | 401542 | . | C | G | missense | p.Asp288Glu | 12.06 | 4.06E-06 |
| 6 | 401545 | . | G | C | missense | p.Gln289His | 23 | 3.23E-05 |
| 6 | 401571 | rs140099868 | A | G | missense | p.Asn298Ser | 0.434 | 0.000148 |
| 6 | 401589 | rs756281792 | T | C | missense | p.Ile304Thr | 0.256 | 1.44E-05 |
| 6 | 401621 | rs780902340 | G | A | missense | p.Val315Met | 28.2 | 8.14E-06 |
| 6 | 401627 | rs745470687 | C | A | missense | p.Leu317Ile | 29.4 | 1.63E-05 |
| 6 | 401637 | rs775326241 | C | A | missense | p.Ala320Asp | 25.2 | 3.23E-05 |
| 6 | 401637 | rs775326241 | C | G | missense | p.Ala320Gly | 17.79 | 4.07E-06 |
| 6 | 401642 | . | G | A | missense | p.Asp322Asn | 26.4 | 4.07E-06 |
| 6 | 401652 | rs773521348 | A | G | missense | p.Tyr325Cys | 24.3 | 4.08E-06 |
| 6 | 401655 | rs201105575 | C | T | missense | p.Ala326Val | 23.2 | 6.88E-05 |
| 6 | 401690 | . | G | A | missense | p.Gly338Arg | 24.4 | 4.09E-06 |
| 6 | 401700 | rs200311468 | C | T | missense | p.Ala341Val | 25.8 | 0.000342 |
| 6 | 401708 | rs756088477 | A | C | missense | p.Asn344His | 16.94 | 4.10E-06 |
| 6 | 401711 | rs753680565 | G | A | missense | p.Asp345Asn | 26.4 | 2.18E-05 |
| 6 | 401714 | . | C | G | missense | p.Arg346Gly | 24.8 | 4.10E-06 |
| 6 | 401714 | . | C | T | missense | p.Arg346Trp | 27 | 4.10E-06 |
| 6 | 401715 | rs199596593 | G | A | missense | p.Arg346Gln | 25.9 | 2.91E-05 |
| 6 | 401720 | . | A | C | missense | p.Asn348His | 25.5 | 4.10E-06 |
| 6 | 401737 | rs199919569 | C | G | missense | p.Asp353Glu | 8.794 | 4.10E-06 |
| 6 | 401748 | rs555752097 | A | G | missense | p.Lys357Arg | 28 | 4.10E-06 |
| 6 | 401763 | rs779841481 | A | C | missense | p.Gln362Pro | 22.4 | 1.09E-05 |
| 6 | 401775 | . | C | T | missense | p.Ser366Leu | 22.2 | 3.23E-05 |
| 6 | 405027 | rs568315642 | C | T | missense | p.Ala370Val | 15.04 | 0.000451 |
| 6 | 405027 | rs568315642 | C | A | missense | p.Ala370Glu | 9.242 | 3.23E-05 |
| 6 | 405037 | rs775948276 | C | G | missense | p.His373Gln | 13.91 | 4.06E-06 |
| 6 | 405041 | . | G | A | missense | p.Gly375Ser | 22.3 | 3.23E-05 |
| 6 | 405044 | rs768893830 | C | T | missense | p.Arg376Cys | 33 | 2.03E-05 |
| 6 | 405045 | rs377483798 | G | A | missense | p.Arg376His | 28.6 | 4.88E-05 |
| 6 | 405045 | rs377483798 | G | C | missense | p.Arg376Pro | 28.3 | 7.22E-06 |
| 6 | 405045 | rs377483798 | G | T | missense | p.Arg376Leu | 27.5 | 4.06E-06 |
| 6 | 405051 | . | T | A | missense | p.Leu378Gln | 23.8 | 4.06E-06 |
| 6 | 405053 | . | C | A | missense | p.Pro379Thr | 29.49 | 4.06E-06 |
| 6 | 405063 | . | A | G | missense | p.Gln382Arg | 21.3 | 3.23E-05 |
| 6 | 405064 | rs750322004 | G | C | missense | p.Gln382His | 26 | 4.06E-06 |
| 6 | 405069 | rs142143231 | C | T | missense | p.Thr384Ile | 19.71 | 4.06E-06 |
| 6 | 405069 | rs142143231 | C | T | missense | p.Thr384Ile | 19.71 | 8.12E-06 |
| 6 | 405088 | rs751335149 | G | C | missense | p.Glu390Asp | 27.8 | 1.44E-05 |
| 6 | 405098 | rs201428294 | C | A | missense | p.Pro394Thr | 25 | 4.06E-06 |
| 6 | 405098 | rs201428294 | C | G | missense | p.Pro394Ala | 17.89 | 2.53E-05 |
| 6 | 405130 | rs141640145 | C | A | missense | p.His404Gln | 7.944 | 4.07E-06 |
| 6 | 406770 | . | T | C | missense | p.Leu406Pro | 5.648 | 6.46E-05 |
| 6 | 406772 | . | A | T | missense | p.Arg407Trp | 2.517 | 1.14E-05 |
| 6 | 407474 | rs199763142 | G | A | missense | p.Arg411Lys | 23.2 | 9.29E-05 |
| 6 | 407514 | rs760504688 | C | A | missense | p.Phe424Leu | 22.9 | 8.14E-06 |
| 6 | 407515 | rs763733876 | C | G | missense | p.Leu425Val | 22.2 | 4.07E-06 |
| 6 | 407522 | . | G | C | missense | p.Gly427Ala | 14.31 | 4.07E-06 |
| 6 | 407527 | . | G | A | missense | p.Asp429Asn | 28.5 | 7.23E-06 |
| 6 | 407528 | . | A | G | missense | p.Asp429Gly | 25.6 | 3.23E-05 |
| 6 | 407542 | rs756736434 | A | T | missense | p.Ile434Phe | 1.929 | 9.69E-05 |
| 6 | 407543 | rs766230745 | TCAG | T | inframe del | p.Ser435del | 12.91 | 7.23E-06 |
| 6 | 407560 | . | T | A | missense | p.Tyr440Asn | 29.1 | 4.09E-06 |
| 6 | 407561 | . | A | T | missense | p.Tyr440Phe | 16.9 | 4.10E-06 |
| 6 | 407564 | rs752069005 | A | T | missense | p.His441Leu | 22.8 | 1.23E-05 |
| 6 | 407566 | . | A | G | missense | p.Arg442Gly | 26.9 | private |
| 6 | 407572 | rs559734928 | A | G | missense | p.Ile444Val | 9.725 | 1.25E-05 |
| 6 | 407575 | rs748516736 | C | T | missense | p.Arg445Cys | 35 | 2.10E-05 |
| 6 | 407576 | rs756602021 | G | A | missense | p.Arg445His | 23.8 | 2.11E-05 |
| 6 | 407576 | rs756602021 | G | T | missense | p.Arg445Leu | 23.8 | 4.23E-06 |
| 6 | 407578 | . | C | T | missense | p.His446Tyr | 29.2 | 4.22E-06 |
| 6 | 407582 | rs749474191 | C | T | missense | p.Ser447Phe | 33 | 4.25E-06 |
| 6 | 407585 | . | C | T | missense | p.Ser448Phe | 32 | 4.27E-06 |

**Supplementary Table S4A.**
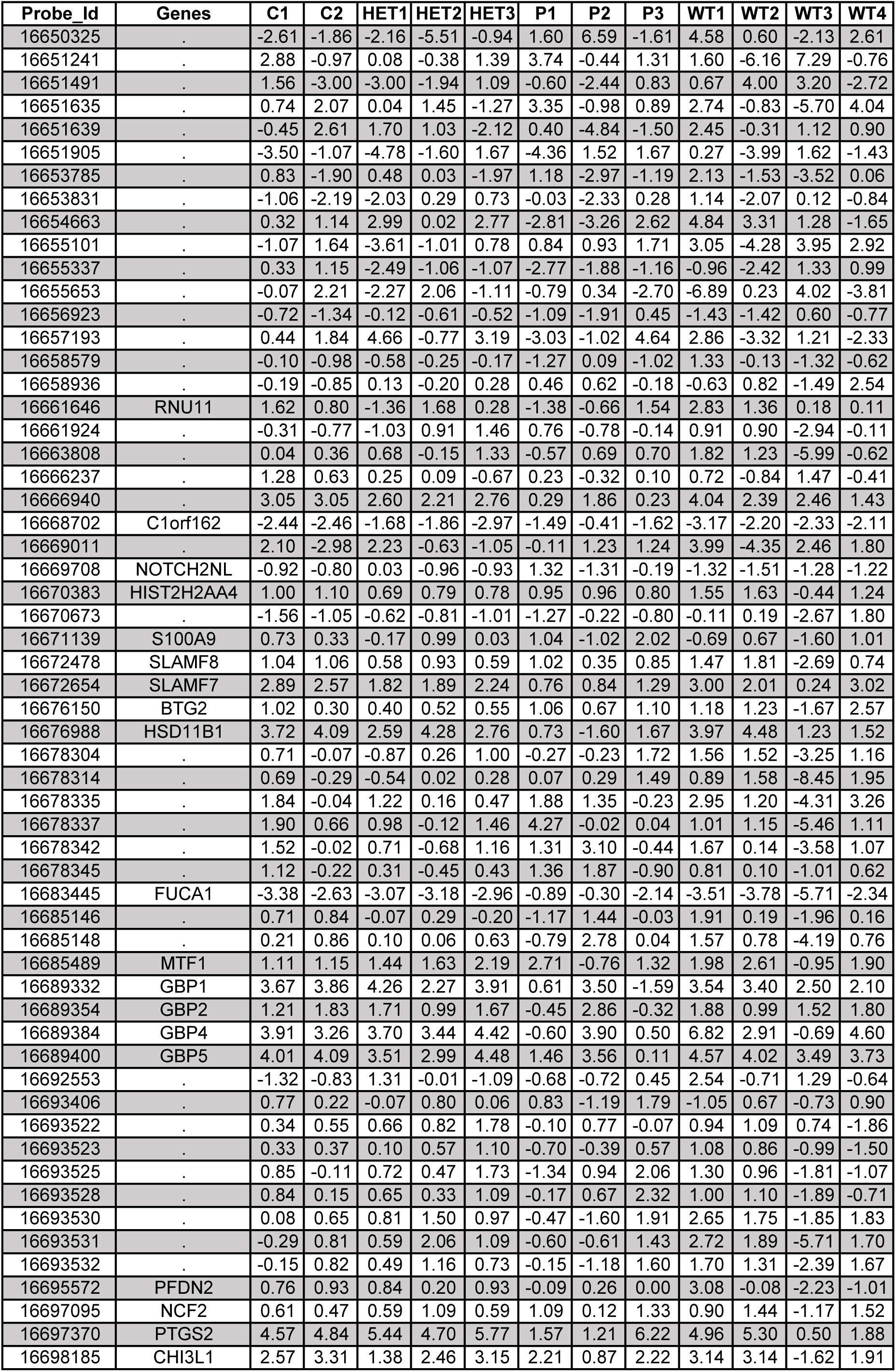
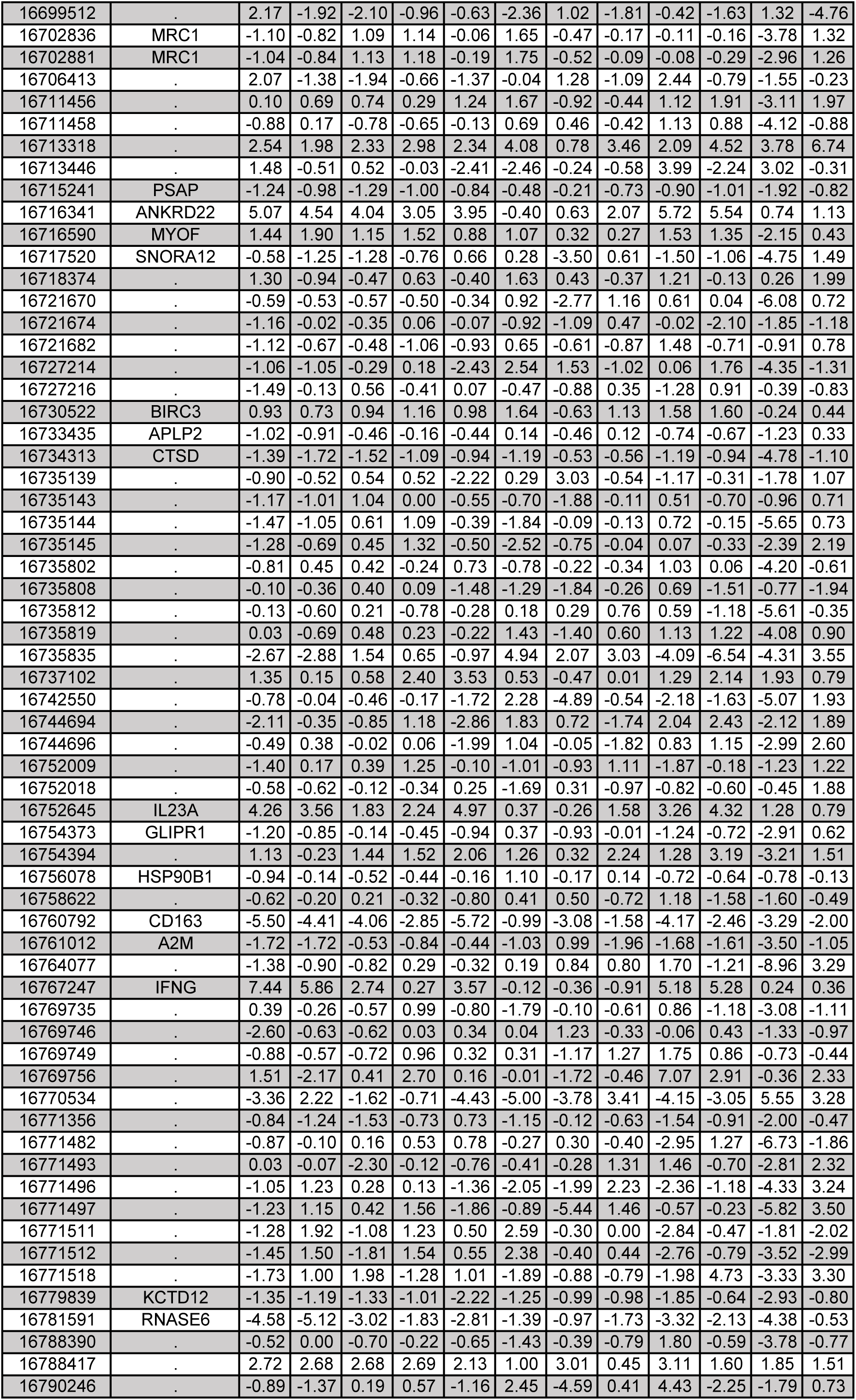
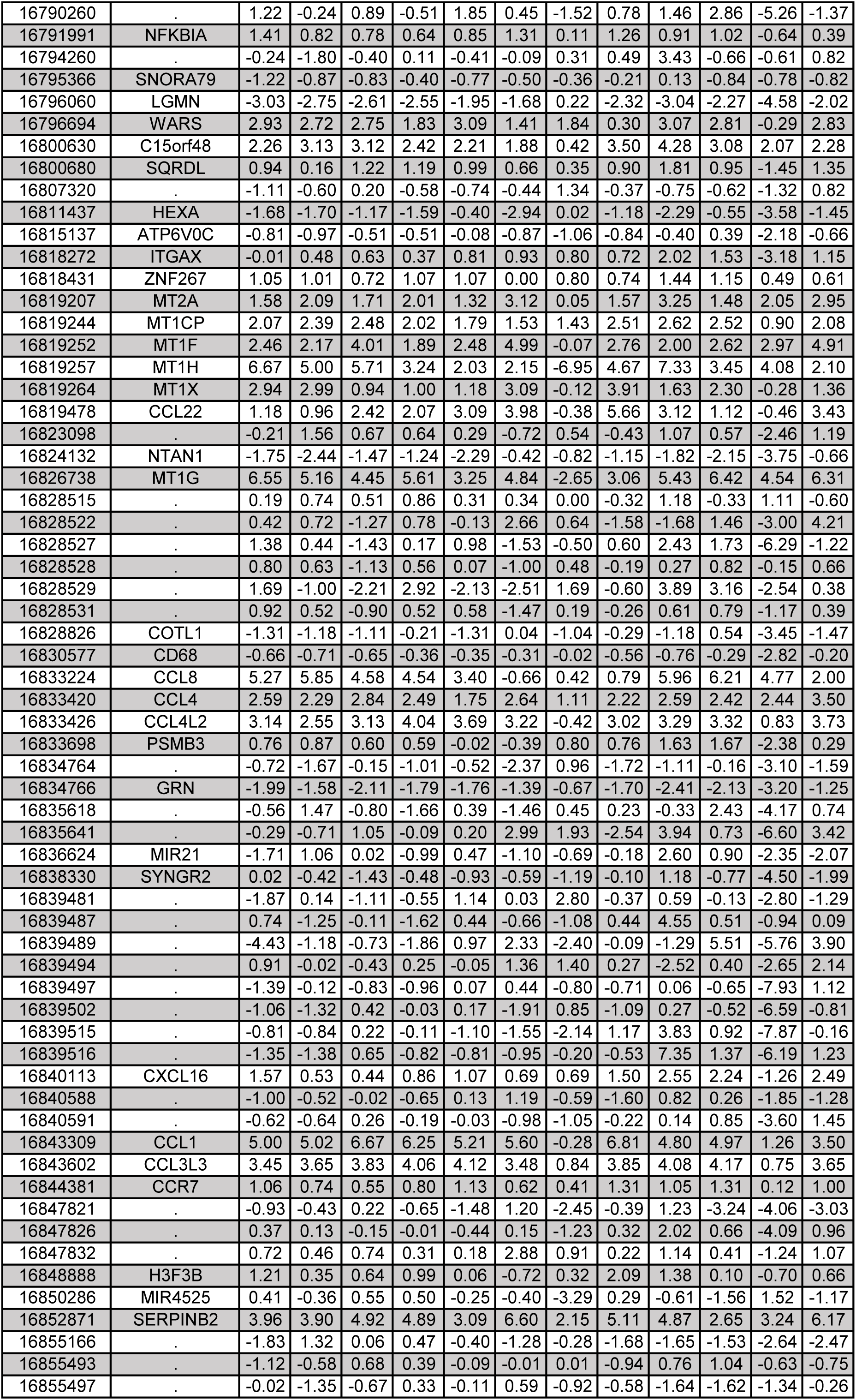
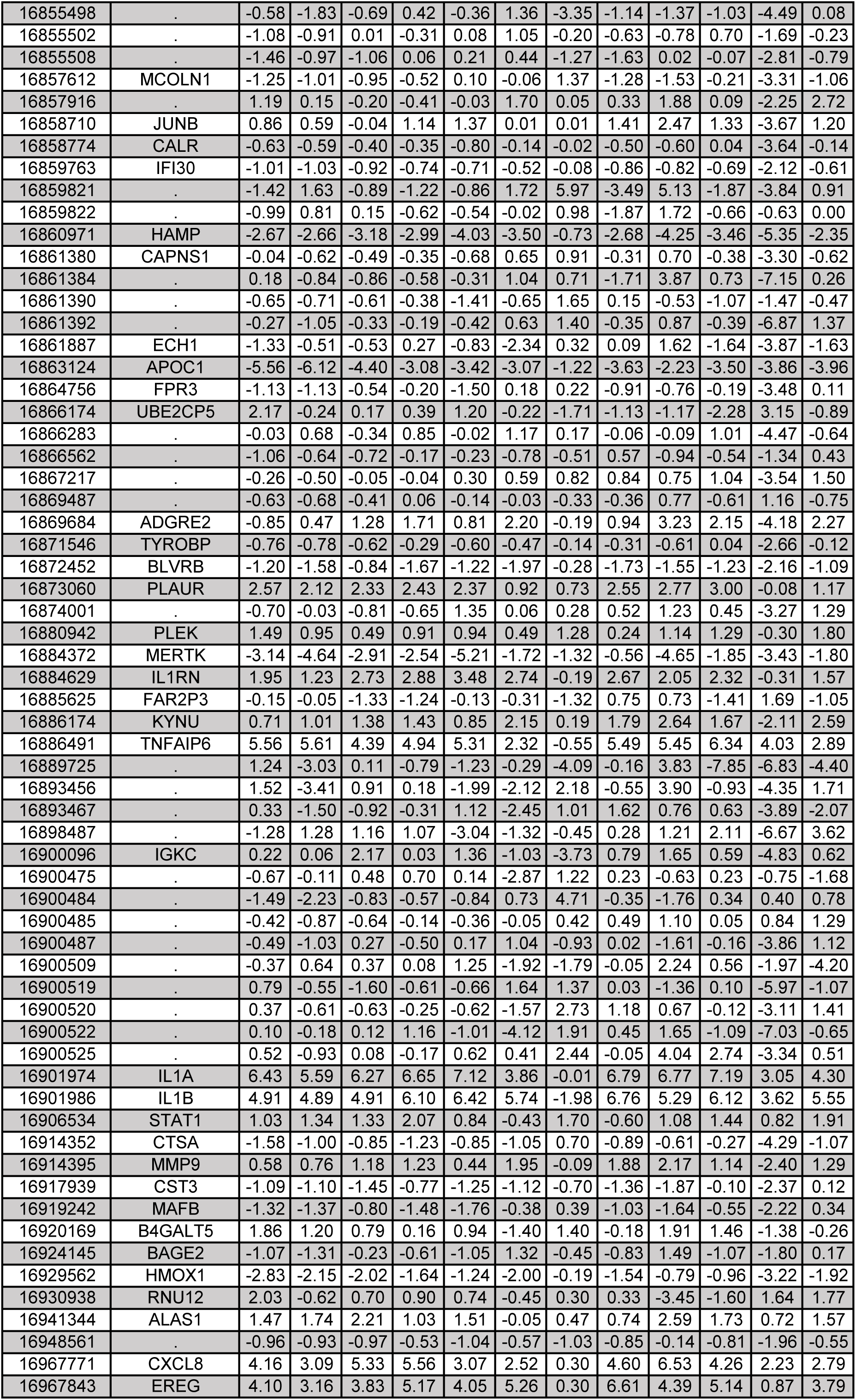
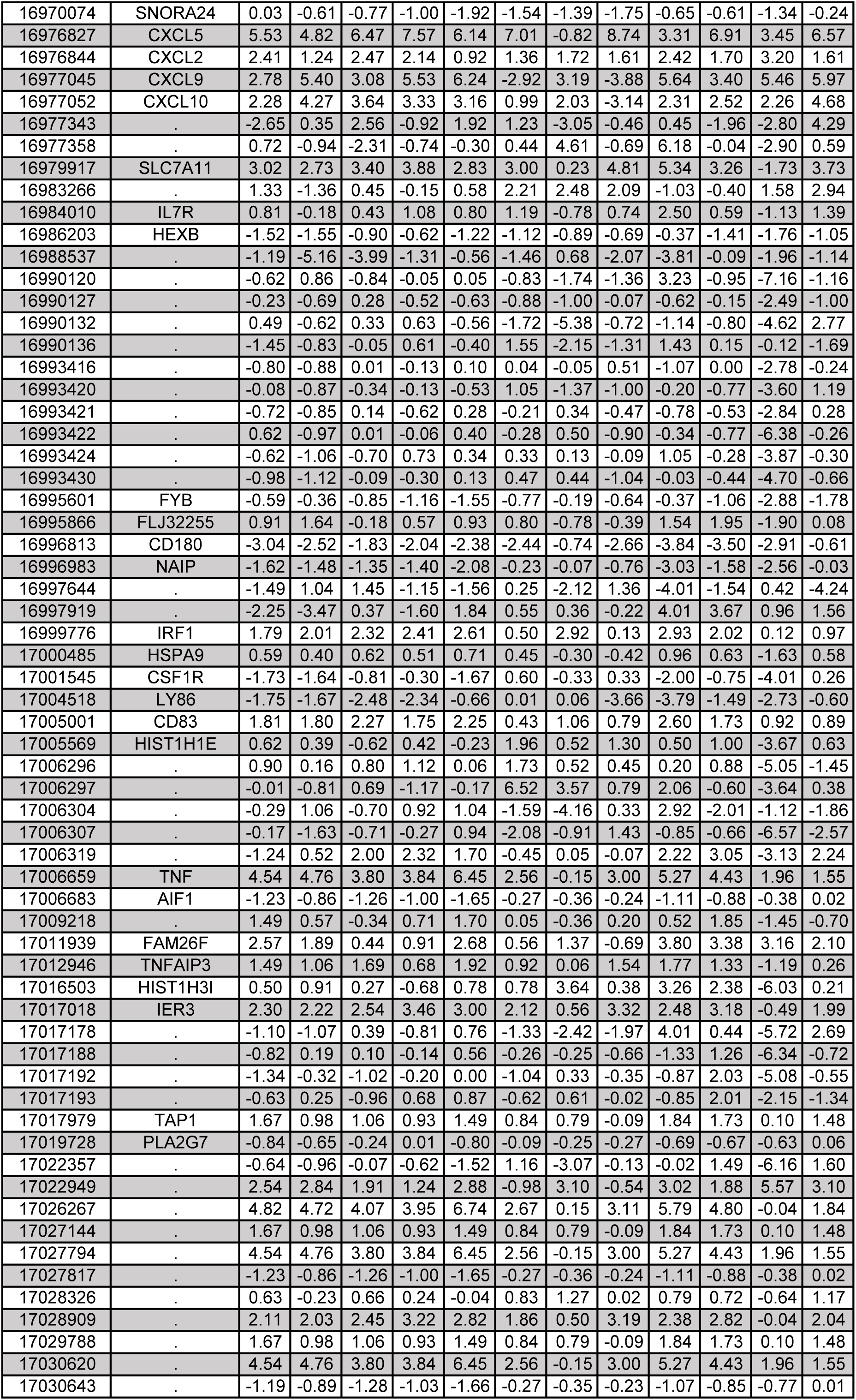
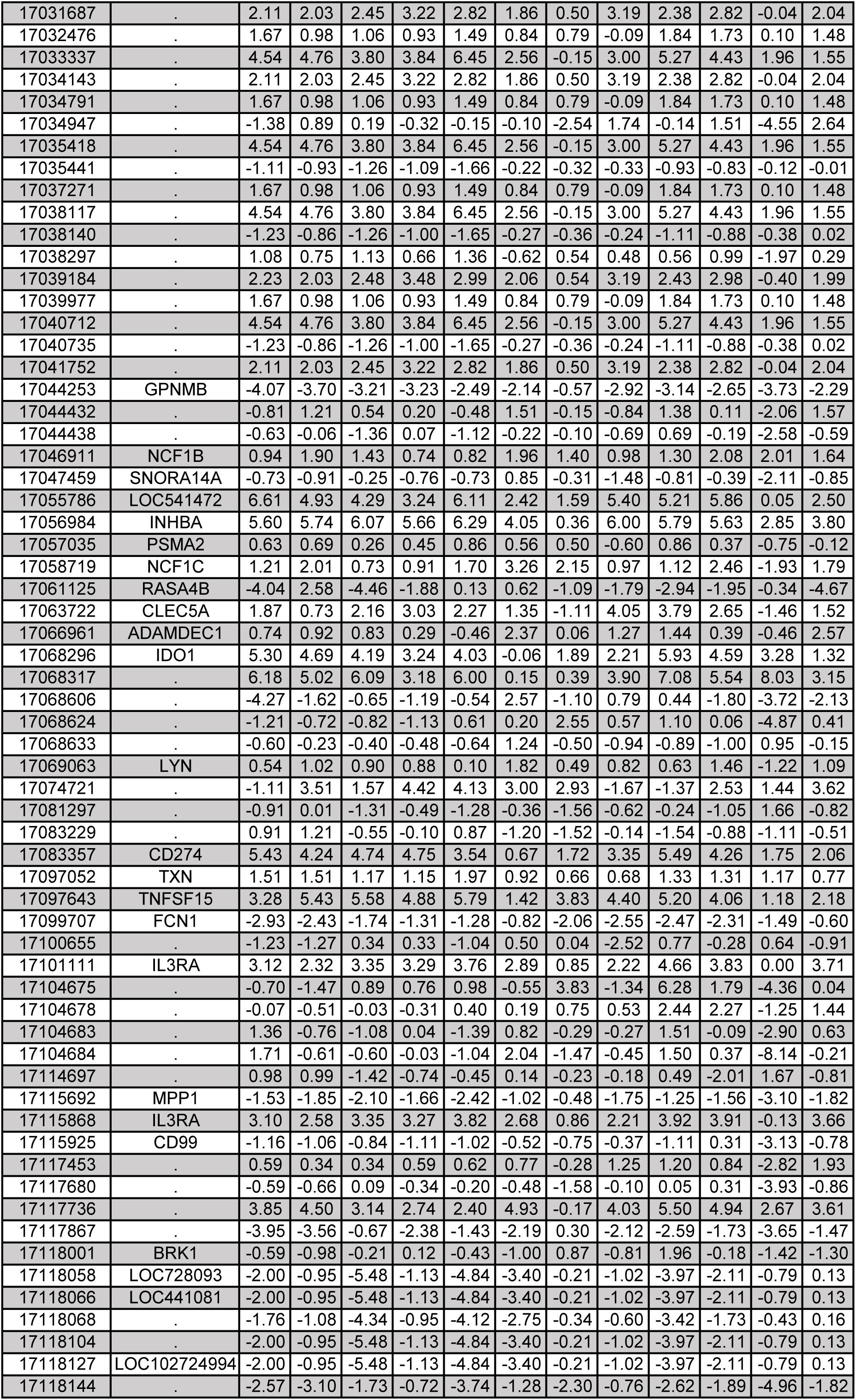
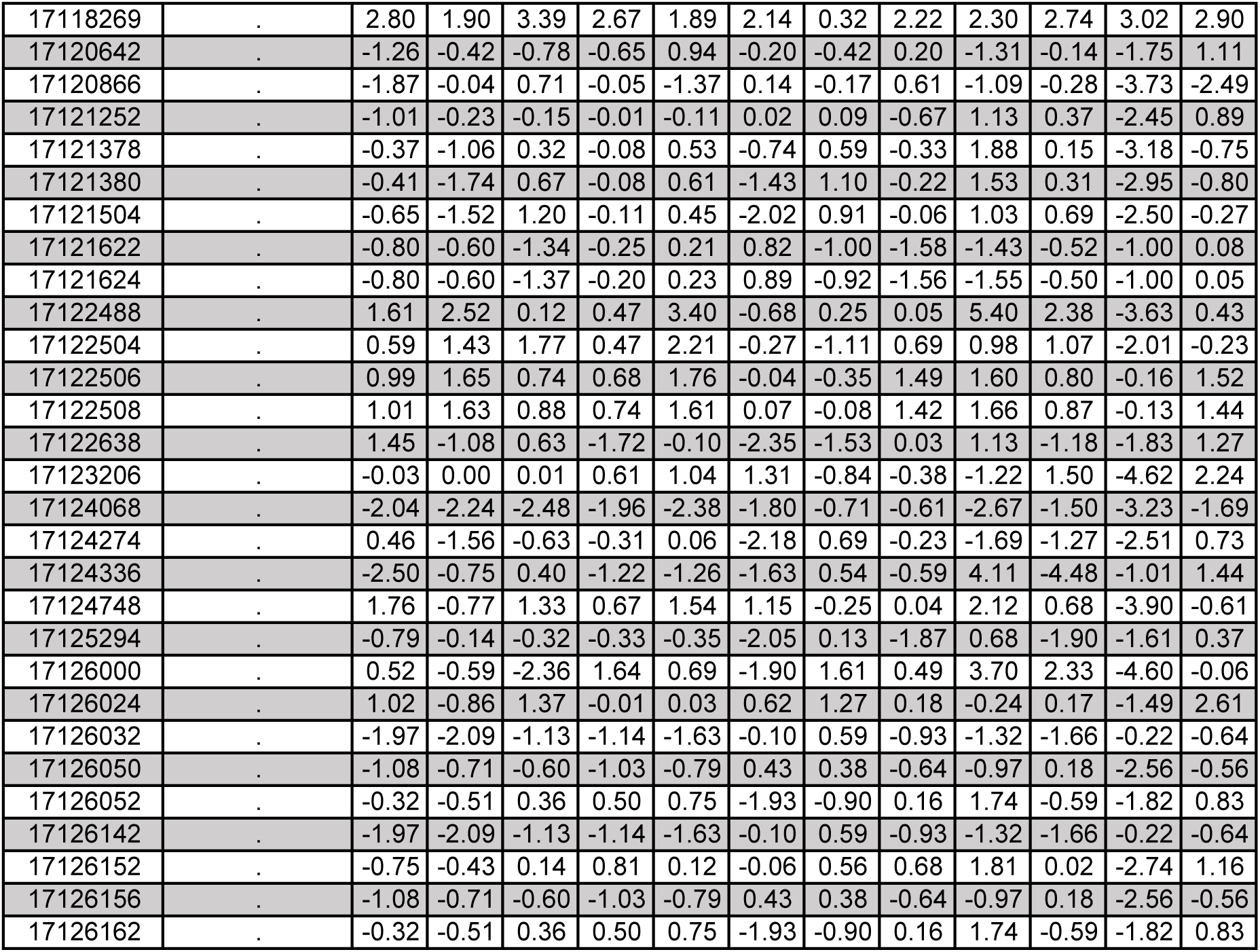
Differentially expressed genes based on transcriptomic data for PBMCs exposed to BCG.

**Supplementary Table S4B.** Differentially expressed genes based on transcriptomic data for PBMCs exposed to Tw.

| Probe_Id | Genes | C1 | C2 | HET1 | HET2 | HET3 | P1 | P2 | P3 | WT1 | WT2 | WT3 | WT4 |
| --- | --- | --- | --- | --- | --- | --- | --- | --- | --- | --- | --- | --- | --- |
| 16650647 | . | -1.33 | -0.08 | 0.55 | -0.11 | -0.02 | -2.39 | -0.06 | -1.93 | -3.58 | 1.55 | -1.81 | 0.97 |
| 16651905 | . | -3.30 | -1.32 | -4.44 | -3.02 | -0.38 | -4.24 | 2.57 | -1.24 | 2.04 | -4.76 | 0.66 | -0.25 |
| 16652101 | . | 3.55 | 0.82 | 1.69 | 0.39 | 0.57 | 1.08 | 1.50 | 4.45 | 4.90 | -7.55 | 2.40 | -2.41 |
| 16653707 | . | 1.66 | -2.88 | 1.37 | 0.85 | 2.59 | 0.10 | 1.68 | 4.39 | 1.30 | -4.11 | 4.95 | -4.14 |
| 16653785 | . | 0.70 | -1.66 | 1.17 | 1.36 | 0.46 | -0.94 | -1.24 | 0.06 | 1.01 | 0.99 | -1.47 | 0.50 |
| 16653795 | . | -0.84 | 0.53 | -0.75 | 0.54 | 0.60 | 1.73 | 0.50 | 0.93 | -1.21 | 1.45 | 0.25 | -1.17 |
| 16654693 | . | -3.80 | -0.07 | 1.15 | 2.22 | -2.32 | -0.34 | -1.92 | -0.22 | 3.08 | -1.70 | -0.15 | -0.72 |
| 16657037 | . | 0.01 | 1.67 | 1.22 | -0.76 | 0.72 | -7.52 | 2.28 | -0.23 | 6.56 | 1.54 | -2.14 | 1.97 |
| 16658933 | . | 0.23 | -1.09 | -0.49 | 0.86 | 0.39 | -0.36 | -1.57 | 0.83 | 0.65 | 2.11 | 0.62 | 2.68 |
| 16658936 | . | 0.74 | -0.63 | 0.17 | 0.44 | 0.02 | 2.56 | 0.29 | -0.22 | 0.10 | -0.39 | 0.94 | 1.50 |
| 16663808 | . | -1.39 | -0.36 | 0.24 | -0.32 | 0.93 | 1.17 | 0.26 | -0.38 | 1.51 | 1.00 | 0.04 | -0.61 |
| 16678342 | . | -0.62 | -0.08 | 0.17 | -1.41 | 1.25 | -2.67 | 2.38 | -0.03 | 1.12 | -0.22 | -0.89 | -1.08 |
| 16693644 | . | -0.59 | -0.96 | -0.24 | -1.15 | -0.34 | 1.08 | 1.30 | -0.37 | 2.18 | 0.68 | -1.63 | 0.10 |
| 16699512 | . | 2.81 | -4.16 | -0.78 | -0.25 | 3.93 | -1.86 | 0.65 | 2.14 | -0.19 | -3.01 | 0.87 | -1.12 |
| 16718373 | . | 1.11 | -0.90 | -0.71 | 0.97 | -0.29 | -0.28 | 0.09 | 1.08 | 2.59 | 1.38 | 0.12 | 1.89 |
| 16732028 | LOC105369519 | -0.87 | -0.95 | -0.05 | -0.08 | 0.34 | -2.11 | 0.07 | 1.72 | 1.87 | 0.34 | 0.77 | -2.22 |
| 16735142 | . | 1.96 | 1.67 | 1.19 | 0.09 | -0.91 | -0.32 | 0.84 | 2.57 | 3.55 | 1.15 | -2.65 | -1.28 |
| 16735819 | . | -0.74 | -0.89 | -0.26 | 0.16 | 0.12 | 0.56 | -0.32 | 0.72 | 0.73 | 1.13 | -0.18 | 1.12 |
| 16735835 | . | 0.42 | 0.80 | 0.41 | 2.54 | -2.38 | -0.73 | 4.84 | 4.26 | -3.65 | -1.97 | 0.38 | 4.69 |
| 16740560 | . | -0.65 | -0.50 | 0.36 | 0.36 | 0.42 | 0.18 | -0.56 | 0.04 | 0.64 | 0.80 | 0.74 | 0.34 |
| 16745154 | . | 0.82 | -0.82 | -0.53 | 0.47 | 0.09 | 0.07 | 0.02 | -0.29 | 1.86 | -0.23 | -0.56 | 3.40 |
| 16747336 | SCARNA10 | 0.89 | 0.29 | -0.70 | 0.02 | 0.37 | -0.20 | 1.30 | 0.51 | 0.84 | -1.44 | 0.04 | -0.73 |
| 16751995 | . | 1.36 | -2.10 | 2.56 | -4.08 | 4.43 | -1.29 | 0.09 | -0.67 | -1.04 | 1.41 | -1.30 | -0.67 |
| 16752007 | . | -1.31 | 0.81 | 2.90 | 0.16 | 0.06 | 1.62 | 0.99 | 1.68 | 5.58 | -0.33 | 1.62 | 0.74 |
| 16755523 | . | 0.31 | -0.83 | 0.42 | -0.15 | 0.01 | 1.03 | -0.72 | 0.45 | 1.96 | -0.21 | -0.67 | 1.73 |
| 16764077 | . | -1.27 | -0.56 | 1.04 | 0.92 | 0.06 | -3.16 | -0.61 | 0.90 | 1.92 | -0.31 | -2.67 | 3.99 |
| 16769725 | . | -1.53 | 1.81 | -0.92 | 1.44 | 0.83 | 0.60 | -4.09 | -0.76 | -1.59 | -2.26 | 0.22 | -4.09 |
| 16769727 | . | -1.29 | -0.47 | 0.85 | 0.31 | -0.81 | 1.14 | -1.04 | 1.92 | -0.76 | -1.15 | -0.82 | -2.14 |
| 16769728 | . | -0.82 | -0.86 | -0.88 | 1.73 | -0.24 | -1.92 | 0.43 | 1.42 | -1.79 | 0.98 | -2.09 | 5.19 |
| 16769749 | . | -0.08 | -1.03 | -0.02 | 0.08 | 0.05 | -1.87 | 0.24 | 1.35 | 2.11 | -0.24 | -0.97 | -0.98 |
| 16769758 | . | -1.16 | -0.62 | 2.16 | 0.87 | 2.15 | -3.75 | -5.68 | -0.53 | 4.47 | 1.98 | -1.91 | 1.08 |
| 16771317 | . | -0.17 | 0.94 | -0.85 | 1.09 | 0.82 | 0.77 | 0.40 | 1.40 | 7.33 | 1.29 | -1.83 | -0.35 |
| 16771492 | . | -2.11 | -1.93 | 2.03 | -0.28 | 2.37 | -1.87 | -1.29 | 0.65 | -1.90 | 2.48 | -0.82 | 1.43 |
| 16771510 | . | 0.12 | 1.16 | -1.12 | 0.11 | 0.48 | 0.36 | 1.84 | 0.36 | 1.62 | 0.60 | 0.58 | -1.99 |
| 16771559 | . | -0.83 | -1.76 | -1.80 | -1.22 | 1.40 | -1.97 | -1.07 | 0.36 | 5.80 | 0.65 | -1.56 | 2.13 |
| 16790473 | . | 0.05 | 0.18 | -0.41 | -0.89 | 0.02 | -0.02 | 0.34 | -0.29 | 1.50 | 0.89 | -0.60 | -0.70 |
| 16794260 | . | -0.73 | 0.02 | 0.44 | -0.29 | 0.13 | 2.22 | -1.34 | 0.99 | 3.21 | -0.75 | -0.05 | -1.39 |
| 16823043 | SNHG19 | 1.06 | 1.10 | 2.45 | 1.09 | -0.59 | -2.24 | 1.63 | -1.37 | -0.35 | 0.53 | -1.93 | -3.87 |
| 16823105 | . | -0.81 | -0.88 | 0.29 | 0.15 | -0.40 | -0.17 | 0.54 | 1.89 | -0.49 | 0.85 | 1.71 | 1.00 |
| 16828270 | . | -0.94 | 0.26 | 1.43 | 0.43 | 1.08 | -0.01 | -0.82 | 0.74 | 0.60 | -0.28 | 1.66 | 0.61 |
| 16828528 | . | 0.89 | 0.85 | -0.50 | 0.61 | -0.03 | 1.40 | 1.07 | 0.45 | -0.20 | -0.31 | -0.95 | 0.94 |
| 16828531 | . | 0.86 | 0.65 | -0.71 | 0.43 | 0.51 | 0.36 | 0.49 | 0.50 | 0.45 | -0.40 | -1.07 | 1.10 |
| 16835618 | . | -0.06 | -1.07 | 1.66 | -1.36 | 1.43 | 0.22 | -0.67 | 0.93 | 5.53 | 1.56 | 1.60 | -2.34 |
| 16835641 | . | 0.27 | 0.80 | -0.32 | -1.26 | -0.29 | 3.90 | 2.49 | -0.36 | 4.45 | 1.03 | -0.97 | 3.40 |
| 16836624 | MIR21 | -1.84 | -1.38 | 1.11 | -0.14 | 0.92 | -1.59 | -0.70 | -0.23 | 2.26 | -0.18 | 1.11 | -2.35 |
| 16839474 | . | -0.41 | 0.72 | 0.63 | 0.24 | -0.24 | -1.83 | 1.61 | -0.53 | 0.80 | -0.87 | -0.74 | 1.92 |
| 16839477 | . | -1.60 | -1.12 | 0.87 | 1.14 | 0.32 | -1.85 | 0.93 | 0.76 | -3.53 | -1.81 | -0.94 | -1.26 |
| 16839481 | . | -0.87 | 0.53 | 0.79 | -0.25 | 0.90 | -3.50 | 0.23 | 1.03 | -1.21 | 0.61 | 1.17 | -1.64 |
| 16839489 | . | -3.93 | 2.15 | 0.79 | -0.73 | 0.85 | -0.54 | -0.75 | 0.31 | 0.28 | 3.46 | 0.79 | 6.30 |
| 16839492 | . | -1.45 | 0.68 | 0.23 | -0.63 | -0.34 | 3.09 | -0.36 | 0.85 | -1.72 | 0.45 | 0.73 | 4.71 |
| 16839494 | . | 1.34 | -0.72 | -0.16 | 0.64 | 0.53 | 2.70 | 0.20 | -0.05 | -2.29 | 0.64 | -1.14 | 1.79 |
| 16839506 | . | -2.63 | -1.15 | -0.16 | 0.23 | -0.17 | -2.67 | 0.88 | 0.39 | 1.14 | 1.61 | 1.37 | -1.63 |
| 16839513 | . | -0.66 | -0.69 | -0.33 | -0.70 | 0.46 | -0.83 | -0.31 | 0.10 | 2.50 | 0.63 | 0.21 | -0.71 |
| 16839515 | . | -0.84 | -1.14 | 1.71 | 0.05 | -0.53 | -0.92 | -0.26 | -0.16 | 3.09 | 0.91 | -1.13 | 0.19 |
| 16840589 | . | -2.62 | 0.60 | 0.96 | 0.31 | 0.87 | 0.50 | -1.11 | 0.67 | -0.29 | 1.08 | -0.88 | -2.81 |
| 16842465 | . | -2.74 | 2.06 | 2.24 | -0.96 | -0.86 | -1.72 | 0.23 | 3.34 | -2.64 | -2.61 | 2.33 | -2.14 |
| 16842466 | . | -0.44 | 0.99 | 1.53 | -1.25 | -0.99 | -0.13 | 0.17 | 1.20 | -2.32 | -2.90 | 1.74 | -2.22 |
| 16847247 | . | 3.41 | -2.06 | 1.77 | 2.75 | 2.02 | -0.87 | 0.89 | 1.19 | 5.20 | -0.68 | 1.46 | -2.23 |
| 16847319 | MIR4737 | -3.30 | -0.02 | 3.15 | -0.26 | 0.29 | -0.89 | 2.03 | -0.72 | 0.68 | -2.86 | -1.57 | -2.04 |
| 16852308 | SCARNA17 | -0.42 | 0.59 | -0.63 | 0.53 | -0.93 | -1.70 | -0.45 | -0.42 | -0.41 | -0.86 | 1.83 | -2.00 |
| 16857916 | . | 0.26 | -0.04 | -1.28 | -0.25 | -0.27 | 1.02 | -0.13 | 0.21 | 1.39 | -1.05 | 0.85 | 2.06 |
| 16859821 | . | -1.04 | -0.09 | -0.63 | 1.02 | 0.05 | -3.23 | 3.29 | 0.39 | 4.46 | -0.84 | 1.14 | -0.27 |
| 16859824 | . | -0.71 | 0.99 | 0.14 | -1.48 | -0.60 | 4.68 | -1.09 | 3.43 | -1.33 | 0.65 | 1.18 | 3.24 |
| 16861887 | ECH1 | 0.62 | -0.24 | -0.28 | 0.45 | -0.62 | -0.61 | 2.38 | -0.23 | 1.02 | -0.50 | -0.87 | -0.67 |
| 16893456 | . | 1.90 | -1.32 | 0.81 | 1.13 | 1.68 | -4.12 | -1.30 | 0.51 | 2.62 | 1.23 | -0.63 | 1.59 |
| 16898487 | . | -1.05 | 0.70 | 3.24 | 2.82 | -0.75 | -2.79 | 2.73 | 0.68 | 0.17 | 1.84 | -4.70 | 1.84 |
| 16900096 | IGKC | -0.59 | -0.81 | -1.13 | -0.18 | 0.98 | 3.37 | -0.41 | 1.39 | 1.21 | 0.79 | -0.17 | 1.06 |
| 16900461 | . | 0.07 | -0.89 | -0.46 | -0.50 | 0.29 | 1.38 | 1.67 | -0.46 | 2.39 | 0.98 | 0.49 | -1.15 |
| 16900505 | . | -1.06 | -0.26 | -1.54 | -0.65 | -0.46 | -1.06 | -0.80 | 0.94 | 1.20 | -0.93 | 1.62 | 0.33 |
| 16900506 | . | -1.45 | -0.92 | -0.19 | -1.42 | 0.86 | -1.22 | 0.41 | 0.58 | 4.77 | 1.10 | 1.50 | 1.77 |
| 16900520 | . | -0.61 | -1.03 | -1.21 | -0.36 | -0.17 | -0.20 | 1.49 | 0.58 | -0.69 | -0.67 | 0.71 | 1.57 |
| 16900530 | . | -1.68 | 1.14 | 0.24 | -0.68 | 2.07 | -2.20 | 0.33 | 0.40 | -0.03 | -0.03 | -0.80 | -2.17 |
| 16915993 | PPDPF | -1.08 | -1.23 | 0.64 | -0.25 | 0.08 | -0.38 | -0.21 | 0.11 | 0.65 | -0.31 | 0.59 | 1.68 |
| 16934240 | . | -1.99 | -1.06 | 1.60 | -0.15 | 0.02 | -0.47 | -0.41 | 0.21 | -0.84 | -1.70 | -2.76 | -0.21 |
| 16934244 | . | -1.55 | -1.79 | 1.61 | -0.59 | 0.14 | -0.64 | -0.66 | 0.42 | -1.39 | -2.41 | -3.01 | -0.05 |
| 16934245 | . | -1.40 | -0.54 | -1.58 | -0.05 | 0.62 | -2.10 | -1.11 | 0.63 | -0.69 | -2.54 | -0.17 | -1.18 |
| 16934595 | . | 1.34 | 1.26 | -0.80 | -1.78 | -0.39 | 1.70 | -0.39 | -0.09 | 4.87 | -5.23 | 1.27 | -7.36 |
| 16934598 | . | 1.00 | -1.06 | 1.23 | -0.11 | 0.02 | 6.14 | -0.57 | -0.80 | 1.58 | -0.67 | -0.25 | 1.50 |
| 16934602 | . | 0.04 | -1.33 | -0.04 | -0.07 | -0.47 | -2.96 | 0.30 | 0.86 | 2.09 | -0.62 | -0.19 | 0.81 |
| 16934603 | . | 0.14 | -0.86 | 1.17 | -0.58 | -0.06 | -2.08 | 0.46 | 0.65 | 1.98 | -0.76 | -1.43 | -0.15 |
| 16961037 | SCARNA7 | -2.22 | 1.02 | -2.26 | 1.35 | -2.27 | 4.58 | -0.47 | -0.69 | 0.78 | 0.22 | -2.31 | -1.37 |
| 16983266 | . | -0.49 | -1.65 | -2.46 | 0.02 | 0.93 | 1.01 | 0.46 | 1.29 | -2.78 | -2.22 | 0.20 | 2.09 |
| 16988537 | . | -0.33 | -1.65 | -3.89 | -1.23 | -1.62 | -1.51 | -0.32 | -1.83 | -1.92 | -3.89 | -1.59 | -2.43 |
| 16989865 | . | -0.14 | -1.53 | 0.27 | -0.33 | 0.25 | -3.85 | 1.00 | 0.48 | -2.07 | -0.96 | 0.37 | 2.26 |
| 16990120 | . | 0.43 | -0.50 | -0.10 | 0.49 | -0.23 | 1.08 | -1.18 | -0.86 | 3.68 | -0.71 | -0.81 | -0.84 |
| 16997644 | . | -2.33 | -0.44 | 1.61 | 0.42 | -1.00 | 1.41 | 1.34 | 0.96 | -2.05 | -1.88 | 0.06 | -2.11 |
| 17006290 | . | 1.47 | 0.06 | 0.63 | 0.72 | 1.58 | -1.62 | 0.80 | 0.22 | 1.19 | -1.40 | 0.10 | -2.06 |
| 17006294 | . | 0.32 | -0.34 | 0.60 | -0.01 | 0.12 | -2.54 | 0.08 | 0.19 | 1.14 | -1.31 | -0.60 | 0.76 |
| 17006295 | . | 0.09 | -1.23 | -0.02 | -0.10 | -0.02 | -0.46 | 0.76 | 0.11 | 1.33 | -1.11 | -1.22 | 0.81 |
| 17006296 | . | 0.77 | -0.84 | 1.34 | 0.08 | -0.22 | -4.68 | -0.40 | -0.41 | 0.61 | -1.76 | -0.16 | -2.33 |
| 17006305 | . | -1.21 | -2.08 | -3.30 | -0.83 | -0.24 | -3.51 | 0.53 | -0.03 | 3.19 | -0.58 | 0.50 | -2.34 |
| 17006306 | . | -1.29 | -2.60 | 1.21 | -1.64 | -1.04 | -1.34 | 1.53 | 0.71 | 6.02 | -3.72 | -0.03 | 0.88 |
| 17006308 | . | -1.59 | -0.44 | -0.42 | -1.35 | 1.21 | -1.30 | 1.34 | 0.23 | 2.20 | -0.74 | 0.88 | -0.17 |
| 17016372 | HIST1H1C | 0.08 | -0.18 | 0.24 | -0.26 | 0.75 | 0.96 | 0.23 | 1.16 | 2.01 | 1.11 | 0.67 | -1.34 |
| 17016503 | HIST1H3I | -1.23 | -0.39 | 0.52 | -0.69 | 0.25 | -2.58 | -2.26 | 0.08 | 2.78 | 1.23 | -0.45 | -2.91 |
| 17017177 | . | 0.46 | -0.77 | -0.49 | -0.64 | 0.78 | -1.45 | -0.08 | 0.09 | 1.33 | 0.98 | 1.09 | 0.10 |
| 17017194 | . | -4.83 | -0.81 | 3.34 | 1.08 | 1.27 | -0.88 | 0.46 | -1.11 | 1.99 | 3.95 | 2.61 | 6.54 |
| 17017197 | . | 0.79 | -0.86 | 1.26 | 0.34 | -0.70 | -2.40 | 0.71 | 0.17 | 5.90 | -1.69 | -0.44 | 4.28 |
| 17018387 | . | -0.72 | -0.79 | 0.75 | 0.45 | 0.43 | -2.84 | 0.54 | 0.18 | 4.74 | 1.47 | -1.47 | 2.83 |
| 17041697 | . | -0.22 | -0.69 | -0.07 | 0.24 | 0.01 | -0.35 | 0.64 | -0.11 | 0.59 | -0.51 | -0.69 | 0.82 |
| 17047459 | SNORA14A | 0.97 | 0.71 | 0.06 | -0.40 | -0.72 | 1.03 | 0.76 | 0.13 | -0.29 | 0.05 | -1.28 | -1.48 |
| 17061125 | RASA4B | -2.13 | 2.69 | -1.90 | -0.58 | 1.01 | 2.04 | -1.45 | -0.85 | -1.78 | 2.23 | 2.23 | -5.88 |
| 17068624 | . | 0.59 | -0.01 | 0.02 | -0.53 | -0.91 | -0.75 | 0.89 | 1.17 | 0.61 | -0.87 | 0.00 | 3.76 |
| 17100655 | . | 1.29 | 0.94 | 0.46 | 0.71 | -1.34 | -0.27 | 0.02 | -0.39 | -0.98 | -0.64 | 0.55 | 0.46 |
| 17100810 | . | -0.09 | -0.69 | -1.52 | -0.15 | 0.19 | 0.05 | 0.30 | -0.34 | 1.35 | -1.05 | 0.92 | 1.48 |
| 17104684 | . | -0.70 | -0.66 | -0.20 | 0.63 | -1.08 | -1.07 | -0.31 | -0.06 | 0.71 | 0.86 | -0.54 | -0.55 |
| 17118001 | BRK1 | 0.56 | -0.61 | -0.70 | 0.01 | 0.02 | -1.19 | 1.82 | -1.01 | 1.99 | -0.98 | 0.33 | -2.36 |
| 17118426 | . | -0.65 | 1.14 | 0.11 | 0.20 | -0.12 | 0.01 | -0.64 | 0.36 | 3.42 | 1.35 | 0.40 | 1.57 |
| 17118850 | . | 0.86 | 1.06 | -0.25 | -0.02 | 0.06 | -0.44 | 0.71 | 0.19 | 3.34 | -0.58 | 0.42 | 2.19 |
| 17120164 | . | 0.05 | 0.63 | -0.22 | 0.18 | 0.40 | -2.58 | 0.12 | -0.27 | 0.18 | -0.77 | -0.95 | -1.29 |
| 17121378 | . | -0.70 | -0.30 | -0.05 | -0.18 | 0.43 | -0.41 | 0.42 | -0.10 | 1.79 | -0.27 | 0.62 | -0.61 |
| 17121380 | . | -0.66 | -1.08 | 0.36 | -0.30 | 0.40 | -0.38 | 0.02 | -0.04 | 1.64 | -0.19 | 0.93 | -1.22 |
| 17121504 | . | -0.40 | -2.05 | 0.86 | -0.31 | -0.05 | -0.22 | -0.60 | 0.06 | 1.55 | 0.22 | 1.46 | -2.10 |
| 17124336 | . | -0.62 | -0.21 | -0.57 | 0.10 | -0.90 | 0.52 | 0.81 | 0.41 | 3.06 | -1.17 | 0.92 | 1.55 |
| 17124536 | . | 2.59 | -1.58 | 0.92 | 0.65 | -0.69 | -5.38 | 0.34 | 2.85 | 5.23 | -2.26 | -1.19 | 2.99 |
| 17124748 | . | 1.16 | 0.35 | 0.64 | 0.43 | 0.80 | 1.94 | 1.66 | -0.33 | 5.98 | 1.98 | 0.27 | -1.11 |
| 17125856 | . | -1.10 | -2.39 | -1.27 | -1.52 | 1.35 | -1.96 | -1.24 | 0.15 | 2.78 | -1.58 | -1.19 | 1.53 |
| 17126000 | . | -1.43 | -1.42 | -2.62 | 0.31 | 0.00 | -1.15 | 1.71 | -0.91 | 3.36 | 1.17 | -2.27 | 0.09 |
| 17126152 | . | -0.37 | -0.66 | 0.58 | 0.14 | -0.41 | 0.29 | -0.28 | 1.23 | 1.28 | -0.14 | 0.79 | 1.17 |

**Supplementary Table S5.** Immunophenotyping of patients (P1, P2 and P3) and a WT homozygous relative.

|  | P1 | P2 | P3 | P1'sister | Normal range |
| --- | --- | --- | --- | --- | --- |
| Phenotype | WD | WD | WD | Healthy |  |
| IRF4 genotype | R98W/WT | R98W/WT | R98W/WT | WT/WT |  |
| lymphocytes number/μl | 1100 | 2400 | 1200 | 2400 | |
| T Lymphocyte Immunophenotyping (%) |  |  |  |  |  |
| CD3+ | 71 | 77 | 58 | 77 | 51-82 |
| CD4+ | 49 | 42 | 33 | 55 | 31-63 |
| CD8+ | 21 | 30 | 25 | 19 | 9-34 |
| CD45RO+/CD4+ | 48 | 81 | 85 | 40 |  |
| CD45RA+/CD4+ | 52 | 19 | 15 | 60 | 20-86 |
| CD31+/CD4+ | 31 | 17 | 8 | 25 |  |
| CD31+/CD45RA+/CD4+ | 24 | 6 | 3 | 20 | 20-28 |
| CCR7+CD45RA+/CD8+ | 20 | 7 | 6 | 15 | 29-57 |
| CCR7+CD45RA-/CD8+ | 17 | 7 | 12 | 2 | 3-14 |
| CCR7-CD45RA-/CD8+ | 39 | 49 | 27 | 44 | 20-41 |
| CCR7-CD45RA+/CD8+ | 24 | 37 | 55 | 40 | 11-26 |
| B Lymphocyte Immunophenotyping (%) |  |  |  |  |  |
| CD19+ | 8 | 11 | 6 | 9 | 4-21 |
| NK Lymphocyte Immunophenotyping (%) |  |  |  |  |  |
| CD16+CD56+ | 21 | 12 | 36 | 12 | 5-33 |

